# Unveiling the evolutionary history of European vipers and their venoms from a multi-omic approach

**DOI:** 10.1101/2024.12.10.627732

**Authors:** Adrián Talavera, Marc Palmada-Flores, Fernando Martínez-Freiría, Gabriel Mochales-Riaño, Bernat Burriel-Carranza, Maria Estarellas, Daniel Fernández-Guiberteau, Álvaro Camina, Sylvain Ursenbacher, Judit Vörös, Bálint Halpern, Davinia Pla, Juan José Calvete, Alexander S. Mikheyev, Tomàs Marquès-Bonet, Salvador Carranza

**Affiliations:** Institute of Evolutionary Biology (CSIC-Universitat Pompeu Fabra), Barcelona, Spain; CIBIO, Centro de Investigação em Biodiversidade e Recursos Genéticos, InBIO Laboratório Associado, Universidade do Porto, Vairão, Portugal; BIOPOLIS Program in Genomics, Biodiversity and Land Planning, CIBIO, Campus de Vairão, 4485-661 Vairão, Portugal; Museu de Ciències Naturals de Barcelona, P° Picasso s/n, Parc Ciutadella, 08003 Barcelona, Spain; Centre de Recerca i Educació Ambiental de Calafell (CREAC-GRENP-Ajuntament de Calafell), Tarragona, Spain; Grupo Atrox, Faunia, Av. de las Comunidades 28, 28032, Madrid; Section of Conservation Biology, Department of Environmental Sciences, University of Basel, St. Johanns-Vorstadt 10, 4056 Basel, Switzerland; info fauna – CSCF and karch, University of Neuchâtel, Avenue Bellevaux 51, 2000 Neuchâtel, Switzerland; HUN-REN Balaton Limnological Research Institute, 8237 Tihany, Klebelsberg Kuno utca 3., Hungary; MME Birdlife Hungary, Budapest, Hungary; Department of Systematic Zoology and Ecology, Institute of Biology, ELTE-Eötvös Loránd University, Budapest, Hungary; HUN-REN – ELTE - MTM Integrative Ecology Research Group, Budapest, Hungary; Evolutionary and Translational Venomics Laboratory, Consejo Superior de Investigaciones Científicas, 46010 Valencia, Spain; Research School of Biology, Australian National University, Canberra, ACT 26000, Australia; CNAG-CRG, Centre for Genomic Regulation (CRG), Barcelona Institute of Science and Technology, Barcelona, Spain; Institut Català de Paleontologia Miquel Crusafont, Universitat Autònoma de Barcelona, Cerdanyola del Vallès, Spain; Catalan Institution of Research and Advanced Studies (ICREA), Barcelona, Spain

**Keywords:** snakes · genomics · *Vipera* · mito-nuclear discordance · adaptive introgression · venom evolution

## Abstract

Snake genomes attract significant attention from multiple disciplines, including medicine, drug bioprospection, and evolutionary biology, due to the unique features found in snakes, especially, the evolution of venom. However, genomic research within the family Viperidae has mostly focused to date on the subfamily Crotalinae, while overlooking Viperinae, the Old World vipers. Among Viperinae, European vipers (*Vipera*) have been the subject of extensive research because of their venoms, phylogeographic, and ecological diversification. Nevertheless, venom research in this group has been conducted using mostly proteomes alone, while phylogeography and systematics in the genus have relied on biased information from mitochondrial phylogenies. Here, we generated chromosome-level genome assemblies for three *Vipera* species and whole-genome sequencing data for 94 samples representing 15 *Vipera* taxa. This comprehensive dataset has enabled us to disentangle the phylogenomic relationships of this genus, affected by mito-nuclear discordance and pervaded by ancestral introgression. Population-level analyses in the Iberian Peninsula, where the three oldest lineages within *Vipera* meet, revealed signals of recent adaptive introgression between ecologically dissimilar species, whereas chromosomal rearrangements isolate species occupying similar niches. Finally, using transcriptomic and proteomic data, we characterized the *Vipera* toxin-encoding genes, in which opposing selective forces were unveiled as common drivers of the evolution of venom as an integrated phenotype.

## Introduction

Albeit still in its infancy, the advent of Next Generation Sequencing is finally galvanizing snake genomics, with around 20 snake genomes assembled to date^1,2^. Snake genomes are receiving increasing attention, owing to the medical importance of snakebite envenoming^3^, the potential for discovering novel bioactive compounds in the venom^4–7^, and from the perspective of fundamental research, given the extraordinary array of evolutionary novelties found in snakes^1^.

One of such evolutionary innovations is, undoubtedly, venom. Snake venom is a complex cocktail of multiple toxin families, but acts as an integrated phenotype while subduing prey^8^. However, most studies have focused on how specific venom families diversify^9–13^, settling gene duplication and positive selection as cornerstones of their evolution^1^. Altogether, the few studies envisaging the evolution of venom-encoding genes as a whole have revealed accelerated evolution in venom genes compared to non-venom paralogs^14^, especially in those most expressed^8^. Within the family Viperidae, venom research has focused on the subfamily Crotalinae^15,16^ and, more recently, Azemiopinae^17^. However, no genomic studies have been conducted on the subfamily Viperinae (the Old World vipers), contrasting with the extensive research on their venom proteomes^18^.

Within the Old World vipers, the European vipers of the genus *Vipera* have garnered attention not only for their venoms^19–22^, but also for their phylogenetic and ecological diversification^23^. This genus encompasses ca. 15 species with multiple levels of intraspecific diversification^24,25^, and displays two conspicuously different ecological affinities: a Eurosiberian set of cold-adapted species with relatively septentrional or mountainous ranges, and a Mediterranean set of warm-adapted taxa with parapatric ranges across the Mediterranean Basin (Fig. 1). The Mediterranean group comprises one species complex and three species (i.e., *Vipera ammodytes* complex, *Vipera aspis*, *Vipera latastei*, and *Vipera monticola*) with high subspecific diversity, but uncertain phylogenetic relationships among them^24,25^. In contrast, the Eurosiberian species form a well-supported monophyletic group nested within the Mediterranean vipers and divided into two clades: I) *Vipera berus*, widely distributed across the Palearctic and the vicariant *Vipera seoanei* mostly endemic to Iberia; and II) a speciose clade severely affected by taxonomic inflation, represented in Europe by the closely-related *Vipera ursinii*, *Vipera graeca* and *Vipera renardi*^24^ (Fig. 1AB).

**Fig. 1.**
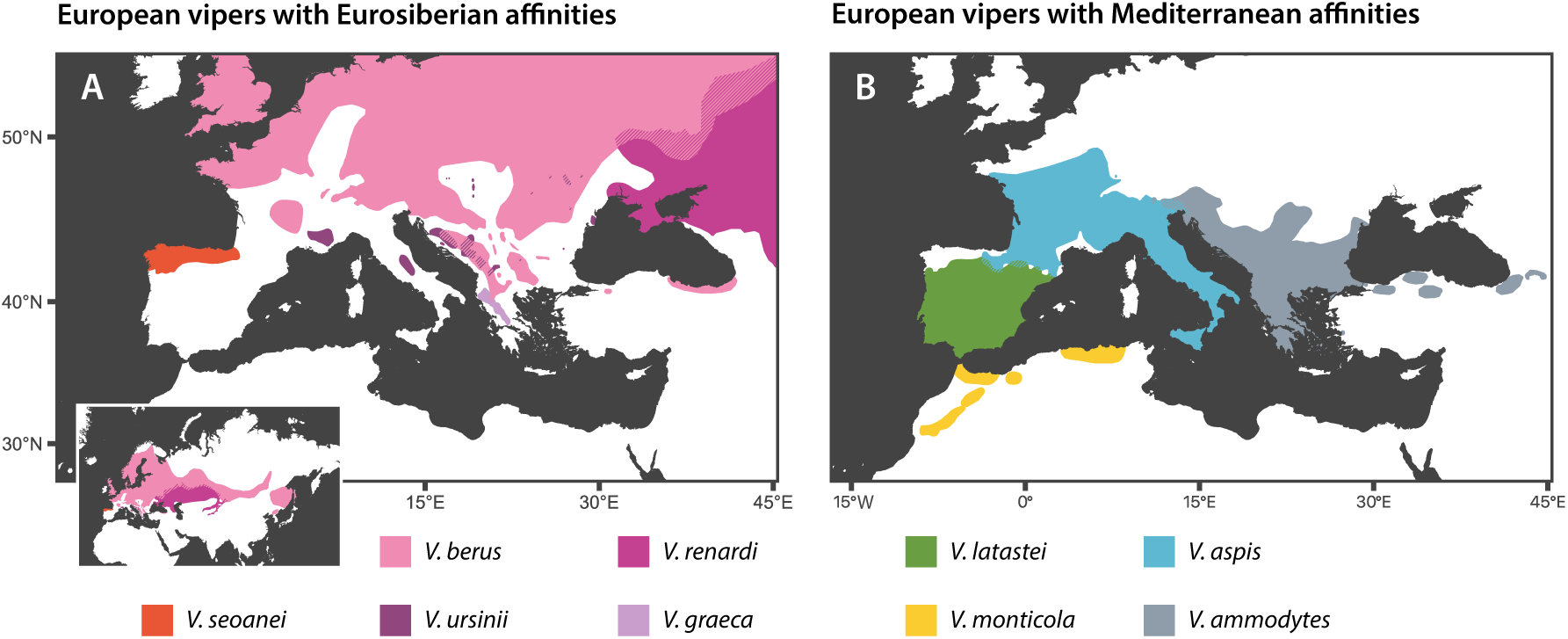
Ranges of the *Vipera* species occurring in Europe, sorted by ecological a:inities. A) Eurosiberian species inhabit septentrional regions, or disjoint mountainous areas towards the South. *Vipera berus* and *V. renardi* extend their distributions out of Europe, shown on bottom-left map. B) *Vipera* species with Mediterranean a@inities have parapatric ranges surrounding the Mediterranean Basin, including one species, *V. monticola*, endemic to North Africa.

Research on the phylogeography and systematics on this genus has heavily relied on mitochondrial DNA (mtDNA) or a few nuclear loci^26–28^. While mtDNA does not recombine and therefore cannot be used to detect hybridization or introgression events, these are nowadays considered a widespread phenomenon during the speciation process^29^. Moreover, a remarkable number of *Vipera* species has been reported to hybridize in nature^30–32^. More recently, the first genome-wide studies, with reduced-representation data, have begun unveiling introgression within the *V. ammodytes* complex^33^ and between some Eurosiberian viper species^34^. However, only one study has approached genus-level phylogenomic analysis in the genus *Vipera*, suggesting deep mito-nuclear discordance^25^, albeit biased in assumptions and focused on the widespread *V. berus*.

Here, we present chromosome-level genome assemblies for the three vipers occurring in the Iberian Peninsula: *V. aspis*, *V. latastei*, *V. seoanei*, and whole genome sequencing (WGS) data from 94 specimens representing 15 *Vipera* taxa (Table S1), including all species occurring in Europe but *V. renardi* and *V. graeca*. The aim of this work is to unravel the uncertain phylogenomic relationships of this genus and its evolutionary history, including chromosomal rearrangements, demographic oscillations, ancient introgression events and current secondary contacts among old-diverged species, for which we discuss their adaptive potential focusing on the Iberian Peninsula. Finally, we integrated available and *de novo* proteomic and transcriptomic data to explore the venom encoding-genes of the three Iberian vipers, characterizing the genes belonging to the main toxin families and analyzing the drivers behind the evolution of this complex and fascinating phenotype in the genus *Vipera*.

## Results

We generated and annotated three chromosome-level genome assemblies for *V. aspis*, *V. latastei*, *V. seoanei*. Assembly sizes are comprised between 1.57-1.60 Gb, scaffold N50 between 114-236 Mb, contig N50 between 35.3-75.9 Mb, and single-copy ortholog genetic completeness between 96.0-97.2% (Table S2, Figs. S1-S3). 99.30% of the assembly is anchored to the expected number of chromosomes in *V. latastei* (17 autosomes and Z), 99.74% in *V. aspis* (20 autosomes, with Z partially assembled into 3 scaffolds) and 100% in *V. seoanei* (17 autosomes and Z)^35–37^.

### Genomic and mitochondrial phylogenies

In order to disentangle the nuclear history of the European vipers of the genus *Vipera*, we inferred the phylogenomic relationships among seven *Vipera* species, which represent all main lineages within the genus. The only two European species not included, *V. graeca* and *V. renardi*, are closely related to *V. ursinii*^25^. We inferred Bayesian species trees from autosomal (SNAPP) and mitochondrial data to account for mito-nuclear discordance. We recovered concordant topologies for the genomic species trees in two SNAPP analyses, including or not intraspecific diversity (Figs. 2, S4-S5), although time calibrations of all nodes within *Vipera* were estimated to be much older when including a rattlesnake (subfamily Crotalinae) as outgroup. Genomic phylogenies supported the existence of three old lineages within *Vipera*, namely, *latastei*, *ammodytes* and *berus* species groups. The *berus* group comprises all Eurosiberian species, i.e., *V. berus*, *V. seoanei*, *V. ursinii*, as well as all the species not included in the study but more closely related to *V. ursinii*^25^. The *ammodytes* group, comprising the *V. ammodytes* species complex and *V. aspis*, is sister to the *berus* group, whereas the early-diverging *latastei* group comprises *V. latastei* and *V. monticola*. In this scenario, the Iberian Peninsula emerges as the only place where the three old lineages or species groups meet. The mitochondrial phylogeny is concordant with previous studies^38^, although incongruent with the nuclear species trees inferred in the present study (Fig. 2, S6).

**Fig. 2.**
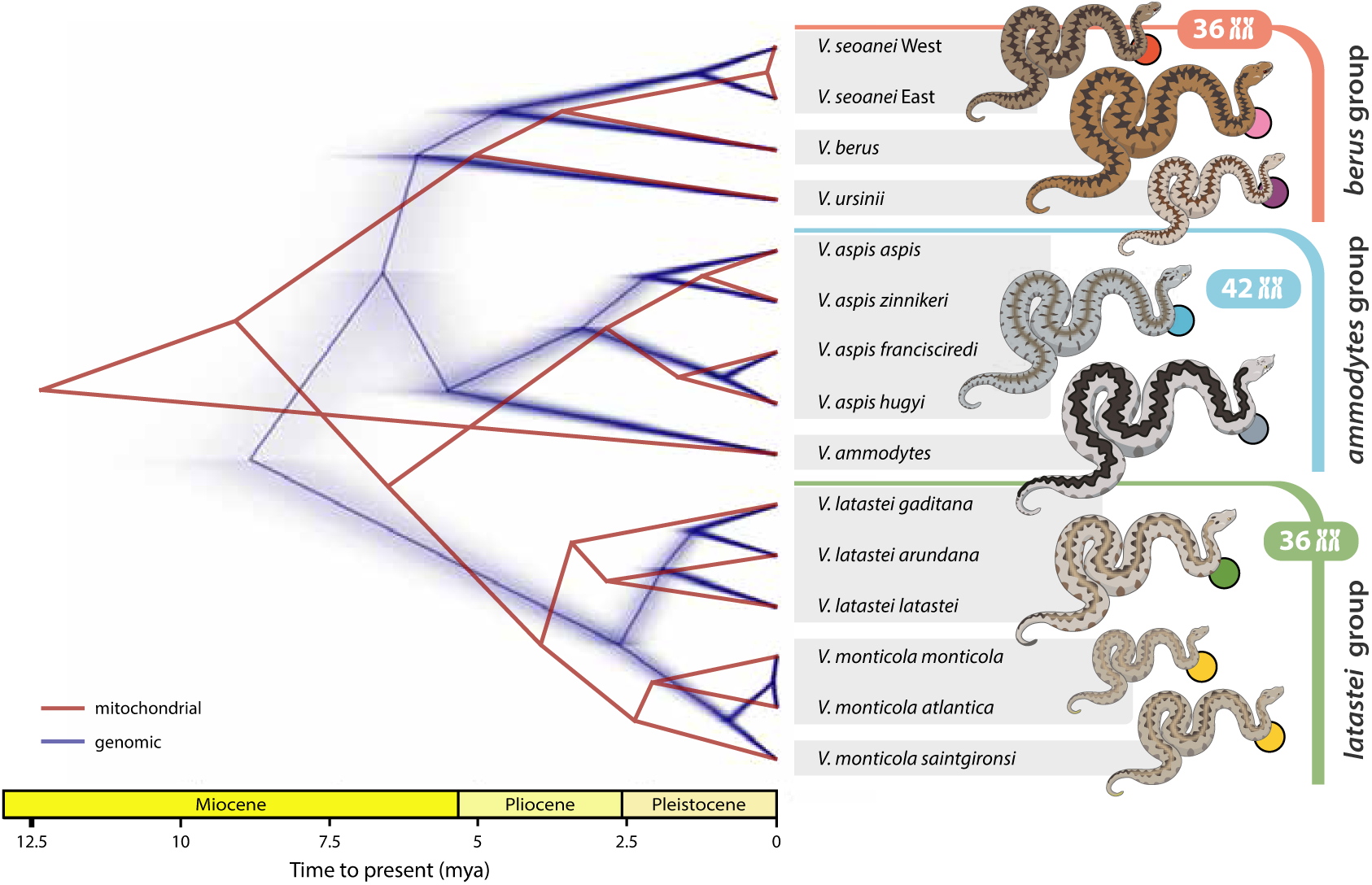
Time-calibrated phylogenomic and mitochondrial species tree of European vipers, showing mito-nuclear discordance. In dark blue, genomic species tree from SNAPP, inferred from 126k unlinked SNPs. In red, mitochondrial species tree from a 10.7-kbp alignment of all mitochondrial protein-coding genes. The genomic species tree recovers three main old species groups, namely: *berus*, *ammodytes* and *latastei* groups. In disagreement with mitogenomes, *Vipera aspis* is sister to the *V. ammodytes* complex, which agrees with the neurotoxicity of their venoms and a higher chromosome number. The *latastei* group represents the earliest diverging branch, instead of *V. ammodytes*, as the mitogenomes estimate. Thus, representatives of the three old species groups are found in the Iberian Peninsula: *V. seoanei*, *V. aspis*, and *V. latastei*. Other *Vipera* species not included in this study are more closely related to *V. ursinii*. Eurosiberian species

### Demographic inference, ROHs, and heterozygosity

We used high-coverage WGS data of the eight oldest lineages studied in this work to infer past demographic oscillations along the evolutionary history of European vipers. The analyses revealed that Eurosiberian and Mediterranean sets of species do not follow similar trends based on their biogeographic affinities. Whereas *V. aspis* West, *V. ammodytes* and *V. seoanei* seem to have thrived during the Pliocene, *V. berus* and the *latastei* group underwent population expansions during the Pleistocene (Fig. 3). Regarding *V. berus*, this expansion matches the proposed recolonization of the continent 1 mya^39^, while the following contraction might be due to a predicted lost in potential habitat during glacial maxima^23^. The demographic trends of the eight lineages clearly coalesce around 8 mya, matching the estimated divergence times with SNAPP (Fig. 2). Species with persistent population declines, like *V. ursinii*, show the highest proportion of its genome covered by short Runs of Homozygosity (ROHs), indicative of ancient bottlenecks^40^ (Fig. S7). The highest mean genome-wide heterozygosity values appear to be found in the Iberian representative of each species group (Fig. S8).

**Fig. 3.**
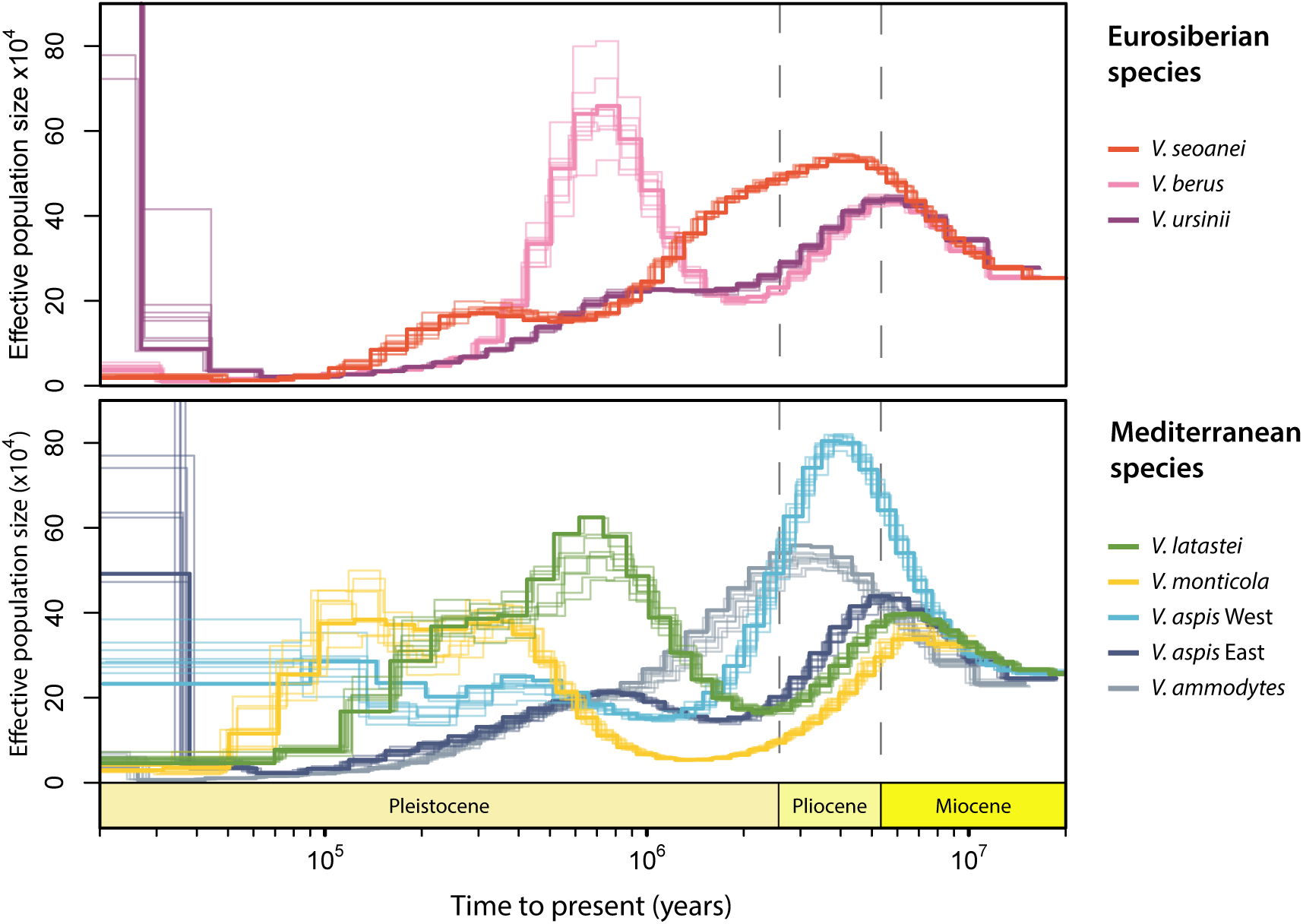
Demographic history of the studied European vipers, sorted by ecological a:inities. Thick and thin lines of same color represent the median and 10 bootstrap rounds, respectively. All species demographic inferences coalesce around 8 mya, in accordance with the time-calibrated genomic species tree (Figs. 2, S5). Eurosiberian and Mediterranean sets of species do not show higher resemblance among them than between ecological groups. Whereas *V. seoanei*, Western *V. aspis* and *V. ammodytes* seem to have thrived during the Pliocene, *V. berus*, *V. latastei*, and *V. monticola* underwent population expansions during the Pleistocene.

### Introgression analyses

We examined introgression events at subspecific level with *f-branch* implemented on Dsuite, and backed these results with TreeMix estimates, which provide insights into the directionality of the events. We found introgression to be pervasive across the genus, with seven events estimated as optimal to explain the history of the studied *Vipera* (Figs. S9-S12), and extraordinary *f-branch* values, with up to 23% of the genome in *V. seoanei* shared with *V. latastei* West (See “Population genomics” results for lineage delimitation; Figs. 4A-B, S13; Tables S3-S5). The detection of excess allele-sharing in *V. monticola* also suggests an ancient onset of this introgression, while an even higher excess between the Western lineages of both *V. latastei* and *V. seoanei* indicates that the introgression spread over time until or near the present. To delve into the introgression from *V. latastei* into *V. seoanei*, we looked at the landscape of this introgression in 25-kb windows across *V. latastei*’s genome. Although introgression is pervasive, certain areas show increased clustering, such as in chromosomes 2 and 7, whereas others, including the Z chromosome in its entirety, are all but devoid of introgression (Figs. 4C, S14). Other remarked introgression is displayed between Northern *V. monticola* and Western *V. latastei*, implying overseas dispersal, which had been already reported in some islands^41^.

**Fig. 4.**
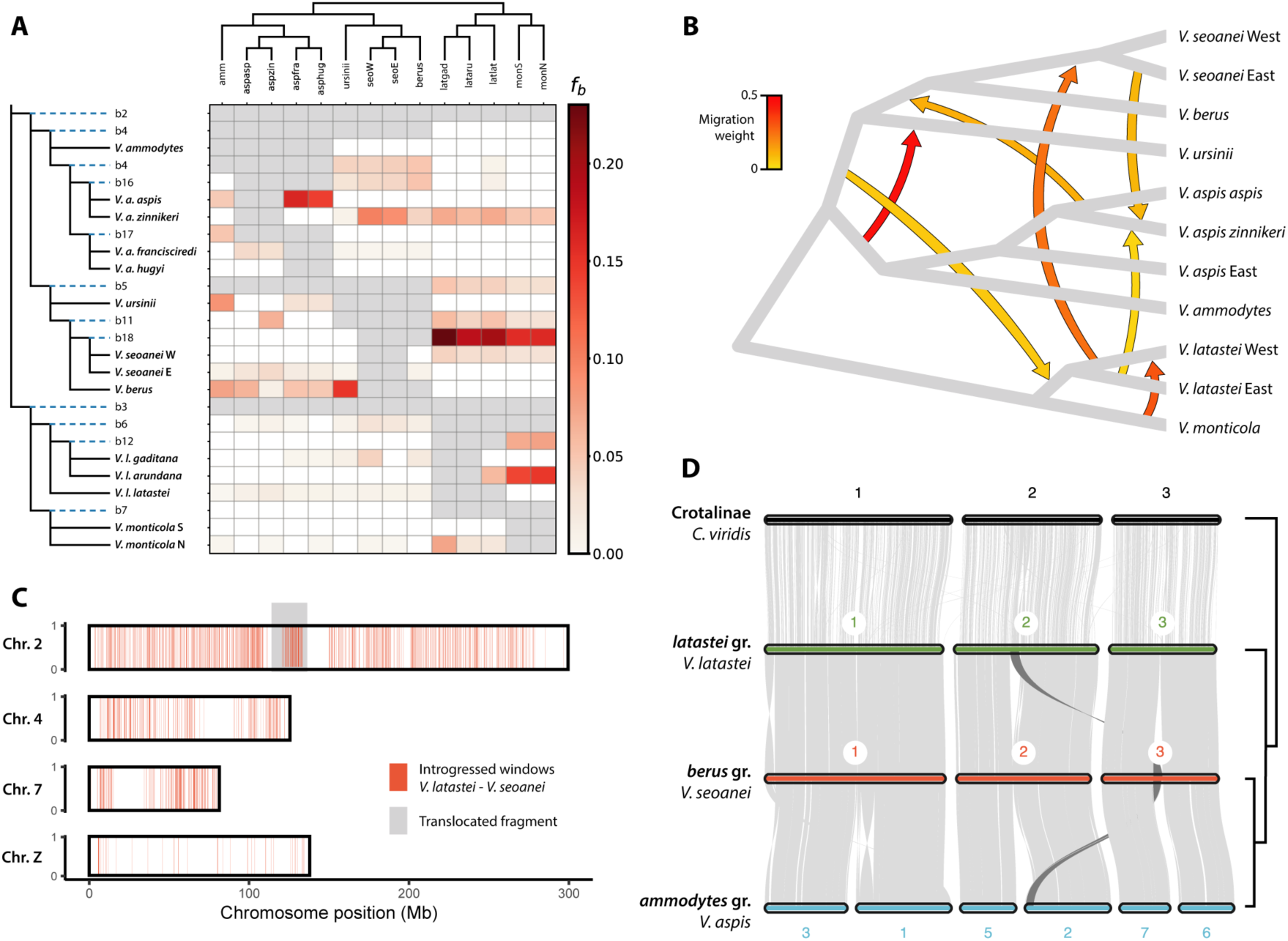
Pervasive patterns of introgression across the genus *Vipera* and macrosynteny among the three largest chromosomes across major lineages. A) Heatmap of *f-branch* (*f_b_*) statistics, representing excess allele-sharing between branches of y- and x-axes. Y-axis shows both terminal branches (in bold) and ancestral branches. Point estimates can be found in Table S4. Statistics have been corrected by a sensitive clustering threshold to avoid false positives due to homoplasy instead of introgression (see results with a robust threshold in Fig. S13). Blank cells depict values not statistically supported, whereas grey cells indicate comparisons that cannot be made due to tree topology. The most notable event of introgression seems to have happened between *V. seoanei* and *V. latastei*. B) Most likely number of introgression events within the species tree of the studied European vipers, calculated with TreeMix and OptM (Figs. S9-S12). Tree topology corresponds with Fig. 2, and inferred introgression events are consistent with *f-branch* statistic, but in this case, the direction of gene flow is inferred as well. Color of the migration edges corresponds to intensity of the geneflow event. C) Introgression landscape of the introgression event from *V. latastei* to *V. seoanei*. Windows with excess allele-sharing between both species are highlighted in *V. latastei*’s reference genome across chromosomes 2, 4, 7 and Z (see Fig. S14 for all chromosomes). Introgression between these species is pervasive, although some areas display higher intensity than others, including the whole Z chromosome. An area a@ected by an interchromosomal translocation is highlighted in darker grey (see Fig. 4D). D) Macrosynteny between the three *Vipera* species groups, represented by the Iberian species of each old lineage, and a rattlesnake as outgroup (*Crotalus viridis*, Crotalinae). Only the three major chromosomes are depicted, see Fig. S15 for the whole comparison. Chromosome architecture is greatly conserved between *berus* and *latastei* groups (2n=36) with a fragment being translocated from chr. 2 to chr. 3. Nevertheless, the *ammodytes* group (2n=42), underwent three events of chromosomal fission, which might hamper hybridization with other *Vipera* species groups.

### Macrosynteny analysis

We investigated chromosomal rearrangements between the three old species groups (represented by the three Iberian species), which may explain hybridization and introgression patterns among them. Macrosynteny analyses confirms the retention of the ancestral snake bimodal karyotype (2n=36)^42–44^ in *V. latastei* and *V. seoanei*, but reveals three fission events in the three largest chromosomes of *V. aspis*, rendering 2n=42 (Figs. 4D, S15). Despite *V. latastei* and *V. seoanei* karyotypes being almost identical, an inner fragment from *V. latastei* chromosome 2 has been translocated to chromosome number 3 of *V. seoanei*.

### Population genomics

We investigated at a population-level the Western Mediterranean, and specifically the Iberian Peninsula, as a potential crucible of ancient lineages: only in Iberian the three old species groups co-occur, with two species, *V. seoanei* and *V. latastei*, sharing a staggering excess of alleles. We used a dataset of 90 genomes at low coverage, covering all species and subspecies in the Western Mediterranean, and yielding 1.59M uSNPs, that we subsequently analyzed by PCAs (Figs. S16-S21), Admixture (Fig. 5A1-2, S22), mean heterozygosity per individual (Fig. 5B1-2, S23) and Isolation-by-Distance (IBD) models (Figs. S24-S25).

**Fig. 5.**
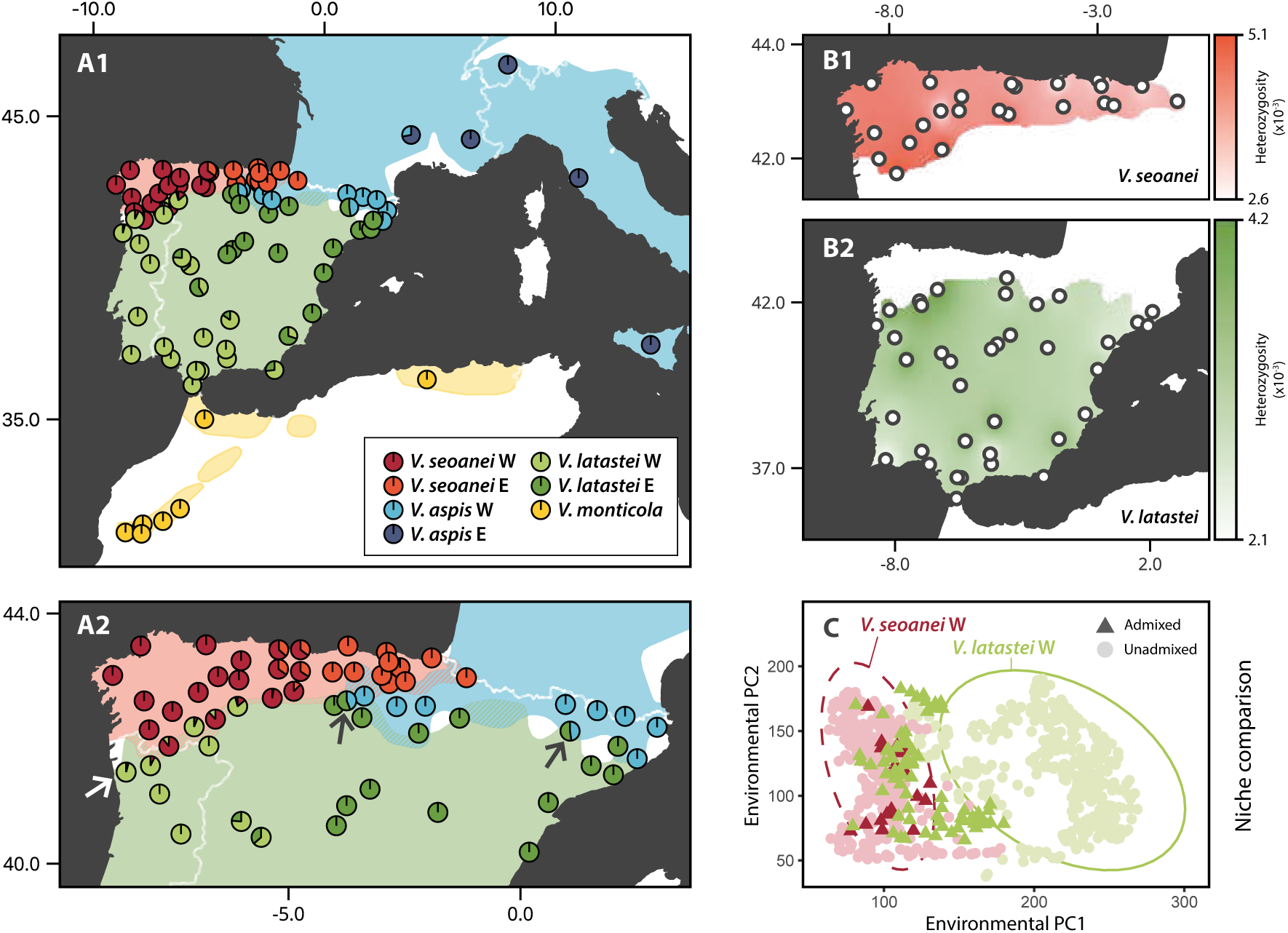
Population genomics of vipers in the Western Mediterranean, a crucible for the three *Vipera* species groups. A1-2) Admixture results for K=7, 1.59M unlinked SNPs and 90 low-coverage whole genomes, covering all subspecific diversity of these species. *Vipera seoanei*, *V. latastei* and *V. aspis* show Western and Eastern clades and exhibit di@erent inter-specific admixture patterns in Northern Iberia (A2). In NW Iberia, the Western lineages of *V. seoanei* and *V. latastei* (2n=36) show admixture despite their marked ecological di@erences. The white arrow points to a *V. latastei* individual with *V. seoanei* admixture that was sampled 70 km away from the closest *V. seoanei* population. On the other hand, on the contact zones of *V. aspis* West and *V. latastei* East, only pure or F1 generation hybrids (highlighted with dark arrows) are found despite the Mediterranean a@inities of both species, which, in addition, di@er in chromosome number. B1-2) Genome-wide heterozygosity interpolations across the distributions of *V. seoanei* and *V. latastei*. Darker colors represent higher mean heterozygosity values per individual. The highest values, for both species, are reached around the contact area in North Portugal and South Galicia (Spain). C) Climatic niche comparison of the Western *V. seoanei* and *V. latastei* lineages. Ellipses represent the dispersion with 95% CI of individuals from unadmixed populations of those species (circles). Individuals from recently-admixed populations are depicted by triangles. While putatively admixed *V. seoanei* individuals inhabit similar localities than unadmixed conspecifics, admixed *V. latastei* specimens occupy more Eurosiberian environments than expected for its lineage.

We found *V. seoanei* and *V. latastei* to sort out into Western and Eastern lineages across their ranges (Fig. 5A). For *V. seoanei*, the summits of the Cantabrian Mountains at Picos de Europa divide a more diverse lineage at the West, from a relatively impoverished lineage in the East (Fig. 5B1), partially explained by IBD (Fig. S25). *Vipera seoanei*’s type locality is comprised within the Western range, which encompasses as well the “*cantabrica*” ecotype, previously considered a subspecies^45^. For *V. latastei*, its Eastern lineage mostly corresponds to *V. l. latastei*, whereas the Western one comprises *V. l. gaditana*, the recently described *V. l. arundana*^46^ and some SE populations traditionally assigned to *V. l. latastei* (Fig. 5A1, S18). However, genomic differences are fully explained by IBD (Fig. S24). *Vipera aspis* seems better resolved by the PCA than Admixture due to sample biases, and the Alps split an impoverished Eastern lineage (*V. a. francisciredi*+*V. a. hugyi*) from two more diverged subspecies in the West: *V. a. zinnikeri* and *V. a. aspis* (Fig. S23). *Vipera monticola*, in contrast, is more homogeneous, and PC1 splits it into a Northern lineage, partially corresponding to *V. m. saintgironsi*^46^, and a Southern lineage, including Eastern High Atlas populations of *V. m. saintgironsi*, *V. m. monticola* and *V. m. atlantica*^46^ (Fig. S19).

In the northern sector of Iberia, the three old species groups meet with uneven outcomes (Fig. 5A2). Between the Mediterranean species *V. aspis* and *V. latastei*, recent admixture is barely detected, although two F1-hybrids have been identified, with half ancestry of each species (Fig. S22) and soared heterozygosity (Fig. S8). However, in NW Iberia, the ecologically-dissimilar *V. latastei* and *V. seoanei* admix in a broad area of secondary contact encompassing South Galicia and North Portugal, in which the highest heterozygosity values are found for both species (Fig. 5B1-2). Admixture even reaches a locality next to the city of Porto, ∼70km far from the closest *V. seoanei* population (Fig. 5A2).

To ascertain the adaptive character of this admixture, we used publicly available occurrence data to estimate ecological niche overlap between *V. seoanei* and *V. latastei* (Table S6) and depicted it by environmental PCAs (Figs. S26-S31, Table S7). Whereas these species and their respective lineages barely overlap (OI=9-20%, Table S6), when we include the admixed populations, their niches converge (32%). PC1 demonstrates that a Eurosiberian-Mediterranean gradient separates the niches of these species. Remarkably, individuals of *V. latastei* admixed with *V. seoanei* tend to occupy Eurosiberian niches typical of *V. seoanei* (Fig. 5C, S29). Western and Eastern intraspecific lineages were found to be similar in *V. seoanei* (62%, Fig. S31), but unexpectedly different in *V. latastei* (12%), with the Eastern lineage inhabiting areas of greater seasonality and/or drier winters (Fig. S30).

### Venom characterization and evolution

We used both transcriptomic and proteomic (new and published) data from the three Iberian vipers, and their reference genomes, to explore how toxin-encoding genes have diversified in the genus *Vipera*. Differential gene expression analyses revealed 40 genes exclusively upregulated in the venom gland, 18 of them being toxin-encoding genes. Other upregulated genes are related to muscle contraction (13), protein synthesis (3), or vacuolar fusion (1), one is an immunoglobulin and four correspond to transposable elements (Fig. S32). For *V. latastei*, genes identified through transcriptomics and through proteomics greatly overlap (Table S8), although proteomic results were more complete, validating proteomic results for the other two species (Tables S9-10). Microsynteny of the primary and secondary toxin families is depicted in Fig. S33. We found one PLA_2_ gene in *V. aspis* to be responsible of the neurotoxicity of its venom reported in some populations^47^. PLA_2_s and other primary families, more expressed in venom, do not necessarily have more gene copies than secondary families, like CTLs/snaclecs (Fig. 6B). Indeed, some families supposed to bear many copies, like SVSPs and SVMPs, do instead express multitude of isoforms out of just one gene copy (e.g., P-I, P-II, P-III and disintegrins produced by alternative splicing of the same SVMP gene). For some families, genome-wide blasts detected candidate genes to be ancestral to each family gene cluster, such as *adam28* for SVMPs^48^. In addition, next to some CRiSPs (Fig. S33), we found *serotriflin* genes, a carrier of an endogenous inhibitor (SSP-2) in serum involved in suppressing the toxic activity of CRiSPs in case of self-envenomation^49^. In total, we found 28, 16 and 41 toxin-encoding genes in the genomes of *V. latastei*, *V. seoanei* and *V. aspis*, respectively, although with similar copy numbers within main families (Fig. 6C; Tables S8–10).

**Fig. 6.**
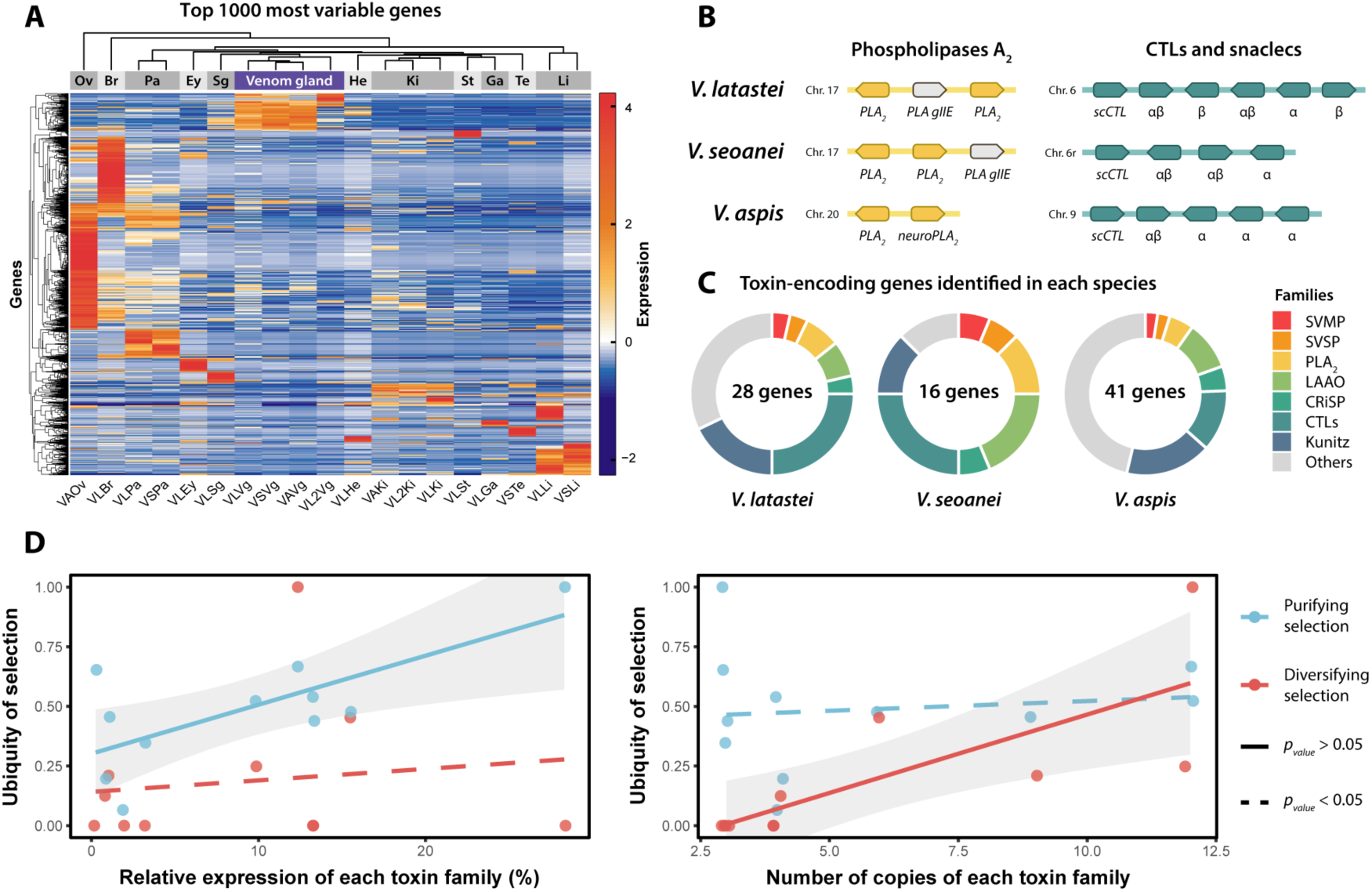
Characterization of toxin-encoding genes of *V. latastei*, *V. seoanei* and *V. aspis*, and venom selection analysis. A) Heatmap of di@erential gene expression analyses, from di@erent organs of the mentioned species (Fig. S35). Y-axis is composed by the top 1,000 most variable genes, and x-axis shows RNA-Seq samples from di@erent organs and tissues. Samples from same organs cluster together, such as the venom glands coming from four specimens of the three species. Red hues indicate upregulation, whereas blue colors downregulation. *Ov* stands for ovary, *Br* for brain, *Pa* for pancreas, *Ey* for eye, *Sg* for salivary gland, *He* for heart, *Ki* for kidney, *St* for stomach, *Ga* for gall bladder, *Te* for testes and *Li* for liver. B) Microsynteny of two selected toxin families in the three Iberian species. Here, only Phospholipases A_2_ and C-type lectins (CTLs) + snaclecs are shown, but all main toxin families are depicted in Fig. S33. PLA gIIE are non-toxic, but related to toxic PLA_2_. PLA_2_ is a main family in the venom of vipers, but only two copies are found in the genome of these species (one of them producing neurotoxic PLA_2_, in *V. aspis*). CTLs, albeit composing a secondary family in terms of total expression in the venom, is richer in gene copies. scCTL stands for the ancestral (albeit venomous) single-chain CTLs, whereas α, β and αβ represent snaclec genes producing either one or both subunits of the snaclec heterodimer. C) Pie charts of all characterized venom-encoding genes in each of the three reference genomes by toxin family (Tables S7-9). D) Selection analyses on the identified toxin-encoding genes. Toxin families with a higher expression undergo a significantly stronger purifying selection. On the other hand, the diversifying selection only increases with the number of gene copies within the family.

Finally, selection analyses revealed episodic selection in all main toxin families but 5’-nucleotidases, svVEGF and svNGF, being the latter two involved in fostering existing prey’s metabolic pathways, instead of disrupting them^50,51^ (Table S11). Regarding pervasive selection, the toxin-encoding gene families that are more expressed, undergo a stronger purifying selection (Fig. 6D; adjusted r^2^=0.47; *p_value_*=0.012), whereas diversifying selection does not depend on the importance of the family (*p_value_*=0.692). Indeed, diversifying selection increases with the number of copies within a family (Fig. 6D; adjusted r^2^=0.54; *p_value_*=0.006), whereas this feature does not correlate with purifying selection (*p_value_*=0.730).

## Discussion

In the present work we have generated three genome assemblies with unprecedented levels of contiguity in Viperidae. Although scaffold N50 values in *V. latastei* and *V. seoanei* are similar to other snake assemblies (Table S2), this parameter, widely used to compare chromosome-level assemblies’ contiguity^1,52^, is dependent on genome architecture. Thus, these two species, with three macrochromosomes accounting for half of their genome, hold unsurprisingly higher values than *V. aspis*, in which these chromosomes are fragmented due to fission. However, contig N50 better depicts contiguity in chromosome-level assemblies. While *Crotalus tigris* was probably the most contiguous viperid genome prior to this work^15^, these three *Vipera* genomes are 16-36 times more contiguous (Table S2).

### The evolutionary history of European vipers

Here we have first estimated the phylogenomic relationships among the European vipers of the genus *Vipera* from WGS data, revealing the monophyly of *V. aspis*+*V. ammodytes*. This is supported by two key synapomorphies: three extra chromosome pairs^36^ and significant neurotoxicity of their venoms^47^. Although *V. berus* has been as well reported to be neurotoxic in some localities^53^, this could be a case of either convergence or introgression. This new topology is incongruent with the only previous genus-level study using genomic data, which used 1.1 Mbp to estimate phylogenomic relationships but incorrectly assumed that *V. ammodytes* was the root^25^, as shown by mitochondrial data.

The discovered mito-nuclear discordance in the phylogeny can arise from multiple sources, such as incomplete lineage sorting (ILS), adaptive introgression of mtDNA or sex-biased asymmetries^54^. A phylogeny is more prone to ILS when a rapid succession of speciation events occurs while ancestral population sizes are large, and both circumstances are met according to our demographic inference results (Fig. 3), which could explain a closer mitochondrial relationship of *V. aspis* to the *latastei* group than to the *berus* group. However, the largest mito-nuclear disagreement is the placement of *V. ammodytes* as sister taxon to all other *Vipera* species included in the analysis. This might be explained by the mitochondrial capture through introgression from an archaic ghost lineage, which was donor of its mtDNA but likely only small portions of its nuclear genome. This overlooked phenomenon is now thought to be widespread in nature^55^, and has even been reported in a species of the *berus* group^56^. Further research should address it in the case of the *V. ammodytes* complex.

Mitochondrial sweeps could as well have caused the shallow mitochondrial structure between Western-Eastern *V. seoanei*, despite important genomic differences (Fig. 2). This species seems to have re-expanded from West to East during the Last Glacial Maximum^45^, but genomic differences would suggest that relict populations in the East contributed somewhat with their genomic material while capturing Western mtDNA. The contrary case is deep mitochondrial divergence (DMD), that reflects long periods of geographical isolation followed by secondary contacts^57^. DMD is fostered in the face of high levels of geneflow, especially if there is strong male-biased dispersal and strict female philopatry^55^, as described in the European vipers^58,59^. This is likely the case of *V. monticola* and, to a lesser extent, *V. latastei* subspecific lineages, whose ancient mitochondrial divergences are not congruent with a relatively shallower nuclear structure, byproduct of IBD.

### Pervasive introgression across European vipers

We have found introgression to pervade the phylogeny of European vipers at multiple levels (Figs. 4ABC-5A). Nevertheless, hybridization among *Vipera* spp. seems to be avoided by a) niche segregation and b) chromosomal rearrangements. On the one hand, *Vipera* ranges in Europe are mainly parapatric (Fig. 1), and in the narrow borders of sympatry between species, they can segregate by selecting slightly different habitats and microhabitats, having non-overlapping mating periods or different thermoregulatory strategies^60–63^. However, latitudinal differences in Europe used to be weaker in the past, with the Mediterranean regime only beginning in the Late Pliocene (3.2 mya^64^). This could have led to broad areas of sympatry, promoting ancient introgressions, and is illustrated by the lack of common demographic oscillations within the Eurosiberian and Mediterranean sets of species (Fig. 3). On the other hand, niche convergence instead of segregation has been observed only between sympatric *V. latastei* and *V. aspis*^65,66^, two species with differing karyotypes. Individuals that are heterozygous for chromosomal rearrangements are partially or completely sterile, quickly generating a strong reproductive isolation^67,68^, in line with F1-hybrids coexisting with pure parental individuals in sympatric localities (Fig. 5A) (although see^30,69,70^ for putative F_2_ specimens).

If reproductive isolation mechanisms have arisen among most *Vipera* spp., it is striking how the situation differs between *V. latastei* and *V. seoanei*. After divergence from *V. berus*, *V. seoanei* possibly underwent a rapid niche shift towards a less-cold Atlantic habitat in contrast to its boreal sister species, and even adapted to an aestival spermatogenesis, typical in Mediterranean vipers like *V. latastei*^23^. Adaptation to new environments can be achieved by introgressive hybridization or new mutations if such variation does not pre-exist in the species. Introgression, though, leads to faster adaptation than mutation, providing genetic variants that have already been tested by natural selection^71^. Indeed, the highest genomic variance values per individual in both species have been found in their secondary contact zone in NW Iberia (Fig. 5B1-2; S24). Higher heterozygosity values are often expected in genome regions involved in adaptive introgression, as both correlate with higher recombination rates^72^.

Different areas across the genome yield differential effects on fitness when introgressed, shaping what is called the introgression landscape^73^. Whereas some areas seem to be more prone to accept the movement of genes, others appear devoid of foreign material (“introgression deserts”^72^). For *V. seoanei*, the Z chromosome remains almost free from *V. latastei* introgression (Fig. 4C; S14), agreeing with multiple genomic studies that have found evidence on reduced introgression on sex chromosomes^74–77^, suggesting a major role of the Z chromosome in the reproductive isolation of *Vipera* spp. However, some weaker areas of introgression could be simply linked to lower recombination rates^76^ (e.g., around the interchromosomal translocated fragment, Fig. 4C-D), and some introgression deserts found in many autosomes might correspond to centromeres (Fig. S14). Finally, compelling evidence on the adaptive nature of an introgression event is niche expansion following hybridization^78^. Although recently admixed *V. seoanei* do not occupy a different niche from their conspecifics, *V. latastei* individuals admixed with *V. seoanei* have successfully occupied areas of Atlantic climate similar to the latter species (Fig. 5C; S29).

### A fine balance of opposing forces drives venom evolution

Diversifying selection and gene duplication are regarded as the cornerstones of the toxin arsenal assembly in snakes. Some of the most important toxin families, like PLA_2_s and SVMPs, are thought to have been recruited as venom genes after undergoing duplication^79^, which is supported too in *Vipera* by non-venomous paralogs next to venomous copies (Fig. S33). Other toxin genes, however, arose by direct co-option without such ancestral gene duplications, like many secondary families or the SVPSs, whose co-option was facilitated by the activity of transposable elements^79,80^. Interestingly, four transposable elements were found to be differentially upregulated in the *Vipera* venom gland, with one located next to an SVSP gene, in line with this hypothesis. Regardless of the origin of the toxin genes, gene duplication was thought to extend further: the most important families seemed to have been expanded afterward their recruitment/co-option as a signature of tacit selection in Viperidae^48^.

However, the genus *Vipera* represents a paradigm shift, as it demonstrates that the most abundant toxin families are not necessarily the most duplicated ones, nor those in which evolution has accelerated the most. Although we cannot ascertain whether some of those families never underwent gene duplications or instead their ancestor suffered gene deletions, as reported in some rattlesnakes^15^. In this new scenario, where these circumstances are untangled, we can better envisage the interplay between gene duplication and diversifying selection. Hence, evolution accelerates once gene duplication has occurred, i.e., there are spare copies to allow innovation. Otherwise, purifying selection governs the evolution of these genes, especially the most important ones (Fig. 6D), whose maintenance is in turn vital for the individual’s fitness. In contrast to diversifying selection, purifying selection has received little attention in snakes, and has only been hypothesized to constrain venom evolution in ancient lineages like cnidarians and scorpions^81^. However, purification stages might be more widespread than expected, even in relatively “young” lineages like snakes, and especially in the absence of ongoing ecological specialization.

## Material and Methods

More detailed information on the Materials and Methods used in this work are provided in a **SI Appendix**.

### Reference genome sequencing, assembly, and annotation

In brief, the reference genomes of *V. latastei*, *V. seoanei* and *V. aspis* were sequenced and assembled with data from different platform combinations, among ONT and PacBio HiFi long reads, Illumina short reads and Omni-C, Hi-C contact map or Bionano optical mapping (see SI Appendix for assembly pipelines), yielding 18, 18 and 21 chromosomes, respectively. For *V. seoanei* we scaffolded contigs using the closely related *V. ursinii* as reference instead. Genome assemblies were depicted as snailplots with BlobToolsKit (Figs. S1-3), and their contiguity and completeness were assessed with gfastats ^82^ and BUSCOs vertebrate database^83^. In addition, data from 19 RNA-Seq samples from twelve different tissues of the three species, full-length transcriptome Iso-Seq data from pooled tissues and venom glands, as well as protein databases and other snake genomes served as evidence to annotate the *Vipera* genomes following Gene-MarkS-T, BRAKER 1 and 2, GeMoMa and Tsebra pipelines^84–88^.

Finally, we explored macrosynteny between the three Iberian reference genomes, and *Crotalus viridis* as outgroup^89^ with MCscan^90^. Protein sequences were aligned between genomes with LAST^91^ and some chromosome reorientation was carried out to correctly visualize chromosomal rearrangements and identify the sexual chromosomes.

### Whole-genome sequencing data production

Furthermore, we build genomic libraries following Carøe et al.^92^, to sequence five additional genomes (apart from the three references) at high coverage (∼40x) with 150-paired end (PE) Illumina short reads to represent other species/lineages within the genus (i.e. *V. ursinii*, *V. berus*, *V. ammodytes*, *V. monticola* and Eastern *V. aspis*). Other 86 genomes were sequenced at low coverage to explore finer-scale population structure and introgression patterns focusing on the Western Mediterranean, including all subspecies within *V. latastei*, *V. monticola* and *V. aspis*. Although *V. seoanei* is thought not to harbor subspecific diversity, its range was exhaustively sampled as well^45^.

Afterwards, WGS data was quality filtered and adapter-trimmed with fastp^93^ and mapped to *V. latastei* reference genome (rVipLat1) with BWA^94^. PCR duplicates were removed with PicardTools and we called SNPs and filtered the resulting dataset with GATK^95^. Further filtering with VCFtools^96^ for subsequent analyses included filtering by missingness, keeping just autosomal bi-allelic SNPs, purging singletons and Linkage-Disequilibrium pruning, whose decay function was calculated with PopLDdecay^97^ (Fig. S34, Table S12). Finally, mitochondrial sequences were extracted and assembled from WGS data from 15 representatives of different taxa within *Vipera* with GetOrganelle^98^. Resulting mitogenomes were annotated with MitoFinder^99^.

### Phylogenomic and phylogenetic analyses

First, genomic species trees were inferred with SNAPP implemented on BEAST2^100^ for two unlinked SNP (uSNPs) datasets without missingness, at species- and subspecies-level, with 1,225k and 126k uSNPs, respectively (Table S12), and three runs of 20M iterations for each dataset. These phylogenies were time-calibrated with the estimated divergence between Viperinae and Crotalinae (38.61±4.85 mya), for the species dataset; and the crown diversification of the *V. berus*+*V. seoanei*+*V. ursinii* clade, i.e. the *berus* group (5.97±0.75 mya), for the subspecies phylogeny without outgroup, following fossil-calibrated phylogenetic studies^38^. On the other hand, from the mitogenomes of all subspecies plus *Crotalus* as outgroup, we aligned their protein coding genes, investigated the best partition scheme with the greedy algorithm of PartitionFinder^101^ and ran three runs of BEAST2 to estimate the mitochondrial tree, accounting for 150M iterations in total. We time-calibrated this last dataset with the crown age of the *berus* group as well. The mitochondrial and genomic trees were depicted overlapped for a better visualization of mito-nuclear discordance.

### Demographic inference, Heterozygosity and Runs of Homozygosity

To explore the influence of climate changes on the demographic history of the European vipers, we ran PSMC on each high-coverage representative sample of the studied *Vipera* spp., following parameter selection by Schield et al.^102^, a rate of 2.4e-9 substitutions/site/year^103^ and a 6-year generation period^104^. Ten bootstrap replicates were calculated per genome, and results were divided into Eurosiberian or Mediterranean species according to ecological affinities.

In addition, we calculated mean autosomal heterozygosity per individual throughout 100-kbp windows, following Mochales-Riaño et al.^105^, using 94 low-coverage genomes. Means per species were plotted as bar plots with ggplot2^106^. Runs of homozygosity (ROHs) across autosomes for the 8 high-coverage samples were estimated with bcftools^107^, filtered by quality (PhredScore>70), sorted out by length, and plotted as a percentage of the total autosomal genome size.

### Estimating interspecific introgression

We explored interspecific introgression among identified *Vipera* lineages/subspecies by means of *f-branch* statistics, derived from ABBA-BABA tests in Dsuite^108^. We excluded F1 hybrids and individuals resulting from secondary contacts between intraspecific lineages (see Admixture results), and included a *Crotalus* species as outgroup, yielding a low-coverage dataset of 86 individuals and >12M linked SNPs. We tested both sensitive and robust clustering thresholds implemented in Dsuite to avoid false positives due to ABBA homoplasy and plotted the statistically significant results. Secondly, we ran TreeMix^109^ to investigate the direction of the introgression events on a subspecies dataset. Models with different migration edges between 1-10 were estimated by blocks of varying numbers of SNPs, and the optimum number of edges was estimated with OptM package^110^ in R. Finally, we investigated the landscape of the most important introgression event in the phylogeny with TWISST^111^, highlighting the windows supporting introgression between *V. seoanei* and *V. latastei-monticola*.

### Population genomic analyses in the Western Mediterranean

First, we explored with genomic PCAs the population structure of Western Mediterranean vipers, using data from 90 low-coverage whole genomes of four species. Following, we delve into the population structure and potential inter-specific secondary contacts with Admixture^112^ (K=1-12). We depicted the optimum K as individual pie charts in the map. In addition, for the two best-sampled species: i.e., *V. latastei* and *V. seoanei*, we examined whether the discovered intra-specific structure could be explained by Isolation-by-distance patterns with the R package prabclus^113^. In addition, per-individual mean heterozygosity values for the three Iberian species were estimated and used to interpolate a heterozygosity raster of each species. Finally, we inferred the climatic niches and investigated niche overlap between these two species and their respective lineages, both excluding or including localities assigned to admixed populations. We extracted 19 environmental variables from WorldClim2^114^ belonging to *V. latastei* (n=1,105) and *V. seoanei* (n=508) and summarized them into climatic PCs. Niche overlap was estimated with both Sørenson and Overlap Indexes^23,26,115^, using the R package hypervolume^116^, and visualization was performed with ggplot2^106^ through 2-D PCAs. We highlighted admixed individuals to illustrate if their niches are congruent with the expected niches for non-admixed conspecifics.

### Characterizing venom

In brief, we integrated transcriptomic and proteomic data from the three species to characterize the toxin-encoding genes of the Iberian vipers. On the one hand, RNA-Seq data from 19 samples from venom glands and other tissues (i.e., gonads, brain, pancreas, eye, salivary gland, heart, kidneys, stomach, gall bladder and liver; Fig. S35) were used to identify positively upregulated genes exclusive to the venom gland (common to the three species, but with *V. latastei* as reference). We used DESeq2^117^ to standardize gene expression and test for significantly upregulated genes (log2 fold-change>2) in the venom glands *vs* all other tissues, correcting for multiple tests (FDR: 5%).

On the other hand, we used available proteome data from venoms of *V. seoanei*^21^ and *V. aspis*^19^, as well as *de novo* produced proteomes from three *V. latastei* (see SI Appendix). We looked for exact matches of the peptide sequences of each specific proteome among the translated proteins annotated in its respective genome to identify the genes involved in the expression of those venoms. Venom-encoding genes derived from proteomics were compared to those derived from transcriptomics in the case of *V. latastei* to verify that proteome data is accurate and sufficient.

Finally, we performed d_N_/d_S_ ratio tests with BUSTED^118^ and FEL^119^ analyses implemented on Datamonkey^120^ to look for episodic diversifying selection and pervasive selection, respectively, in each toxin-encoding family. The dependence of the corrected relative ubiquity of pervasive diversifying or purifying selection from FEL tests was assessed based on the number of gene copies within a family and the abundance of this family in the venom composition (Table S13). Linear regressions were plotted with ggplot2.

## Data and Material Availability

Genome assemblies, mitogenomes, WGS, Iso-Seq, RNA-Seq raw data and *V. latastei* proteomes will be released upon publication of this manuscript.

### Sampling permits

Samplings were authorized in: Morocco by the Haut Commisariat aux Eaux et Forêts et à la Lutte Contre la Désertification of Morocco, with permission numbers HCEFLCD/DLCDPN/DPRN/CFF N° 20/2013, 19/2015, 35/2018 and 12/2022; Portugal by the Instituto da Conservação da Natureza e das Florestas with permission references 537/2018, 295/2020 and 326/2022; Andalucía by Consejería de Agricultura, Ganadería, Pesca y Desarrollo Sostenible from Junta de Andalucía, with permission reference SGYB/DBP; Álava by Departamento de Medio Ambiente y Urbanismo from Arabako Foru Aldundia/Diputación Foral de Álava, with permission number 2021/017; Aragón by Departamento de Agricultura, Ganadería y Medio Ambiente from Gobierno de Aragón, with verification code CSVR3-6PK6D-0KYB4-FWREG; Principado de Asturias and Picos de Europa National Park by Consejería de Medio Rural y Cohesión Territorial from Gobierno del Principado de Asturias, with permission numbers DECO/2021/9850 and CO/09/0230/2021, respectively; Bizkaia by Departamento de Sostenibilidad y Medio Ambiente from Diputación Foral de Bizkaia with permission number AM-30-2021; Cantabria by Consejería de Desarrollo Rural, Ganadería, Pesca, Alimentación y Medio Ambiente from Gobierno de Cantabria, with registration number 2021GA001S023262; Castilla-La Mancha by Dirección General de Medio Natural y Biodiversidad from Consejería de Desarrollo Sostenible, with permission number DGMNB/SEN/avp_21_204; Castilla y León by Consejería de Fomento y Medio Ambiente from Junta de Castilla y León, with permission numbers EP/CyL/94/2016, 56/2018, 192/2020 and AUES/CyL/198/2021; Catalonia and Els Ports Natural Park by Direcció General de Polítiques Ambientals i Medi Natural from Generalitat de Catalunya, with permission numbers SF/0118/2019 and SF/761, respectively; Extremadura by Consejería para la Transición Ecológica y Sostenibilidad from Junta de Extremadura, with permission number CN0034/21/ACA; Galicia by Consellería de Medio Ambiente, Territorio e Vivenda from Xunta de Galicia, with permission numbers EB-017/2019, 018/2020, 015/2021, and 048/2021; Comunidad de Madrid by Consejería de Medio Ambiente, Vivienda y Agricultura from Gobierno de la Comunidad de Madrid, with permission number 10/283514.9/21; Comunitat Valenciana by Consellería de Agricultura, Desarrollo Rural, Emergencia Climática y Transición Ecológica from Generalitat Valenciana, with permission number 926/2021-VS (FAU 31_21); Comunidad Foral de Navarra by Departamento de Desarrollo Rural y Medio Ambiente from Gobierno de Navarra, with reference 177E/2021; Región de Murcia by Consejería de Agua, Agricultura, Ganadería Pesca y Medio Ambiente from Gobierno de la Región de Murcia, with permission number AUF/2021/045. Euthanasia of selected individuals was authorized by Consejería de Medio Ambiente, Vivienda y Ordenación del Territorio from Junta de Castilla y León with code AUES/CYL/228/2023 for *Vipera latastei* and *V. seoanei*, and Direcció General de Polítiques Ambientals i Medi Natural from Generalitat de Catalunya with code SF/0199/22 for *V. aspis*.

## Acknowledgements

We must thank Juan Pablo González de la Vega, Juan Timms, Rodrigo Bustos, Emiliano Mori, María Casero (Centro de Recuperação e Investigação de Animais Selvagens (RIAS) - Associação ALDEIA (Portugal)), Ricardo Brandão (Centro de Ecologia, Recuperação e Vigilância de Animais Selvagens (CERVAS) - Associação ALDEIA (Portugal)), Luis Guilherme Sousa (HerpEbora (Portugal)), Alberto Sánchez Vialas and Marta Calvo (Museo Nacional de Ciencias Naturales – CSIC (Madrid, Spain)), Inês Freitas, Nahla Lucchini, Soumia Fahd, Abdellah Bouazza, Jon Buldain, Ismael Espasandín, Fèlix Amat, David Donaire, Neftalí Sillero, João Campos, Víctor Ortega, Marc Albiac, Egoitz Ikaza, Aritz Ibarzabal, and Paco Moreno for their help in sample collection. We thank as well Jonathan Wood for his thorough comments on genome assembly manual curation, Klara Eleftheriadi for her kind help in genome assembly and annotation, Ignazio Avella for providing the *V. seoanei’*s proteome, and Ignacio O’Mullony for his graphic design advice. Also, we thank CNAG genome assembly team, Tyler Alioto, Jessica Gómez and Fernando Cruz for their comments on genome assembly, and Sandra Ruibal and Laia Llovera for their help in WGS library preparation.

This work was funded by grant PID2021-128901NB-I00 funded by MCIN/AEI/10.13039/501100011033 and by ERDF, a way of making Europe, and grant 2021 SGR 00420 from the Departament de Recerca i Universitats de la Generalitat de Catalunya to SC. AT is supported by “la Caixa” doctoral fellowship programme (LCF/BQ/DR20/11790007). MP-F is supported by “la Caixa” doctoral fellowship programme (LCF/BQ/DR20/11790032). FM-F is supported by FCT – Fundação para a Ciencia e Tecnologia de Portugal (ref. DL57/2016/CP1440/CT0010; DOI: 10.54499/DL57/2016/CP1440/CT0010). BB-C was funded by FPU grant from Ministerio de Ciencia, Innovación y Universidades, Spain (FPU18/04742). GM-R is funded by an FPI grant from the Ministerio de Ciencia, Innovación y Universidades, Spain (PRE2019-088729). ME is funded by an FPI grant from the Ministerio de Ciencia, Innovación y Universidades, Spain (PRE2022-101473). Some fieldwork campaigns have been supported by the Natuurmuseum Fryslân of Leeuwarden, Netherlands.

