## Supplementary Methods for "Unveiling the evolutionary history of European vipers and their venoms from a multi-omic approach"

### Supplementary Information. Material and Methods

#### Sampling

We extracted blood samples from four different individuals to produce the sequencing data for three reference genomes: a male (CNPOR1) *V. latastei* from Els Ports (Catalonia, Spain), a female and a male (CNMOL1) *V. aspis* specimens from La Molina (Catalonia, Spain) and a male (CNSOC1) *V. seoanei* from Socobio (Cantabria, Spain). Another four specimens were euthanized (a female *V. aspis* from La Molina, VA; a male *V. seoanei* from Burgos, VS; and two male juvenile and adult *V. latastei* from Burgos as well, VL and VL2, respectively) to collect their tissues, which were flash-frozen at -80ºC until RNA extraction. Regarding WGS, samples from 94 individuals of different European and North African viper species of the genus *Vipera*, including *V. latastei*, V. *seoanei*, V. *monticola*, *V. aspis*, *V. berus*, *V. ursinii* and *V. ammodytes* were sequenced in the present study (See Table S1). We focused on the Western Mediterranean diversity, including representatives from all described lineages of *V. latastei*, *V. seoanei*, *V. aspis* and *V. monticola*, specially aiming to cover most of the Iberian Peninsula and the contact zones among species. Samples varied from tail tips, tissue from road killed animals, museum specimens or scale clips, and were stored in 99% ethanol at -20ºC. Finally, venom samples were collected in 2018 from three *Vipera latastei* specimens in Segovia (adult female, Vl2), Hervás, Cáceres (adult male, Vl14), and Navezuelas, Cáceres (adult male, Vl5).

#### DNA extraction, library preparation and sequencing

We extracted high molecular weight genomic DNA from diverse tissues (tail tips, ventral scale clips or blood, mainly) from wild and museum specimens of 94 individuals of the genus *Vipera*. We followed manufacturer's protocol of MagAttract HMW Kit (Qiagen), sonicated samples with QSonica Q800R3, built whole genome libraries following the BEST method[1] with minor adjustments, and quality-checked with Bioanalyzer (Agilent). We sequenced 150bp pair-end reads aiming for 10x coverage for 86 samples in Illumina NovaSeq6000 lanes from Macrogene and Novogene Co. Five samples were sequenced at high coverage (45x) as representatives of their species, and another three samples were sequenced at very high coverage (60-75x) for reference genome assembly purposes (see Table S1). Secondly, long-read Oxford Nanopore Technologies (ONT) data was sequenced from blood DNA extractions of male *V. latastei*, *V. seoanei* and *V. aspis* at CNAG, aiming for 60x coverage for each species. Additionally, 60x long-read data from PacBio HiFi was sequenced for *V. latastei*. Finally, we combined Bionano optical mapping (248 cmaps) and Hi-C data (100x) to scaffold *V. latastei*'s draft genome into pseudochromosomes. On the other hand, for *V. aspis* we built Omni-C libraries from another female conspecific's blood using Omni-C kit by Dovetail Genomics and sequenced 150bp PE reads with Illumina NovaSeq6000 (CNAG) aiming for 120x coverage.

To annotate the reference genomes and to explore differential gene expression among organs, four individuals (young and adult *V. latastei*, *V. aspis* and *V. seoanei*) were sacrificed and their individualized tissues (heart, lungs, brain, venom gland, salivary gland, tongue, liver, eye, gall bladder, kidney, stomach, pancreas, and gonads) were flash frozen with liquid nitrogen. RNA was extracted with HigherPurity™ Tissue Total RNA Purification Kit (Canvax) and quality-checked with Bioanalyzer. The 19 best-preserved RNA samples (including 4 venom glands) were used for building stranded RNA libraries with poly-A enrichment. 12 Gb of data per sample were sequenced with Illumina short reads in NovaSeq PE150 (Novogene Co.). Furthermore, for each species, one pooled non-venomous RNA sample and one venom gland sample (pooled for adult and young *V. latastei*) were sequenced with PacBio long reads (Iso-Seq) to obtain HiFi full-length transcriptomes for a better isoform annotation. In total, six samples of this project and two external ones were sequenced on a PacBio HiFi SMRT Cell 8M using the Sequel II sequencing kit (Novogene Co).

**Genome assemblies**

*Vipera latastei*'s PacBio HiFi reads were trimmed from PacBio adapters using Cutadapt v3.4[2]. ONT long reads for the three species were quality checked with nanoQC[3]. To obtain the draft genome assemblies, *V. latastei* PacBio reads were assembled using Hifiasm v0.13[4] with default settings, and *V. seoanei* and *V. aspis* long read data above 8 kbp was assembled using NextDenovo v2.5.2[5]. Illumina short read data was quality checked with FastQC v0.11.9[6] and multiQC[7], adapters were trimmed with fastp v0.20[8] and mapped to the respective draft assembly with BWA v.0.7.17-r1188 mem algorithm[9], and used to polish *V. seoanei* and *V. aspis* drafts with two rounds of Pilon v1.24[10]. Afterwards, haplotigs were purged from the three draft assemblies with *purge_dups* v1.2.5[11]. A first round of scaffolding in *V. latastei* was performed using Bionano data with Bionano Solve v3.6.1 (Bionano Genomics), and a second round using SALSA2 v2.2[12] with parameters "-e GATC,GANTC -m yes". A final gap filling step was performed using RagTag v2.0.1[13] and another draft assembly of *V. latastei* from ONT data. This ONT raw data was filtered with Q>7 and minimum length of one kbp, assembled with Flye v2.8.3[14] with two polishing iterations, polished again with HyPo v2.4.1 [15] and scaffolded as the PacBio draft assembly, first with Bionano Solve and then with SALSA2. *Vipera aspis* Omni-C short read data was again quality-checked, and adapters were removed as described above. Three iterations of SALSA2 were implemented to scaffold and correct previous misassemblies. Finally, manual curation was performed on PretextView[16], with mapping threshold Q=10, to visualize the contact map, aided by coverage, gap, telomere, and repetitive regions graphs, following the rapid curation manual (https://gitlab.com/wtsi-grit/rapid-curation). The gap bed was obtained through the fasta-stats.py script (available in https://github.com/nextgenusfs/genome_scripts/tree/master). Telomere regions were identified using the conserved vertebrate telomeric sequence[17]. We soft-masked repetitive areas in the genomes with RED[18] using the redmask.py wrapper (https://github.com/nextgenusfs/redmask/blob/master/redmask.py). On the other hand, for scaffolding *V. seoanei*'s genome, we used the assembly of the closely related *V. ursinii* (rVipUrs1.1, PRJEB57183), with the same chromosome number[19], as reference input for RagTag. Finally, potential contamination in the assemblies was assessed with BlobToolsKit[20]. Two small scaffold/contigs were removed in the case of *V. seoanei* for extremely high coverage and the detection of mitochondrial genes, resulting in all genomic information contained in 18 scaffolds, matching the expected number of chromosomes[21]. Genome assemblies were depicted as snail plots with BlobToolsKit (Fig. S1-S3), and their contiguity and completeness were assessed with gfastats[22] and BUSCO (Vertebrate databases[23]), respectively.

**Genome annotation**

We annotated our three reference genomes by combining four annotation pipelines: 1) First we used GeMoMa v1.9[24], which is based on the conservation of intron positions in related organisms. As input for GeMoMa, we downloaded the annotated reference genomes of *Crotalus tigris* (ASM1654583v1; NCBI BioSample ID:

SAMN12497667; Margres et al., 2021), *C. adamanteus* (Cadam_11369; NCBI BioSample ID: SAMN16378231; Hogan et al., 2021), *Naja naja* (Nana_v5; NCBI BioSample ID: SAMN11155092), *Ophiophagus hannah* (OphHan1.0; NCBI BioSample ID: SAMN02439592; Vonk et al., 2013) and *Pantherophis guttatus* (UNIGE_PanGut_3.0; NCBI BioSample ID: SAMN12106767; Ullate-Agote et al., 2014). 2) Based on *de novo* sequenced full-length transcriptomes of different tissues. Subreads from Iso-Seq PacBio were first converted into circular consensus sequences (CCS) with the ccs function from pbccs v6.4 (Pacific Biosciences of California, Inc), followed by demultiplexing and primer removal with lima v2.7.1 (Pacific Biosciences of California, Inc). Non-chimeric full-length (NCFL) transcripts were then obtained by removing concatemers with isoseq3 v3.8.3 (Pacific Biosciences of California, Inc) and trimming poly-A tails with the python script tama_flnc_polya_cleanup.py (https://github.com/GenomeRIK/tama/wiki) mapped to each reference genome assembly with the splice-aware mapper pbmm2 v1.10 (Pacific Biosciences of California, Inc). The resulting mapped reads were collapsed into unique isoforms with isoseq3 v3.8.3 with the --do-not-collapse-extra-5exons flag. Collapsed unique isoforms were then used to identify protein-coding regions with GeneMarkS-T[29]. We employed transcripts from all the six libraries to annotated each of the three genomes to account for the maximum number of putative genes, even if they are not expressed in a particular species. 3) Based on RNA-Seq evidence from different tissues within each species, following BRAKER1 pipeline[30]. We quality-checked RNA-Seq Illumina reads with fastQC, and trimmed poly-A and poly-G tails, remove adapters and low-quality bases with fastp. Filtered reads were mapped to the respective reference using the splice-aware mapper STAR v2.7.10b[31] and then used as input for BRAKER1, which creates gene sets with GeneMark-ET to train Augustus. 4) Based on external protein databases following BRAKER2 pipeline[32]. We ran BRAKER2 twice per species, using the ortholog database Sauropsida OrthoDB v10[33] and all Sauropsida toxin isoforms from the Animal Toxin Annotation Project (https://www.uniprot.org/help/Toxins) as input to creates gene sets with GeneMark-EP+ to train Augustus. Finally, we integrated the different annotation files with TSEBRA[34], keeping all predictions from GeMoMa and GeneMarkS-T. BUSCO scores and general metrics from the resulting annotation files for each of the three species were calculated (Table S2).

**Macrosynteny analysis**

Chromosomal rearrangements were explored with MCscan[35] in between the three *Vipera* assemblies: *V. latastei*, *V. aspis* and *V. seoanei*, the latter representing the general chromosomal organization of the *berus* group, as its scaffolding was done using the closely related *V. ursinii* as reference. As outgroup we used the reference genome of *Crotalus viridis* (UTA_CroVir_3.0; NCBI BioSample ID: SAMN07738522; Pasquesi et al., 2018) to represent the sister subfamily Crotalinae. Protein sequences from each viperid species were extracted using AGAT v0.8.0[37] and pairwise aligned using LAST[38] with the JCVI python module[39]. Initial alignments between species were used to identify the Z chromosomes as well as chromosomes assembled in the reverse complement, which were corrected in some of the assemblies by reverse complementing the fasta sequence with SAMtools faidx v0.7.2.2[40]. Afterwards, GeMoMa was used to generate a draft annotation of the new genomes with reversed sequences, using as reference the original reference genome of each respective species. Then, we re-ran the MCscan pipeline to generate final re-oriented synteny plots.

**Processing of whole genome sequencing**

Raw Illumina data of all WGS samples (n=94) was quality-checked with fastQC and processed with fastp, filtering out poly-X and poly-A tails and bases with Q<30. Filtered reads were mapped to the reference genome of *V. latastei* (rVipLat1) with BWA, and then filtered by mapping quality with SAMtools (-q 30 -F 260). PCR duplicates were marked and removed with PicardTools v3.0.0 (https://broadinstitute.github.io/picard/). The individual read mean coverage was determined with SAMtools depth function. At this point, data processing was split into a low- and a high-coverage dataset. On the one hand, for the former one, samples with coverages higher than 10x were downsampled with SAMtools to 7x, which was the mean coverage among all the low-coverage samples. On the other one, high-coverage samples were kept at their original depth or downsampled to 45x in the case of the Illumina resequencing data from the three reference genomes to be part of the high-coverage dataset. Afterwards, we used GATK v4.1.3[41] to proceed with the SNP-calling in both datasets, through the functions HaplotypeCaller, CombineGVCFs and GenotypeGVCFs. The resulting datasets of genotype callings underwent then a stringent filtering pipeline. Density plots of six highly informative annotations from GATK tools were visualized to set optimized thresholds for each dataset: i.e. Quality by Depth (QD), Fisher Strand bias (FS), Strand Odds Ratio (SOR), the root mean square Mapping Quality over all the reads at the site (MQ), the u-based z-approximation from the Rank Sum test for mapping qualities (MQRankSum) and the u-based z-approximation from the Rank Sum test for site position within reads (ReadPosRankSum). For the high-coverage dataset, calls were kept only if QD > 7, FS < 70, SOR < 4, MQ > 55, -1.5 < MQRankSum < 1.5, -5 < ReadPosRankSum < 5, whereas for the low-coverage dataset: QD > 5, FS < 50, SOR < 4, MQ > 55, -1.5 < MQRankSum < 1.5, -3 < ReadPosRankSum < 3. Indels were normalized and SNPs within 10 bp to any indel were as well discarded. Furthermore, we excluded SNPs within repetitive regions previously inferred with RED, and sites which mapping was prone to be ambiguous following the SNPable pipeline (http://lh3lh3.users.sourceforge.net/snpable.shtml). These regions were inferred by splitting the reference genome of *V. latastei* into overlapping fragments of 100 bp (overlapping by 99 bp), mapping those regions back to the reference, and estimating the number of fragments that correctly mapped at each site. For those sites with a mapping rate below 95%, calls were discarded. Several datasets were split from both the high- and low-coverage branches of the data processing into more specific datasets to accommodate different purposes before further filtering (Table S12).

**Genomic and mitochondrial phylogenies**

Genomic species tree inferences of the studied European *Vipera* were performed using SNAPP[42] implemented on BEAST v2.6.4[43]. We produced two different datasets of low-coverage autosomal bi-allelic unlinked SNPs and 0% missing data with VCFtools[44]. The first one, a species dataset (i.e., *snapp_species* dataset, Table S12, samples in Table S14) comprising one representative of *V. latastei*, *V. monticola*, *V. aspis*, *V. ammodytes*, *V. berus*, *V. seoanei*, *V. ursinii* and *Crotalus viridis* as outgroup (Illumina raw data GenBank accession number SRA: SRR10593867; Schield et al., 2019). For the second dataset, we included representatives from all putative subspecies (i.e., *snapp_subspecies* dataset, see Tables S11,13) from the above-mentioned species plus another one from the Western lineage of *V. seoanei* (see Admixture results). We built two SNAPP time-calibrated species trees following Stange et al. [46]. The species dataset was LD-pruned by 0.5-kbp windows, yielding 1,225,190 uSNPs, whereas the subspecies dataset by 10-kbp windows to reduced computational demands, yielding 126,083 uSNPs (see "Population genomics" section below for LD-pruning thresholds). The vcf files of both datasets were converted to xml format by using the script *snapp_preb.rb* (available at https://github.com/mmatschiner/snapp_prep). First, we time-calibrated the species phylogeny with the divergence between Viperinae and Crotalinae, set with a lognormal distribution in 38.61±4.85 mya following a fossil-calibrated phylogeny[47]. For the subspecies phylogeny without *Crotalus*, we opted for dating the diversification of the crown *berus* group, which appears to be the only one whose topology is consistent across the literature and with our own results at species level (see SNAPP results, Fig. S4), with a lognormal distribution in 5.97±0.75 mya[47]. We carried out three different runs of each of the two SNAPP analyses for 20 million iterations each, sampling every 5,000 generations. Stationarity (ESS>200) and convergence between runs from the same dataset was checked with Tracer v1.7.2[48]. Converging log and tree files were combined with logCombiner v1.10 discarding 10% as burn-in. The cloudograms were visualized with DensiTree v.2.6.4[49], and TreeAnnotator v2.6.3 was used to construct the maximum clade credibility tree and annotate nodes with their corresponding posterior probabilities.

On the other hand, we produced a mitochondrial phylogeny to account for potential mito-nuclear discordance, using the whole mitogenomes of the same samples as the *snapp_subspecies* dataset (Table S14). From the adapter-trimmed Illumina reads of those samples, we extracted and assembled 15 mitogenomes with GetOrganelle v1.7.7.0[50], using the published mitochondrial assembly of *V. berus* (GenBank accession number: MF945570.1[51]) as reference and k-values: 21, 55, 85, 115. Additionally, we downloaded the mitogenome of *Crotalus horridus* (GenBank accession number: HM641837.1) as outgroup. The 16 mitogenome assemblies were annotated with MitoFinder[52] using *C. adamanteus* as reference (GenBank accession number: NC_041524.1). We aligned the thirteen protein-coding genes present in the mitogenomes with MEGA11[53], using the MUSCLE algorithm[54] and manually inspected for stop codons. We used only a 714-bp fragment of the gen ND4 to avoid stop codons. Single-gen alignments were concatenated with Geneious Prime v2023.2.1, yielding a 10,695-bp alignment. The best partition scheme was determined with PartitionFinder v2.1.1[55] using the *greedy* algorithm, constraining results to "beast" models, specifying AICc for model selection, linked branch lengths and potential partitions for each gen and each codon position within a gen in the alignment. It arranged the data into 18 partitions (Table S15). The model for partition 7, K81UF+I, was replaced by GTR+I instead in BEAUTi. We linked a relaxed clock model with frequencies sampled from a lognormal distribution to all partitions, but a relative substitution rate was estimated for each partition and trees among partitions were linked. Similar to the genomic phylogeny built on SNAPP, we calibrated the mitochondrial tree both with the Viperinae-Crotalinae split and the diversification of the *berus* group, with the same dates stated above. Applying such partition scheme and priors, we ran BEAST v2.7.4 three independent times for 50 million iterations each, sampling every 1,000 generations. We checked for the stationarity and convergence of the three runs in Tracer and used TreeAnnotator to construct the maximum clade credibility tree. The resulting mitochondrial phylogeny was overlapped to visualize mito-nuclear discordance with the subspecies cloudogram of SNAPP.

**Introgression analyses**

On the grounds of exploring introgression and geneflow events among the *Vipera* lineages, while considering ILS, we calculated ABBA-BABA tests, and their related estimate of admixture fraction, *f4-ratio*, with the function Dtrios implemented in Dsuite v0.5 r53[56]. For that purpose, we filtered out from the low-coverage dataset individual genotypes with a read depth below four or above 150, or a quality score below 30, non-biallelic alleles, SNPs with more than 10% missingness, singletons (adjusting a different minimum allele frequency threshold per dataset) and invariant sites and only kept SNPs from the autosomes. As an outgroup was needed to test the maximum trio combinations, we included *Crotalus viridis* in the analysis. On the other hand, from this dataset we excluded *V. latastei x aspis* putative hybrids, as well as individuals resulting from secondary contacts between intraspecific lineages, as depicted by Admixture (see Admixture results), yielding finally 86 individuals and 12,645,411 SNPs (*linked_SNPs* dataset, Table S12). We assumed the genomic species tree inferred by SNAPP to be the true phylogeny, and used all the taxa and lineages supported by that tree but *V. monticola* sspp., which were split in North and South lineages instead due to incongruent relationships among subspecies (see Results). Dtrios was ran with the "--ABBAclustering" option to avoid false signals of introgression due to substitution rate variation among branches, which can lead to "ABBA" homoplasies[57]. Then, for a better interpretation of the resulting $f_4$*-ratio* values, and to correctly take correlated excess allele-sharing into account, we calculated *f-branch* statistics[58] implemented as well in Dsuite. Per default, all $f_4$*-ratio* values are set to zero when the *p-value* of the associated D statistic for that trio is > 0.01[56], but we chose to equally purge out Z scores |Z| < 3 and, moreover, the ABBA-clustering tests' p-values > 0.01 (Table S3). These tests calculate two sets of *p-values* for each trio with different sensitivity and robustness (Tables S4-S5). We plotted the results following both types of thresholds (sensitive and robust) with the Dsuite script dtools.py. Finally, to investigate the direction of the detected introgression events, we ran TreeMix v1.13, which uses SNP frequency data to evaluate population splitting, drift and directional introgression[59]. We used a dataset of 15 individuals representing all subspecies or lineages in the studied *Vipera* spp. with 0.5-kb thinning and 0% missingness, yielding a total of 1,095,590 bi-allelic autosomal uSNPs (*treemix_subspecies* dataset, Tables S11-S13). We analyzed the data in blocks of 20, 100 and 500 SNPs to account for non-independence of the data, although we already used LD-pruned SNPs. Migration edges were calculated for values of *m* ranging from 1-10 and the options "-noss" and "-global" activated. The *V. latastei-monticola* complex was used to root the resulting trees. Finally, the optimum number of migration edges was calculated with the R package OptM v0.1.6[60] following the Evanno method[61]. The highest $\Delta m$ is supposed to be reached with the optimum number of edges that can best explain the genetic variance from complex histories.

To explore the introgression landscape of the strongest identified introgression event (i.e., *V. latastei* to *V. seoanei*, see Results), we investigated the local window topologies across the genome. We first phased haplotypes of a high coverage dataset of all studied species (*high_coverage+out* dataset, see Table S12) with Beagle v5.2[62]. Then we inferred Neighbour-Joining local trees in non-overlapping 25-kbp windows with PhyML v3.3[63] and the script phyml_sliding_windows.py (available in https://github.com/simonhmartin/genomics_general), with the requirement of bearing at least 50 SNPs per window. For the resulting 27,217 local trees, complete topology weightings were computed by TWISST[64], using two haploid sequences per species at 45x coverage. Topologies in which *V. latastei-monticola* clustered with *V. seoanei* were grouped and highlighted across the genome.

**Demographic inference**

We used Pairwise Sequentially Markovian Coalescent (PSMC) models[65] to infer the demographic histories of the eight main studied lineages, namely *V. latastei*, *V. monticola*, Western and Eastern *V. aspis*, *V. ammodytes*, *V. ursinii*, *V. seoanei* and *V. berus*, at high coverage. The three Iberian samples come from the resequencing data used for each of the three reference genomes, which was downsampled to 45x to equal the coverage of the other samples and filtered for low mapping (<30) and base quality (<30). Diploid consensus sequences for autosomal data of the eight high-coverage genomes were obtained by the pileup command of SAMtools. Minimum and maximum depths were set at half and double the average coverage of each sample, respectively. A rate of 2.4e-9 substitutions/site/year[66] and a 6-year generation period, as estimated for *V. aspis*[67], were used. Other parameters were set following previous studies on Viperidae[68] and ten bootstraps were calculated for each genome. Results were plotted in base R, dividing Eurosiberian (*V. seoanei*, *V. berus* and *V. ursinii*) and Mediterranean species (*V. latastei*, *V. monticola*, Western and Eastern *V. aspis* and *V. ammodytes*) in two different plots to analyze common climatic oscillations due to similar affinities.

**Population genomics**

In order to explore population genomics of the genus *Vipera* in the Western Mediterranean, we created a dataset including only samples from *V. seoanei*, *V. aspis, V. latastei* and *V. monticola* (*westmed* dataset, see Table S12); apart from four species-specific datasets for the above mentioned species and another one for the joint *V. latastei-monticola* complex. Two putatively *V. latastei x aspis* hybrids were included only in the *westmed* dataset. For all these resulting low-coverage datasets, we filtered out individual genotypes with a read depth below four or above 150, or a quality score below 30, non-biallelic alleles, SNPs with more than 10% missingness, singletons (adjusting a different minimum allele frequency threshold per dataset) and invariant sites, and only kept SNPs from the autosomes with VCFtools. Then, to work with unlinked SNPs (uSNPs), we calculated Linkage Disequilibrium (LD) decay with PopLDdecay[69] in *V. seoanei*. As LD falls below 20% of correlation in between sites 0.5 kbp apart (Fig. S34), we pruned the datasets to kept one SNP per 0.5 kbp with VCFtools (--thin).

We first explore population structure of the four above-mentioned species and phylogenomic relationships between them by performing Principal Component Analyses (PCA) on the following datasets: *westmed*, *latastei-monticola*, *latastei*, *monticola*, *seoanei* and *aspis* (Table S12). The PCAs were implemented in Plink v1.90b6.21[70]. Secondly, to delve into population structure and secondary contacts in between lineages, we ran Admixture v1.3.0[71] on the *westmed* dataset, calculating 10 cross-validation errors with 10 replicates. K values were set to 12, coinciding with the number of putative subspecies/lineages comprised within the four species. Optimal K values were chosen based on CV scores (see Table S16), the plateau profile of those values and the likelihood of the results in biological terms. Admixture results were visualized and plotted R v4.3.1[72], and the most likely K was depicted with georeferenced pie charts using QGIS[73].

Finally, we explored genomic Isolation-by-distance (IBD) patterns within *V. latastei* and *V. seoanei* following Hausdorf and Hennig.[74]. From the *latastei* and *seoanei* datasets, we excluded all admixed individuals (in regard to the chosen Admixture results for the *westmed* dataset) to calculate genomic chord distances matrices among all individuals of each species with genodive v3.06[75]. Two analogous matrices of geographic distances among samples were calculated as well with genodive. Geographic distances were transformed as $f_{ij} = log_{10}(d_{ij} + c)$, being $d_{ij}$ the geographical distances in meters and c the 0.25-quantile of such raw distances[74]. For these two species, we used the R package prabclus[76] to perform a Jackknife-based test for the equality of two independent within-lineage regressions (i.e. within the Western and Eastern lineages of *V. latastei* and *V. seoanei*, independently) to confirm that IBD develops alike within each lineage. Under those circumstances, within-lineage regressions of the same species can be merged into a single joint within-lineage regression. Secondly, either this joint regression or the independent specific within-lineage regressions previously calculated were compared to another regression over all comparisons within a species. The non-equality of them imply that the observed genomic differences cannot be explained by IBD alone. IBD regressions were plotted with ggplot2[77] in R.

#### Heterozygosity and Runs of Homozygosity

We calculated autosomal heterozygosity per individual from the unfiltered low-coverage dataset, following Mochales-Riaño et al.[78]. We generated 100-kbp non-overlapping sliding windows along the autosomes and took variant and invariant sites with high base quality (Q>30). Per window heterozygosity values, relative to the number of called sites, were calculated and then individual genome-wide heterozygosity means were obtained. For each species and the *V. latastei x aspis* putative hybrids, the individual mean or a species mean among all samples were plotted as a barplot with ggplot2. To geographically depict gradients in genomic diversity and identify possible barriers to geneflow, we created three raster surfaces in QGIS through inverse distance weighted interpolation of the mean individual heterozygosity values for *V. latastei-monticola*, *V. seoanei* and *V. aspis*. *Vipera latastei x aspis* hybrids were excluded from these interpolations.

Furthermore, we explored how different *Vipera* species are affected by Runs of Homozygosity (ROHs). First, we filtered the high-coverage dataset with different *Vipera* spp. from individual genotypes with a read depth below four or above 150, or a quality score below 30, non-biallelic alleles, and invariant sites and only kept SNPs from the autosomes. This dataset, with 10,754,630 SNPs was divided into several single-sample vcf files and purged out invariant sites again. ROHs were calculated then based on the density of those remaining heterozygous sites by the implemented hidden Markov model (HMM) in bcftools roh function[79], setting the allele frequency as 0.4 per default, being thus conservative when identifying ROHs[80]. We kept ROHs with a PhredScore above 70 and a minimum length of 100 kbps[78]. Finally, we divided the amount of ROHs in each high-coverage genome by size, as different-sized ROHs arise from different causes[81], in the following categories: short (0.1-0.5 Mbp), medium (0.5-1 Mbp) and long (>1 Mbp); and plotted the result with ggplot2, showing the amount of ROHs as a percentage of the total autosomal genome (*V. latastei* as reference).

#### Climatic niche overlap

With the purpose of investigating the role of adaptative introgression in between the ecologically divergent *V. latastei* and *V. seoanei*, we performed pairwise niche overlap tests between the major lineages identified in both species (see Admixture results). We included 1,105 and 508 occurrence records for *V. latastei* and *V. seoanei*, respectively, with a spatial resolution of 5 arc minutes, throughout the entire distribution of the two species, extracted from[82–84]. For these locations, we obtained a set of 19 environmental variables from WorldClim v2.1[85]. To correctly assign the records into the genomically-identified lineages (i.e. Eastern or Western *V. seoanei* and Eastern or Western *V. latastei*), we interpolated the genomic affinities inferred from Admixture results using ordinary kriging as interpolator method in ArcGis v.10.5[86]. We separated intraspecific lineages in both species, calculating the contact zones between them and delimitating the records for each cluster with a genomic affinity higher than 0.73 for each lineage. Afterwards, based again on Admixture results (K=7 for *westmed* dataset), we identified isolated populations of the Western *V. seoanei vs* Western *V. latastei* contact zone as putatively admixed: the nuclei of Montalegre and Eastern Galicia, south to Sil River for *V. seoanei* and the nuclei of Gerês, Montesinho-Sanabria and northern coastal Porto for *V. latastei* (admixed individuals for those five nuclei shown in Fig. A2). Later, we subset six different record datasets with their respective environmental variables: "Western *V. seoanei*" (n=362 records; including all populations with or without admixture from *V. lastatei*), "Western *V. seoanei* no-admix" (n=326, excluding records from the cited two admixed nuclei), "Eastern *V. seoanei*" (n=146), "Western *V. latastei*" (n=488, with or without *V. seoanei* admixture), "Western *V. latastei* no-admix" (n=412, excluding records from the three cited nuclei) and "Eastern *V. lastatei*" (n=617). Climatic data was summarized by a PCA into three Principal Components (factor loadings in Table S7) and, following a three-dimensional hypervolume approach[87], we used those PCs to describe the environmental niche of the subgroups. Analyses were performed in R with the hypervolume package[88], using a Silverman bandwidth estimator and a set of 999 random points to sample the kernel density to build the hypervolumes. To quantify the climatic niche overlap the Sørenson index (K) and the Overlap Index (OI) were used[87–89]. These statistics quantify predicted niche similarity, and they both range from no overlap (0) to identical niche (1). The OI index relates the observed and maximum values of K and is preferred when niches present different sizes[89,90]. We statistically tested the significance of the overlap comparisons with a randomization procedure[89]. By comparing the produced K and OI indices, we tested if observed overlaps statistically differed from overlaps obtained from random lineage-blind occurrences. This procedure was performed 999 times to generate a null distribution and to evaluate the significance of each overlap. Depending on how much the niche overlap in between Western *V. seoanei* and Western *V. latastei* increases when adding admixed populations, this could be a line of evidence supporting the adaptive introgression, i.e. both species benefit from the introgression with a distant and ecologically differing species, as these events allow the resulting admixed individuals to increase their range beyond the potential range of the species. Once lineage-pairwise niche overlap was tested through hypervolumes, we plotted as well 2D PCAs in between sympatric species/lineages, highlighting putatively admixed individuals to better visualized if they inhabit areas beyond the common niche of the species in absence of recent introgression.

**Differential Gene Expression analysis**

RNA-Seq data, already filtered and adapter-trimmed (see "Genome annotation" section), was again mapped to the annotated reference genome of *V. latastei* with STAR v2.7.10b. Mapping was quality-checked with PicardTools CollectRnaSeqMetrics and then used the sorted bam files to count gene hits with the count function of HTSeq v2.0.4[91], in which the number of reads are taken as an indicator of gene expression. We used the mode "-m intersection-nonempty" to be as stringent as possible, finally generating a gene expression matrix of raw read counts, which was then normalized by both library size and composition using the median-of-ratios algorithm of DESeq2[92] in R. After normalization, genes with lower than ten counts in total were filtered out, keeping 17,811 genes, and data was transformed with the regularized logarithm of DESeq2 (rlog function). We calculated the variance of each gene among the 19 samples and selected the top 1,000 genes with the highest variance to perform an unsupervised hierarchical clustering analysis, plotted with the R package ComplexHeatmap[93]. In this plot, we checked whether venom gland samples and other duplicated organs (i.e., kidneys, liver, pancreas) clustered together. Gene expression was standardized with Z-score scaling. Finally, we tested for significantly upregulated genes (log2 fold-change > 2) in the venom glands *vs* all other tissues with DESeq2, correcting for multiple tests with Benjamini and Hochberg adjusted *p-values* (FDR: 5%). The function of the putatively venom gland-upregulated genes was identified through protein BLAST[94].

**Venom decomplexation and initial characterization of *V. latastei* proteomes**

Crude lyophilized venom was dissolved in 0.05% trifluoroacetic acid (TFA) and 5% acetonitrile (ACN) to a final concentration of 10 mg/mL. Insoluble material was removed by centrifugation in an Eppendorf centrifuge at 13,000xg for 10 min at room temperature, and 600 µg were fractionated by RP-HPLC using an Agilent LC 1200 Infinity II High Pressure Gradient System equipped with a Teknokroma Europa C18 (25 cm x 5 mm, 5 µm particle size, 300 Å pore size) column and a DAD detector. The column was developed at a flow rate of 1.0 mL/min with a linear gradient of 0.1% TFA in MilliQ® water (solution A) and in acetonitrile (ACN, solution B), isocratic (5% B) for 5 min, followed by 5-25% B for 10 min, 25-45% B for 60 min, and 45-70% B for 10 min. The elution was monitored at 215 nm with a reference wavelength of 400 nm. Fractions were collected manually and dried in a vacuum centrifuge (Savant™, ThermoFisher Scientific). Molecular masses of the purified proteins were estimated from Coomassie Brilliant Blue G-250 SDS-10% and 15% polyacrilamide gels, run under non-reduced and reduced conditions, respectively. Molecular masses were also measured by electrospray ionization (ESI) mass spectrometry (MS). To this end, the proteins eluted in the different RP-HPLC fractions were separated by nano-Acquity UltraPerformance LC® (UPLC®) using BEH130 C18 (100µm x 100mm, 1.7 µm particle size) column in-line with a Waters SYNAPT G2 High Definition Mass Spectrometry System. The flow rate was set to 0.6 µL/min and the column was developed with a linear gradient of 0.1% formic acid in MilliQ® (solution A) and ACN (solution B), isocratically 1% B for 1 min, followed by 1-12% B for 1min, 12-40% B for 15min, 40-85% B for 2min. Monoisotopic and isotope-averaged molecular masses were calculated by manually deconvolution of the isotope-resolved multiply-charged MS1 mass spectra.

**Identification of the venom proteins eluted in the RP-HPLC fractions of the *V. latastei* venoms**

Protein bands found in each RP-HPL fraction were excised from Coomassie Brilliant Blue-stained SDS-PAGE and the pieces were subject to automated in-gel disulphide bond reduction (10 mM dithiothreitol, 30 min at 65 ºC), cysteine alkylation (50 mM iodoacetamide, 2h in the dark at room temperature), and overnight digestion with sequencing-grade trypsin (66 ng/µL in 25 mM ammonium bicarbonate, 10% ACN; 0.25 µg/sample), using a Genomics Solution ProGest™ Protein Digestion Workstation. Tryptic digests were dried in a vacuum centrifuge (SPD SpeedVac®, ThermoSavant), redissolved in 14 µL of 5% ACN containing 0.1% formic acid, and 7 µL submitted to LC-MS/MS. Tryptic peptides were separated by nano-Acquity UltraPerformance LC® (UPLC®) in-line with a Waters SYNAPT G2 High Definition Mass Spectrometry System as above. For peptide ion fragmentation by collision-induced dissociation tandem mass spectrometry (CID-MS/MS), the electrospray ionization (ESI) source was operated in positive ion mode, and both singly and multiply charged ions were selected for CID-MS/MS at sample cone voltage of 28 V and source temperature of 100 °C. The UPLC eluate was continuously scanned from 300 to 1990 *m/z* in 1 s and peptide ion MS/MS analysis was performed over the range m/z 50–2000 with scan time of 0.6 s. Search parameters were: taxonomy: bony vertebrates; enzyme: trypsin (two-missed cleavage allowed); MS/MS mass tolerance was set to ±0.6 Da; carbamidomethyl cysteine and oxidation of methionine were selected as fixed and variable modifications, respectively.

All matched MS/MS data were manually checked. For missing/incomplete identifications, the MS/MS fragmentation spectra were interpreted manually (*de novo* sequencing), and amino acid sequence similarity searches were performed at https://blast.ncbi.nlm.nih.gov/Blast.cgi. against the non-redundant NCBI protein sequence database, using the default parameters of the BLASTP. Fragmentation spectra of peptides that yielded daughter ions diagnostic of the endogenous peptides BPP (*m/z* 116.1, y1 = P, and m/z 213.1, y2 = PP) and tripeptide inhibitors of metalloproteinases (SVMPi) at m/z 205.1 (y1 = W), m/z 112.1 (b1 = pyroglutamate, Z) and m/z 240.1 (b2 = Z(K/Q) or m/z 226.1 (b2 = ZN)[95,96] were also sequenced manually. The final proteomics-gathered proteomes were manually assigned to the subset of venom toxins annotated in the reference genome of *V. latastei*.

**Quantification of the venom proteomes**

The relative abundances of the chromatographic peaks obtained by reverse-phase HPLC fractionation of the crude venom were calculated as "% of total peptide bond concentration in the peak" by dividing the peak area by the total area of the chromatogram[97,98]. For chromatographic peaks containing single components (as judged by SDS-PAGE and/or MS1), this figure is a good estimate of the % by weight (g/100 g) of the pure venom component[99]. When more than one venom protein was present in a reverse-phase fraction, their proportions (% of total protein band area) were estimated by densitometry of Coomassie-stained SDS-polyacrylamide gels using MetaMorph® Image Analysis Software (Molecular Devices). Conversely, the relative abundances of different proteins contained in the same SDS-PAGE band were estimated based on the relative ion intensities of the three most abundant peptide ions associated with each protein by MS/MS analysis. The relative abundances of the protein families present in the venom were calculated as the ratio of the sum of the percentages of the individual proteins from the same toxin family to the total area of venom protein peaks in the reverse-phase chromatogram, and these figures were used to compile a compositional pie chart for the venom proteomes[97,98].

**Characterization of venom-encoding genes and venom selection analyses**

We used proteomic data (i.e. peptide sequences) to identifying the venom-encoding genes in each *Vipera* spp. reference genome. For *V. latastei*, we used the data produced in this study (Methods above), whereas for *V. seoanei* and *V. aspis* we used already published data[100,101]. We translated into protein sequences all annotated isoforms of the three genomes with AGAT and searched for exact matches from each peptide database with the translated isoforms of each corresponding species using custom bash scripts. For all matching genes (i.e. candidate venom-encoding genes), we kept the isoform with the highest number of hits against unique peptide sequences as representative for that gene. The identified the function of each protein sequence with BLASTP, manually discarding candidate venom-encoding genes matching non-toxic proteins. For candidate genes with more than one isoform with the same number of hits, we kept the one with the higher resemblance to known toxins (i.e. a higher e-value in BLASTP). We checked that results from proteomic evidence were similar to the ones based on differential gene expression for *V. latastei*. For the main toxin families (i.e. SVMPs, SVSPs, $PLA_2$s, CTLs, Kunitz peptides), we also checked that the chromosomal location of each gene cluster agreed between the three *Vipera* species (see Macrosynteny results) and the sister subfamily Crotalinae. Finally, once the chromosomal location of each of the above-mentioned toxin families was known, we conducted chromosome-wide blasts of all isoforms annotated in such chromosome against custom databases of Viperidae toxins. In that way, we were able to identify putative candidate genes that were not expressed in the sampled venoms/venom glands.

**Selection signals on venom-encoding genes**

Once the toxin-encoding genes were identified, we extracted the coding sequences (CDS) with AGAT and aligned with all copies belonging to each toxin family (n=11, families with at least 3 copies present in 2 species) using the MUSCLE algorithm implemented on MEGA 11. We trimmed the terminal stop codons and performed $d_N/d_S$ (ω) ratio tests in each family of toxins with BUSTED[102] and FEL[103] analyses implemented in Datamonkey 2[104]. BUSTED tests for episodic diversification, whereas FEL looks for pervasive diversifying and purifying selection. For FEL tests, *p-value* was left as default (0.1). Later, to estimate the relative ubiquity or pervasiveness of both types of selection in each family, we divided the number of sites under selection estimated with FEL tests by the number of codons in the alignment and the number of copies in the family, since the more copies or the more codons, the more likely is to find selection signals. This selection ubiquity values were normalized and used to explore the relationship between the ubiquity of diversifying and purifying selection with the number of copies within a family or its relative expression in the venom. The expression was measured in relative abundance of the family in the venom, taken from Giribaldi et al.[101]. The disintegrin expression percentage was added to SVMPs, as they derived from the same genes. Regressions were performed in base R and plotted with ggplot2.
