## Supplementary Figures and Tables for "Unveiling the evolutionary history of European vipers and their venoms from a multi-omic approach"

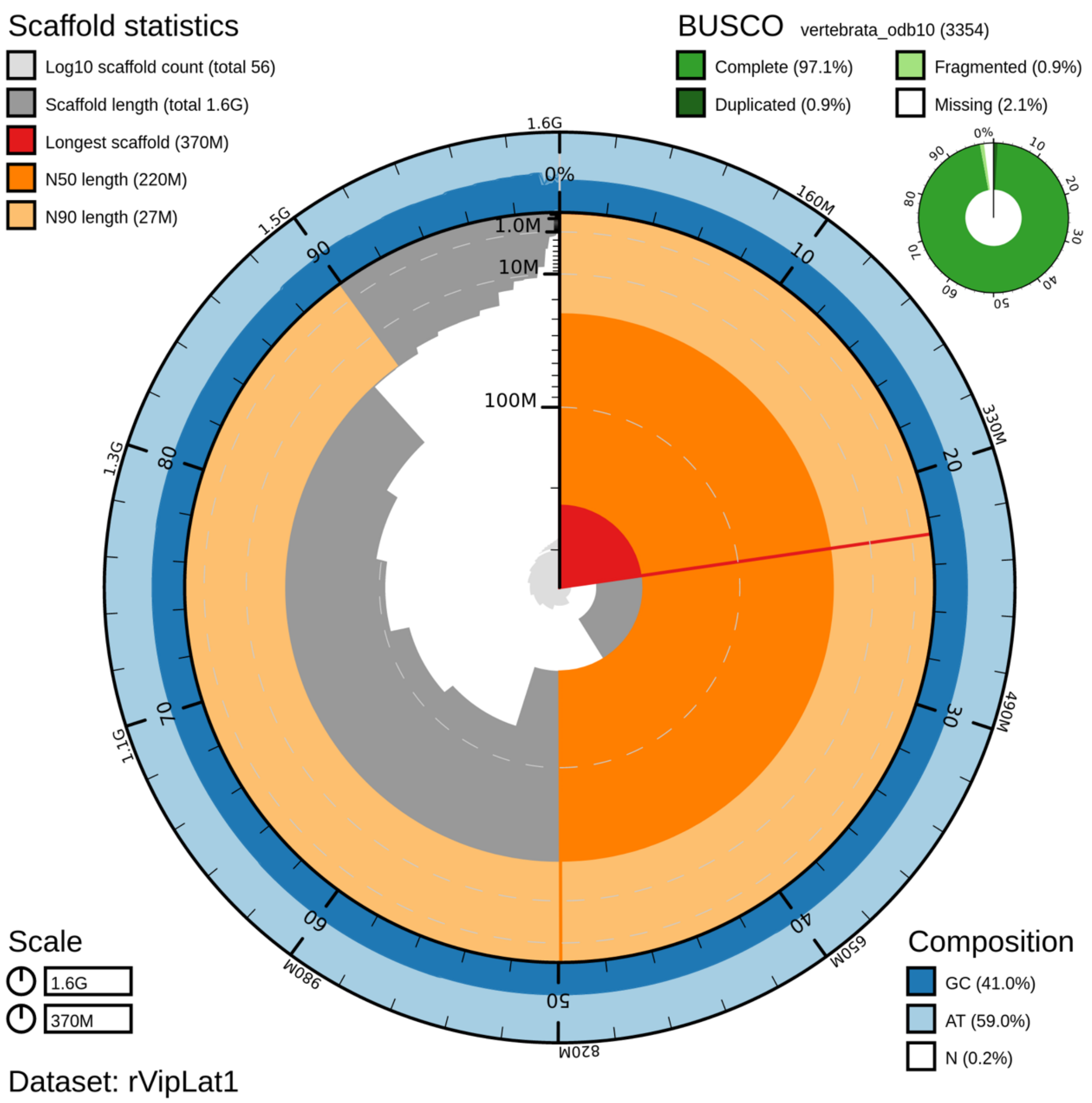


**Fig. S1. Snail plot of *Vipera latastei* reference genome assembly presented in this study (rVipLat1), summarizing some contiguity and completeness metrics.** M=Mega bases; G=Giga bases. The scaffolds contained in the assembly are shown ordered by length in the inner circle of the snail plot. The sequences within the N50 (220 Mb) and N90 (27 Mb) length marks are painted in dark or light orange, respectively. In the outer circle, the GC content of the sequences in represented in blue, whereas AT content in pale blue. Top left corner shows contiguity statistics like the longest scaffold (1.6 Gb) or Scaffold length (total 1.6 Gb) which indicates the assembly size. Top right corner represents the assembly genetic completeness using BUSCO scores with odb10 vertebrata database with 3354 genes, with 97.1% of the database genes found as Single-Copy Complete genes. Bottom left has a legend with the length in bases represented in the snail plot, the perimeter (1.6 Gb) and the radius (370 Mb, i.e., chromosome 1 length).

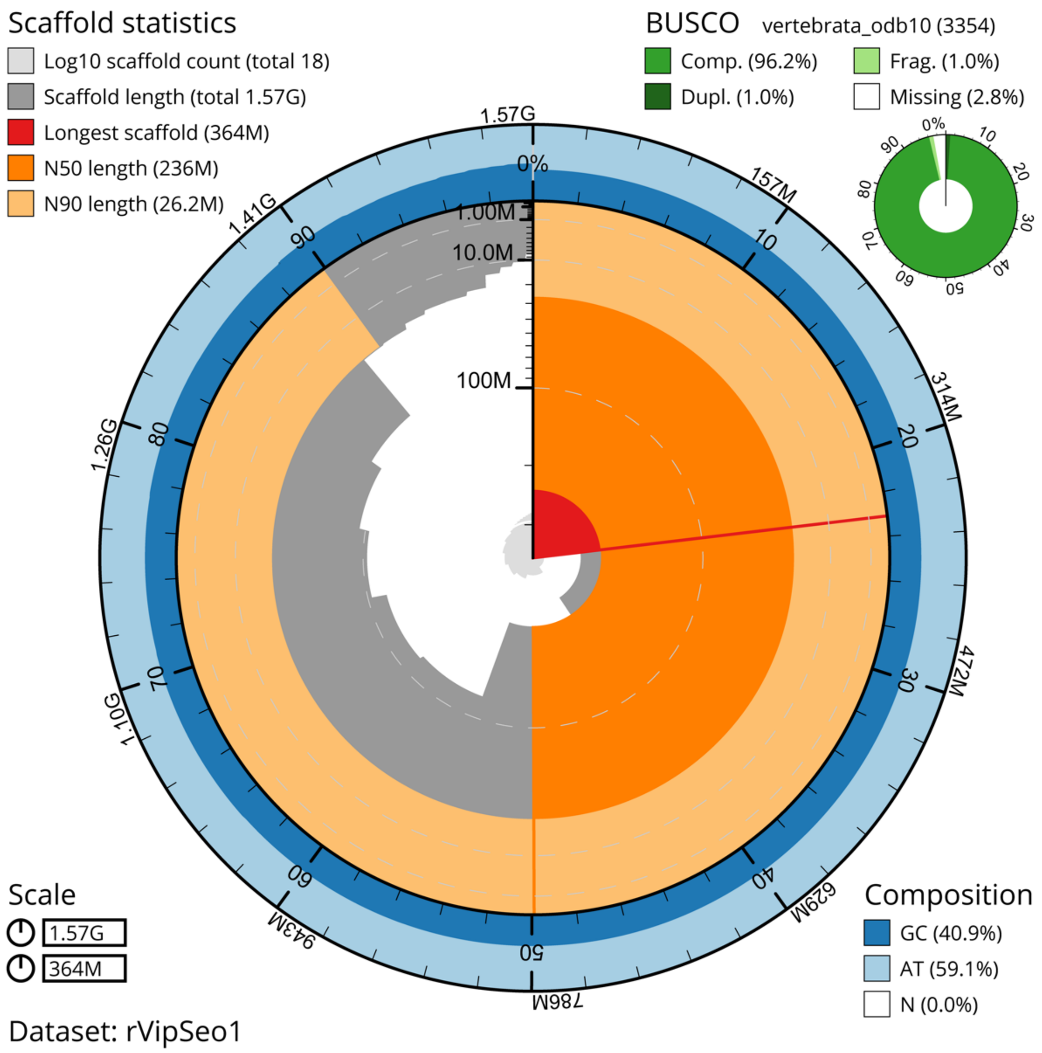


**Fig. S2. Snail plot of *Vipera seoanei* reference genome assembly presented in this study (rVipSeo1), summarizing some contiguity and completeness metrics.** M=Mega bases; G=Giga bases. The scaffolds contained in the assembly are shown ordered by length in the inner circle of the snail plot. The sequences within the N50 (236 Mb) and N90 (26.2 Mb) length marks are painted in dark or light orange, respectively. In the outer circle, the GC content of the sequences in represented in blue, whereas AT content in pale blue. Top left corner shows contiguity statistics like the longest scaffold (364 Mb) or Scaffold length (total 1.57 Gb) which indicates the assembly size. Top right corner represents the assembly genetic completeness using BUSCO scores with odb10 vertebrata database with 3354 genes, with 96.2% of the database genes found as Single-Copy Complete genes. Bottom left has a legend with the length in bases represented in the snail plot, the perimeter (1.57 Gb) and the radius (364 Mb, i.e., chromosome 1 length).

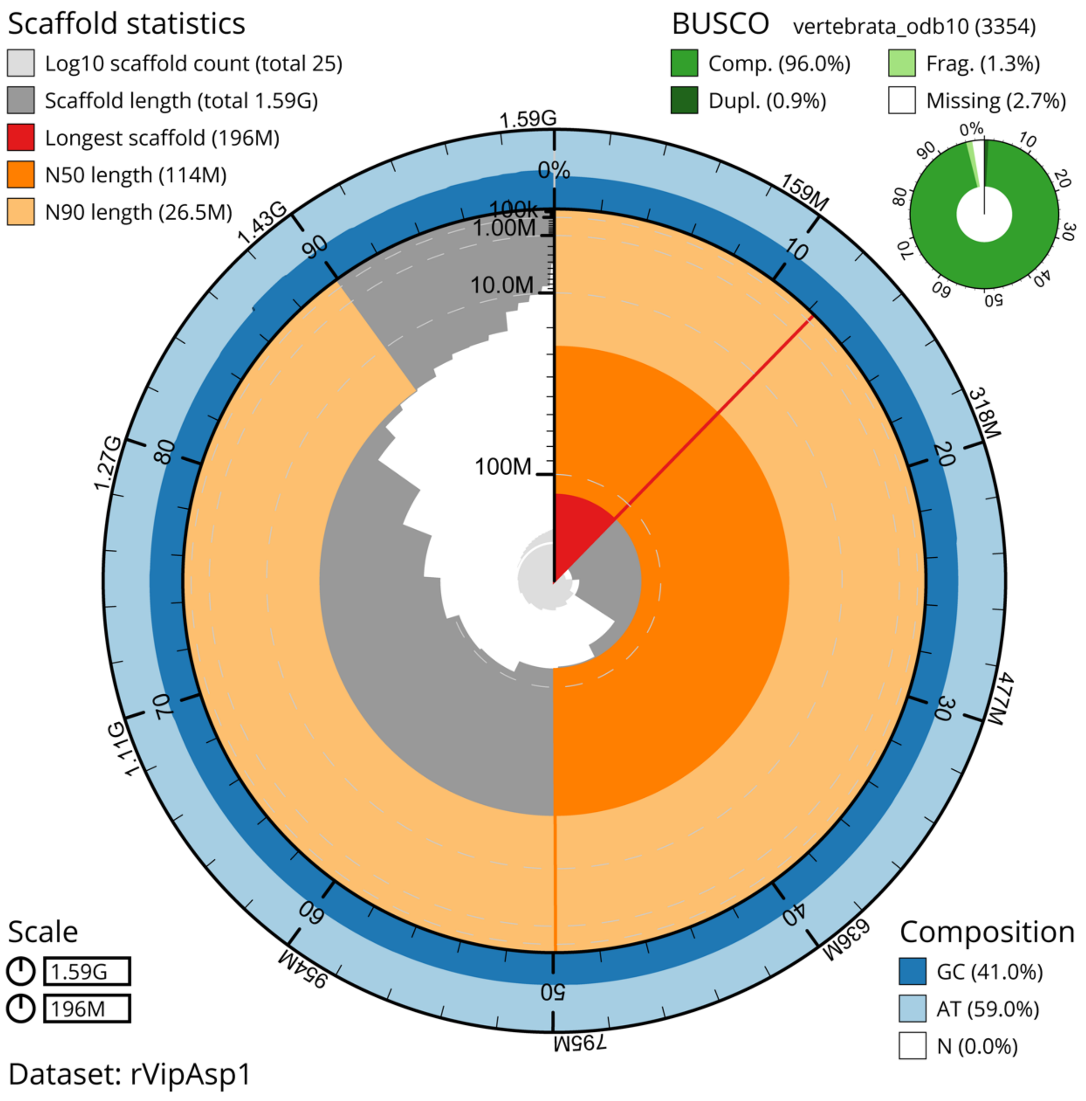


**Fig. S3. Snail plot of *Vipera aspis* reference genome assembly presented in this study (rVipAsp1), summarizing some contiguity and completeness metrics.** M=Mega bases; G=Giga bases. The scaffolds contained in the assembly are shown ordered by length in the inner circle of the snail plot. The sequences within the N50 (114 Mb) and N90 (26.5 Mb) length marks are painted in dark or light orange, respectively. In the outer circle, the GC content of the sequences in represented in blue, whereas AT content in pale blue. Top left corner shows contiguity statistics like the longest scaffold (196 Mb) or Scaffold length (total 1.59 Gb) which indicates the assembly size. Top right corner represents the assembly genetic completeness using BUSCO scores with odb10 vertebrata database with 3354 genes, with 96.0% of the database genes found as Single-Copy Complete genes. Bottom left has a legend with the length in bases represented in the snail plot, the perimeter (1.59 Gb) and the radius (196 Mb, i.e., chromosome 1 length).

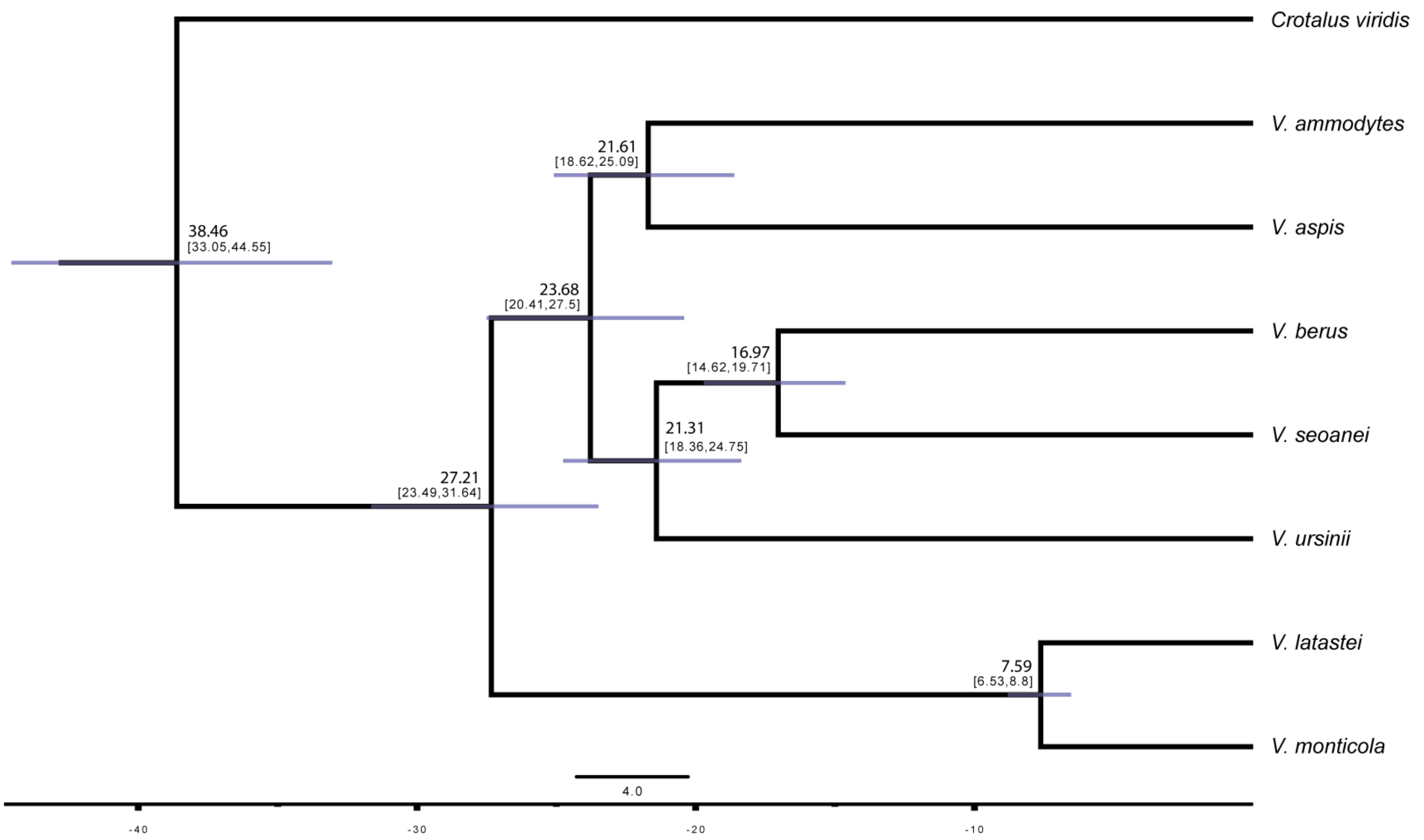


**Fig. S4. Time-calibrated genomic species tree inferred with SNAPP, using 1.23M uSNPs and eight species.** Each node is annotated with its median age and 95%CI between brackets. The posterior probabilities of all nodes are 1.

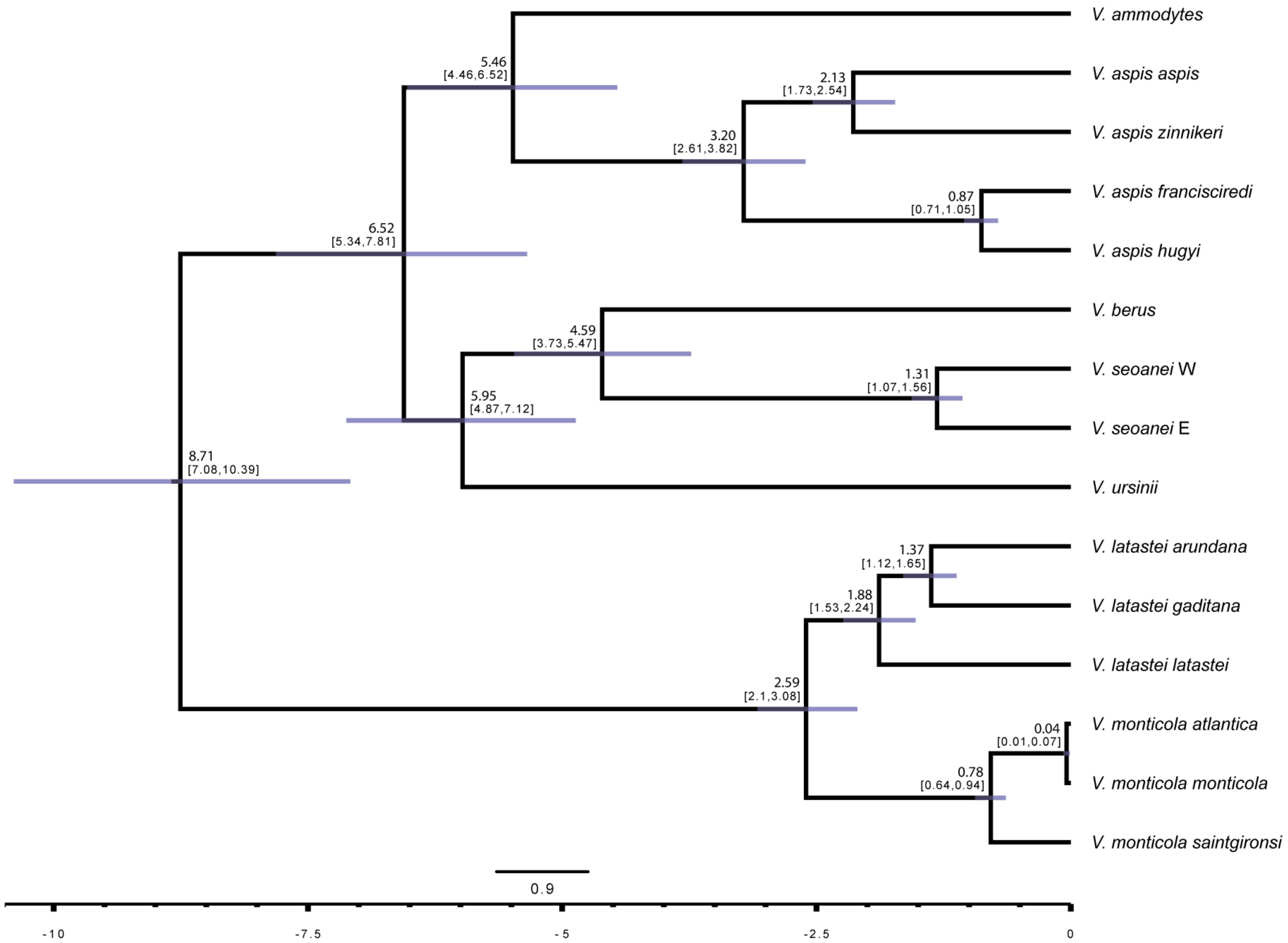


**Fig. S5. Time-calibrated genomic species tree inferred with SNAPP, using 126k uSNPs and 15 taxa.** Each node is annotated with its median age and 95%CI between brackets. The posterior probabilities of all nodes are 1.

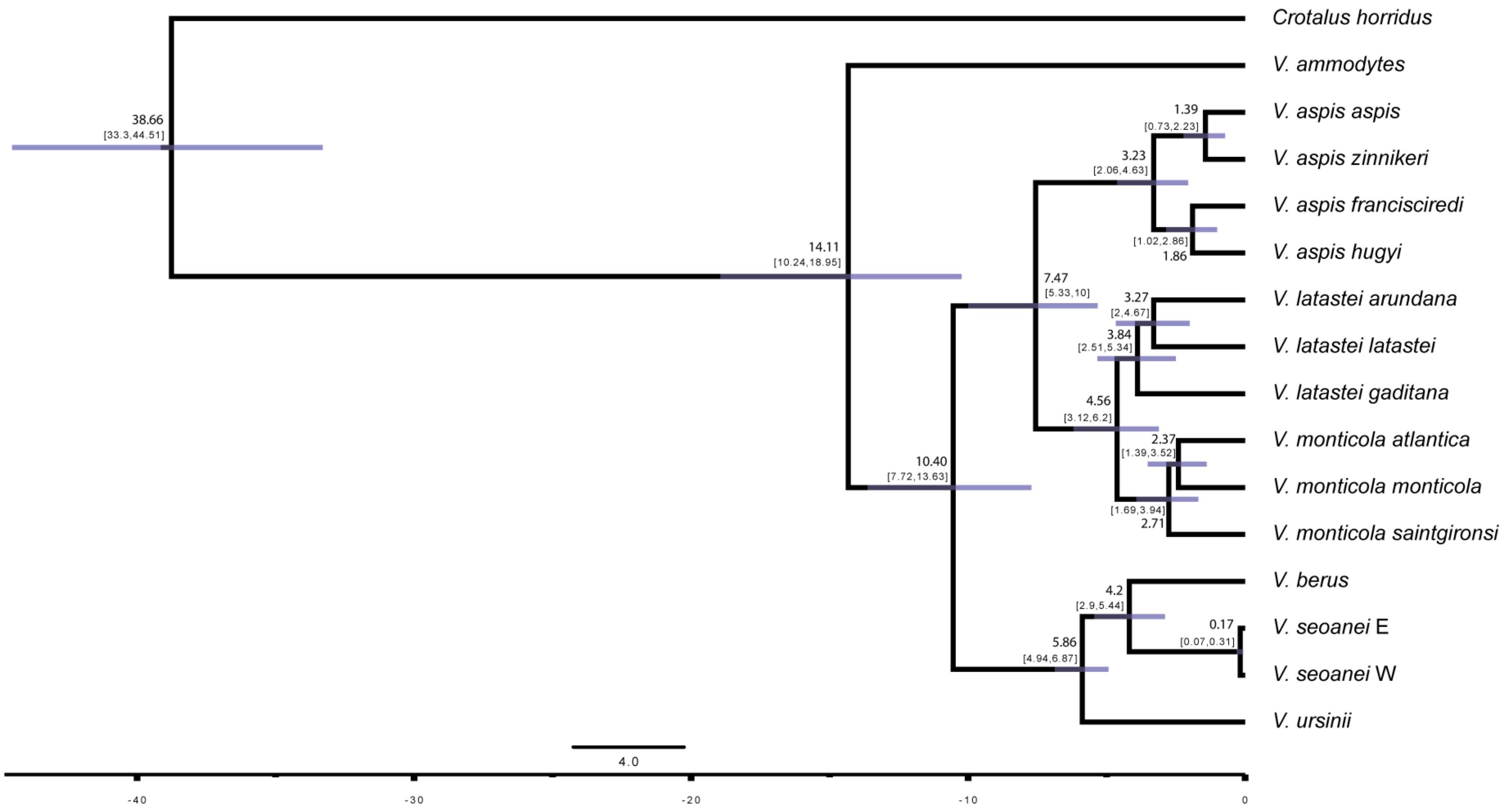


**Fig. S6. Time-calibrated mitochondrial tree inferred with BEAST2, using a 10.7-kbp alignment, 18 partitions and 13 protein coding genes from 15 taxa.** Each node is annotated with its median age and 95%CI between brackets. The posterior probabilities of all nodes are 1.

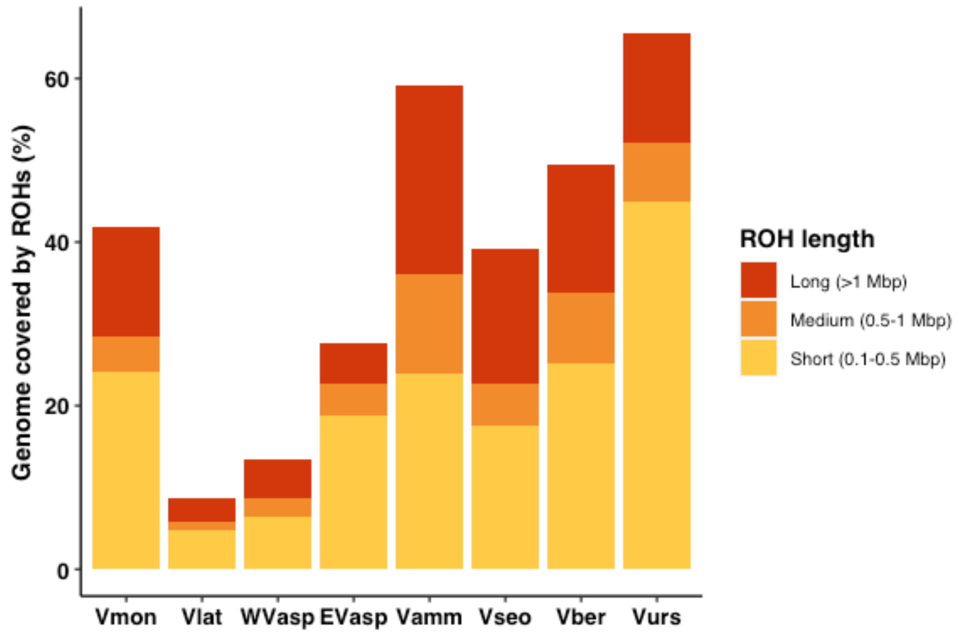


**Fig. S7. Percentage of autosomes covered by Runs of Homozygosity (ROHs).** Vmon stands for *V. monticola*, Vlat for *V. latastei*, WVasp for Western *V. aspis*, EVasp for Eastern *V. aspis*, Vamm for *V. ammodytes*, Vseo for *V. seoanei*, Vber for *V. berus* and Vurs for *V. ursinii*. Western *V. aspis* and *V. latastei* show the lowest percentage of their genomes covered by short ROHs, whereas *V. ursinii*, the highest.

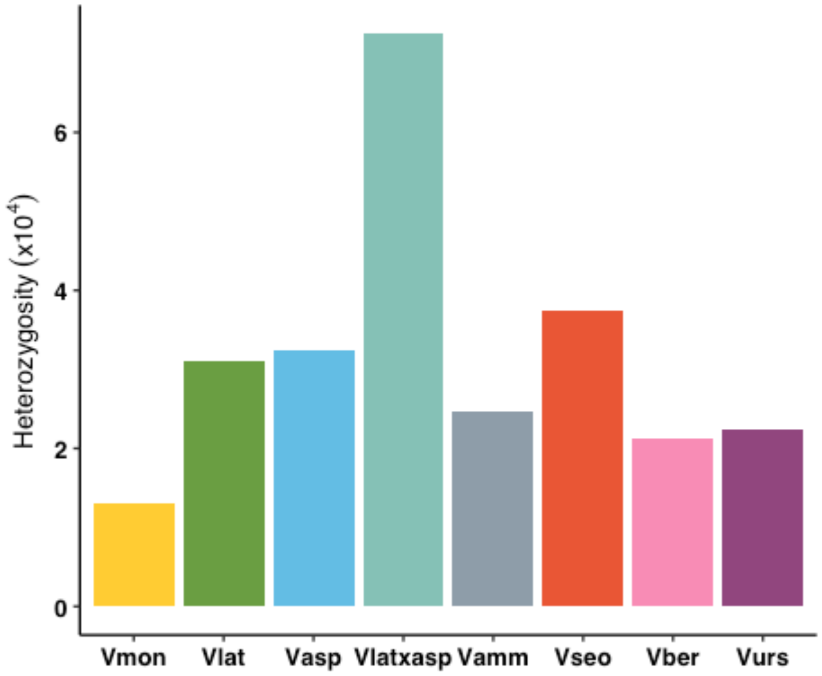


**Fig. S8. Mean genome-wide heterozygosity values per species.** Vmon stands for *V. monticola*, Vlat for *V. latastei*, Vasp for *V. aspis*, Vlatxasp for *V. latastei x aspis* F1 hybrids, Vamm for *V. ammodytes*, Vseo for *V. seoanei*, Vber for *V. berus* and Vurs for *V. ursinii*. *Vipera monticola* (n=8) shows the lowest mean diversity, whereas the hybrids in between *V. latastei x aspis* (n=2), the highest.

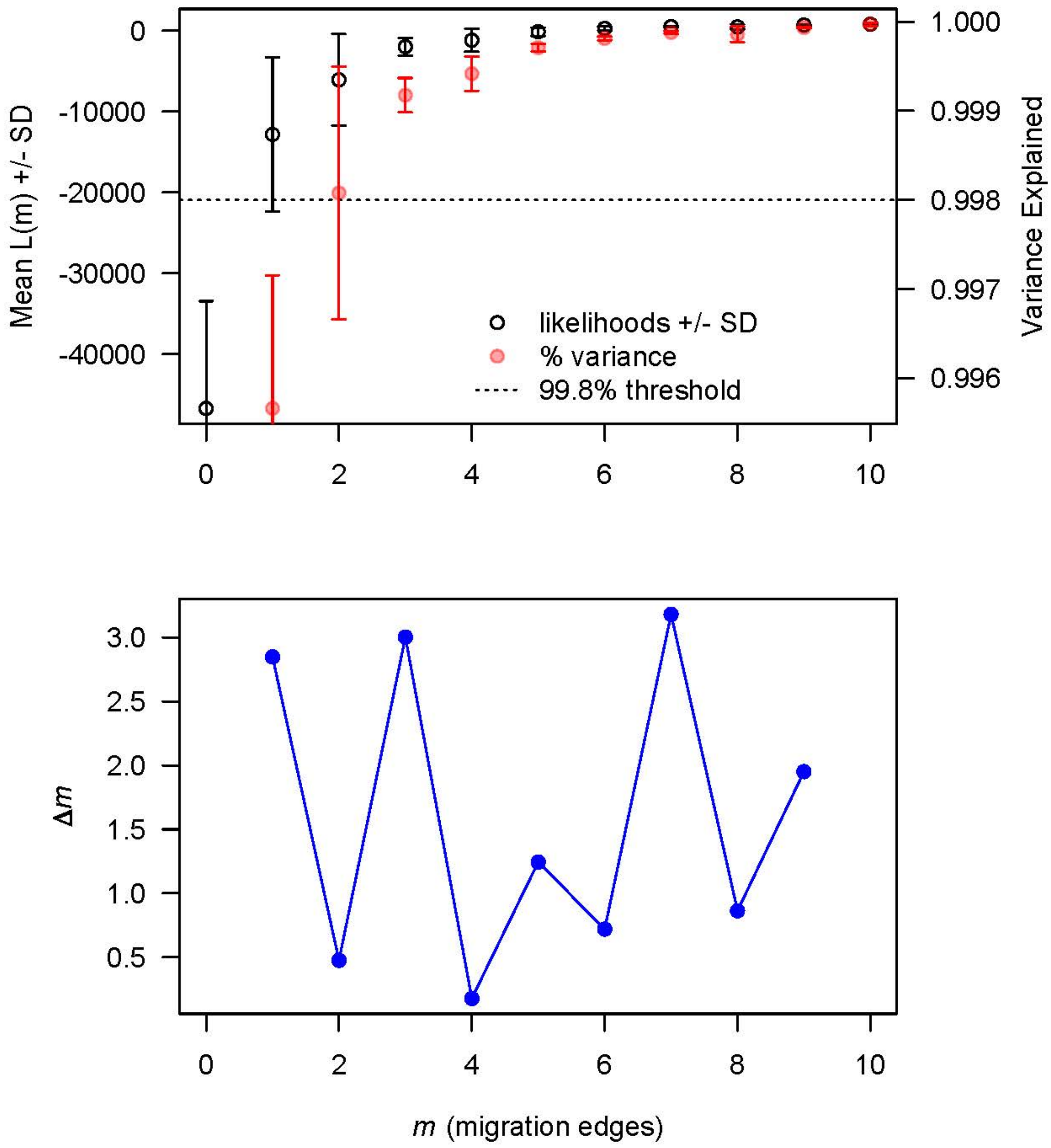


**Fig. S9. Output produced by OptM to find the optimum number of migrations in TreeMix.** At the top, the mean and standard deviation (SD) for the composite likelihood L(m) (left axis, black circles) and proportion of the variance explained (right axis, red circles). The 99.8% threshold is that recommended by Pickrell and Pritchard (2012). On the bottom, the second-order rate of change (*Δm*) across values of *m*. The peak is reached at *m*=7 edges.

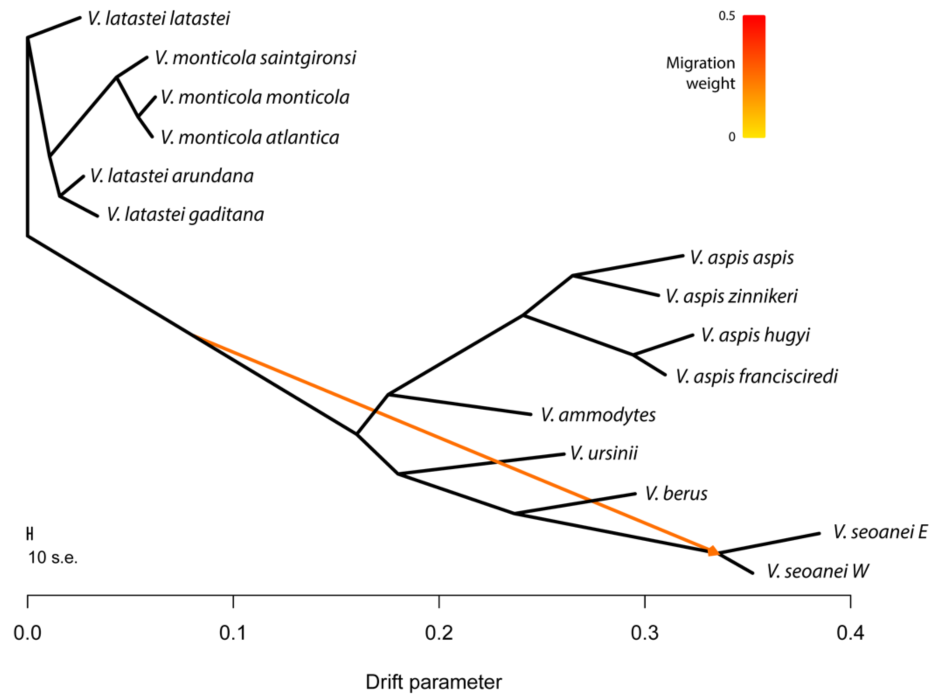


**Fig S10. Tree with one migration edge inferred by TreeMix.** *m*=1 is the second most likely value inferred by OptM. Color of the edge indicates the weight of migration. The drift parameter is a relative temporal measurement, and the scale bar indicates 10 times the average standard error of the relatedness among taxa based on the variance-covariance matrix of allele frequencies.

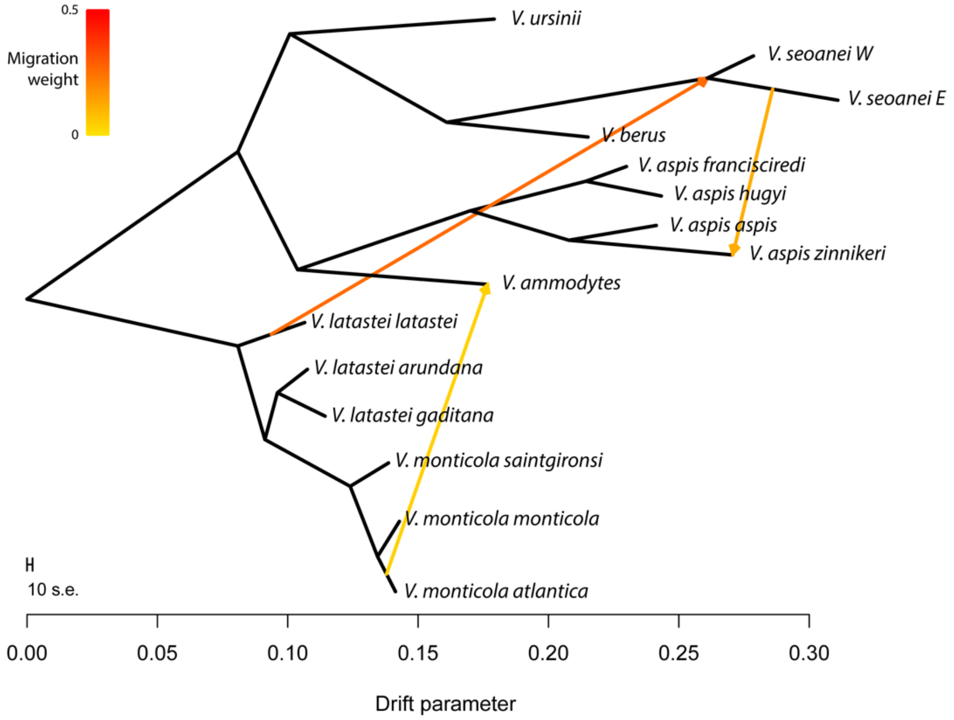


**Fig S11. Tree with three migration edges inferred by TreeMix.** *m*=3 is the third most likely value inferred by OptM. Color of the edges indicates the weight of migration. The drift parameter is a relative temporal measurement, and the scale bar indicates 10 times the average standard error of the relatedness among taxa based on the variance-covariance matrix of allele frequencies.

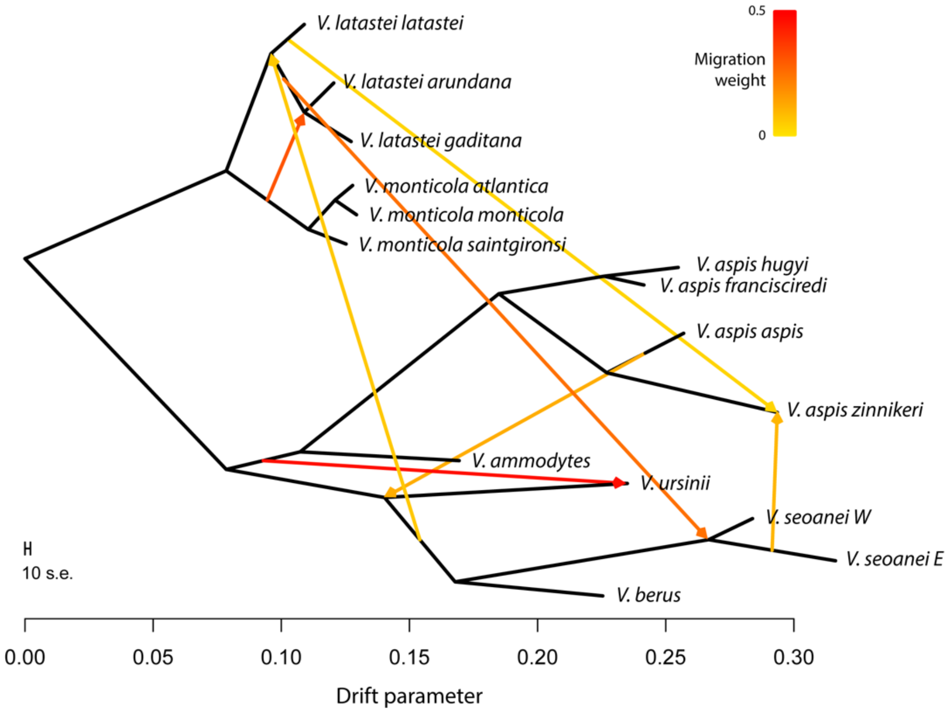


**Fig S12. Tree with seven migration edges inferred by TreeMix.** *m*=7 is the most likely value inferred by OptM. Color of the edges indicates the weight of migration. The drift parameter is a relative temporal measurement, and the scale bar indicates 10 times the average standard error of the relatedness among taxa based on the variance-covariance matrix of allele frequencies.

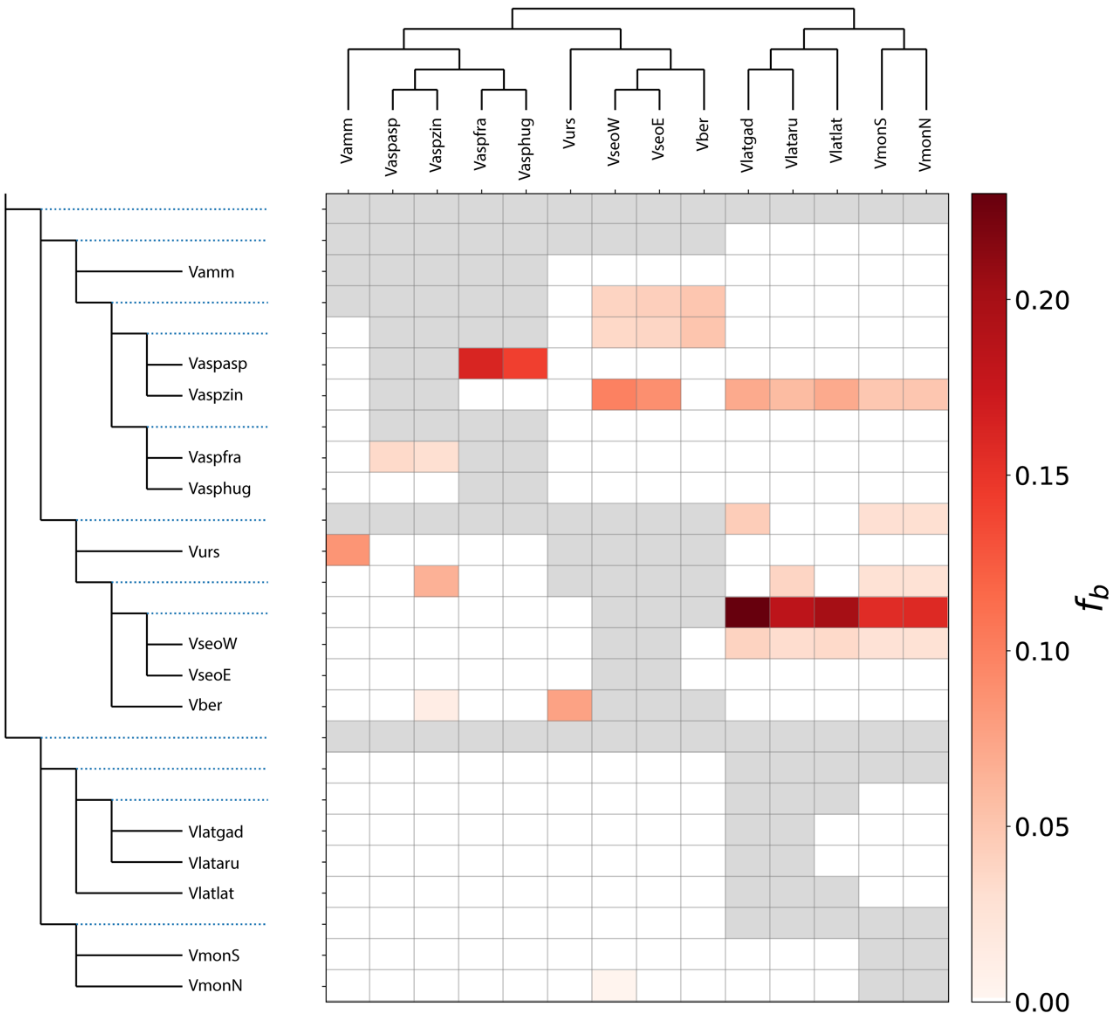


**Fig. S13.** Heatmap of *f-branch* ($f_b$) statistic, representing excess allele sharing between branches of y- and x-axes. Statistics have been corrected by a robust clustering threshold to avoid false positives due to homoplasy instead of introgression (see sensitive clustering threshold results in Fig. 4A). Blank cells depict values not statistically supported, whereas grey cells indicate comparisons that cannot be made due to tree topology. Exact *f-branch* values are shown in Table S4. Vamm stands *V. ammodytes*, Vaspasp for *V. aspis aspis*, Vaspzin for *V. aspis zinnikeri*, Vaspfra for *V. aspis francisciredi*, Vasphug for *V. aspis hugyi*, Vurs for *V. ursinii*, VseoW for *V. seoanei* West, VseoE for *V. seoanei* East, Vber for *V. berus*, Vlatgad for *V. latastei gaditana*, Vlataru for *V. latastei arundana*, Vlatlat for *V. latastei latastei*, Vmon S for *V. monticola* South and VmonN for *V. monticola* North.

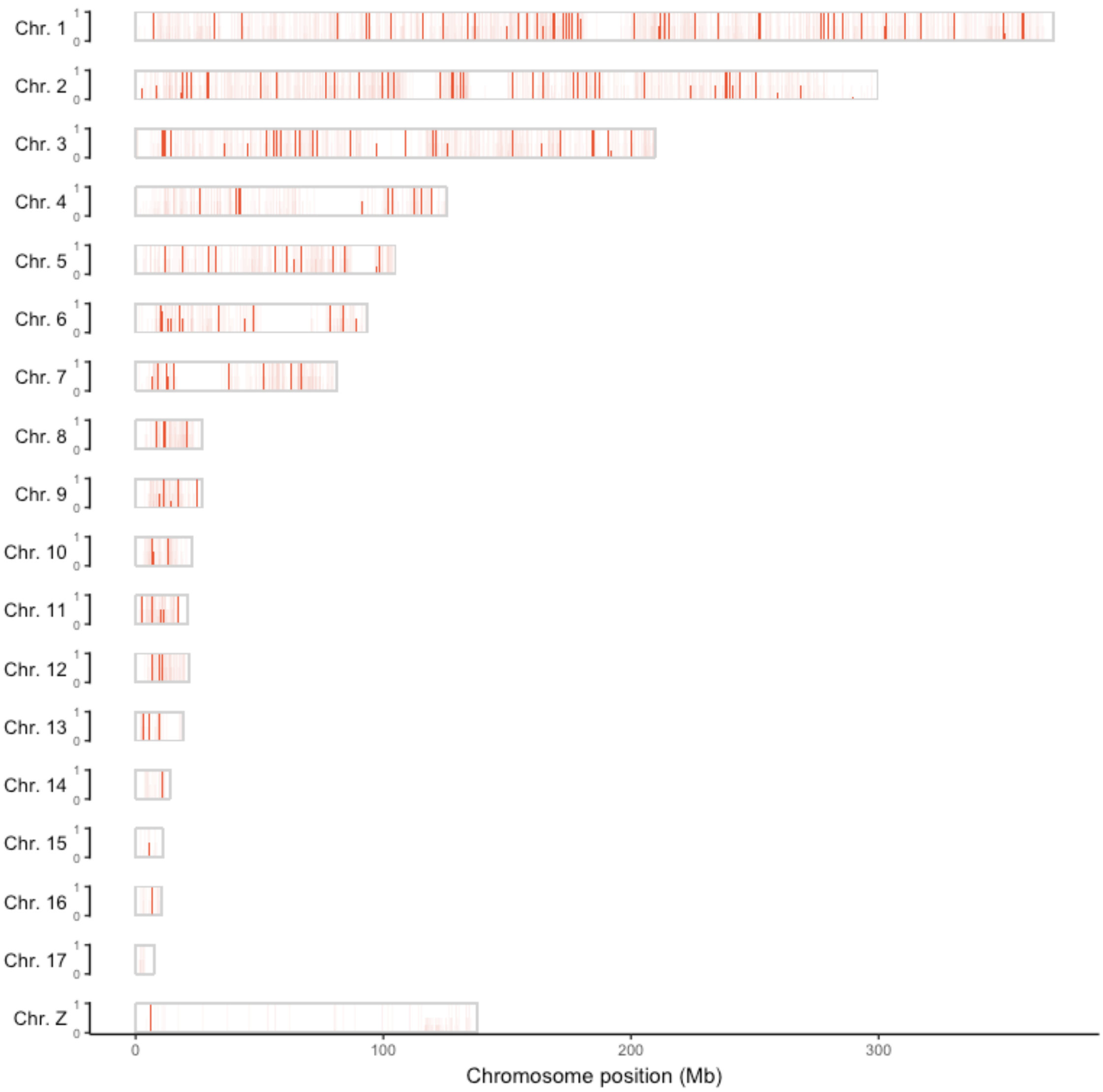


**Fig. S14. Landscape of introgression from *V. latastei* to *V. seoanei*, shown in *V. latastei*'s reference genome highlighting windows with excess allele-sharing.** Introgression between these species seems to be pervasive, although some areas display higher intensity whereas others, including the whole Z chromosome, are less introgressed (i.e., "introgression deserts").

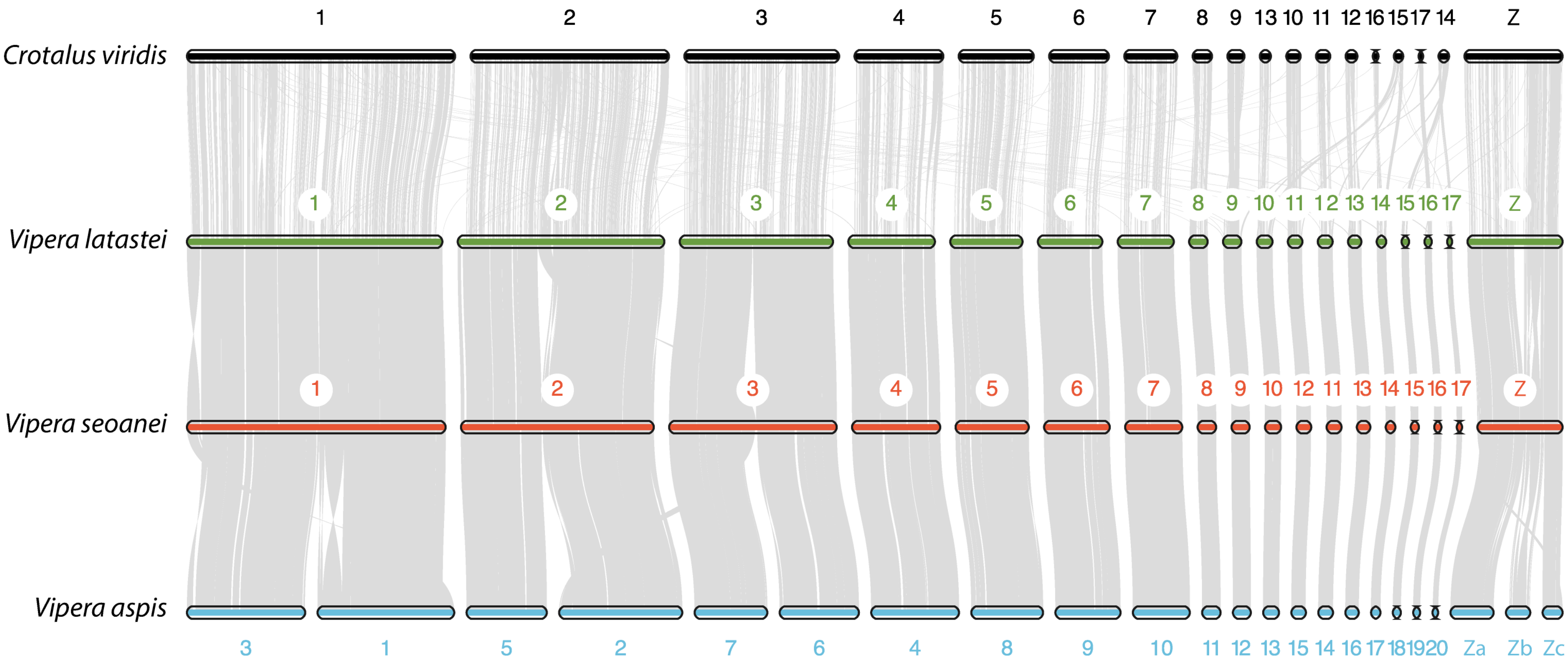


**Fig. S15. Macrosynteny across the three old species groups in *Vipera*, represented by the three Iberian species, and a rattlesnake as outgroup**. Some microchromosomal rearrangements have occurred in between the sister subfamilies Crotalinae and Viperinae, while macrochromosomes have been mostly conserved. The ancestral chromosome number, i.e., 2n=36 is conserved in *V. latastei* and *V. seoanei* but increased in *V. aspis* as a result of three fission events in the three major macrochromosomes. Z chromosome in *V. aspis* appears fragmented due to lower Omni-C coverage during the assembly, since it was sequenced from a female (ZW) and the other three assembled genomes correspond to males (ZZ). A fragment of *V. latastei* chromosome 2 is translocated to *V. seoanei* chromosome 3. Some chromosomes have been reoriented on the grounds of a better visualization.

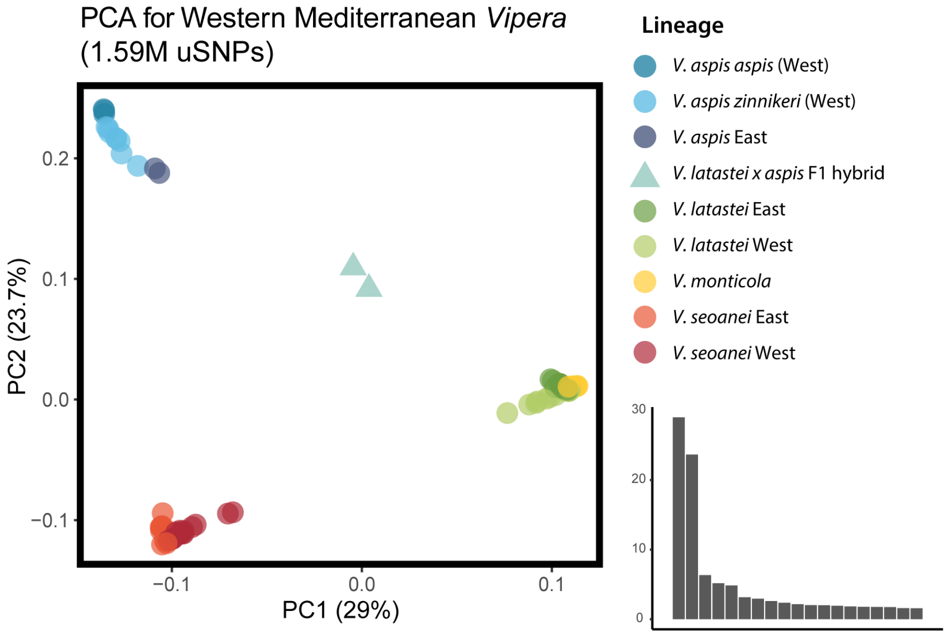


**Fig. S16. Genomic PCA of Western Mediterranean *Vipera* spp. performed with 1.59M uSNPs and 90 whole genomes.** PC1 (29% of explained variance) splits *Vipera latastei-monticola* complex from *V. seoanei* and *V. aspis*, whereas PC2 (23.7%) splits the latter two. Two individuals seem to be F1 hybrids of *V. latastei x aspis*, found in sympatry areas of N and NE Spain. The Western lineages of *V. seoanei* and *V. latastei* seem to be closer, probably due to higher admixture between them. More discretely, the Eastern lineages of *V. seoanei* and *V. latastei* seem to be relatively closer, but more slightly, to *V. aspis*. The bar plot on the right shows the explained variance by component.

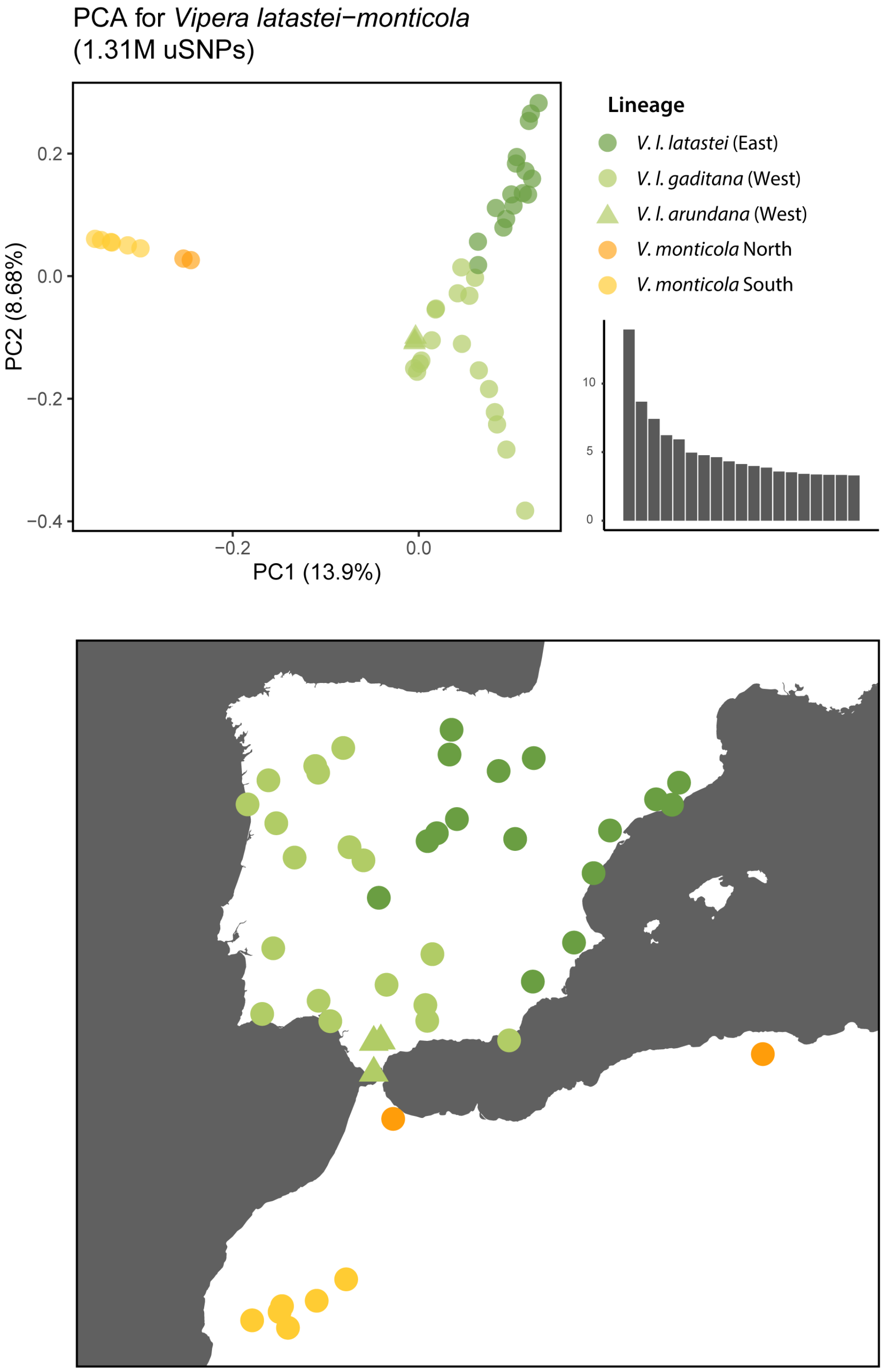


**Fig. S17. Genomic PCA of the *latastei* group (top), performed with 1.31M uSNPs and 45 whole genomes and geographic origin of the samples (bottom).** PC1 (13.9% of explained variance) splits *Vipera latastei* from *V. monticola*, as well as depicts a South-North gradient within each. PC2 (8.68%) explains an East-West gradient of *V. latastei* across the Iberian Peninsula. The individuals belonging to *V. latastei arundana* (triangles) fall within other Western *V. latastei*, which mostly corresponds to *V. latastei gaditana*. *Vipera latastei* East mostly corresponds to the nominotypical *V. latastei latastei*. The bar plot on the right shows the explained variance by component.

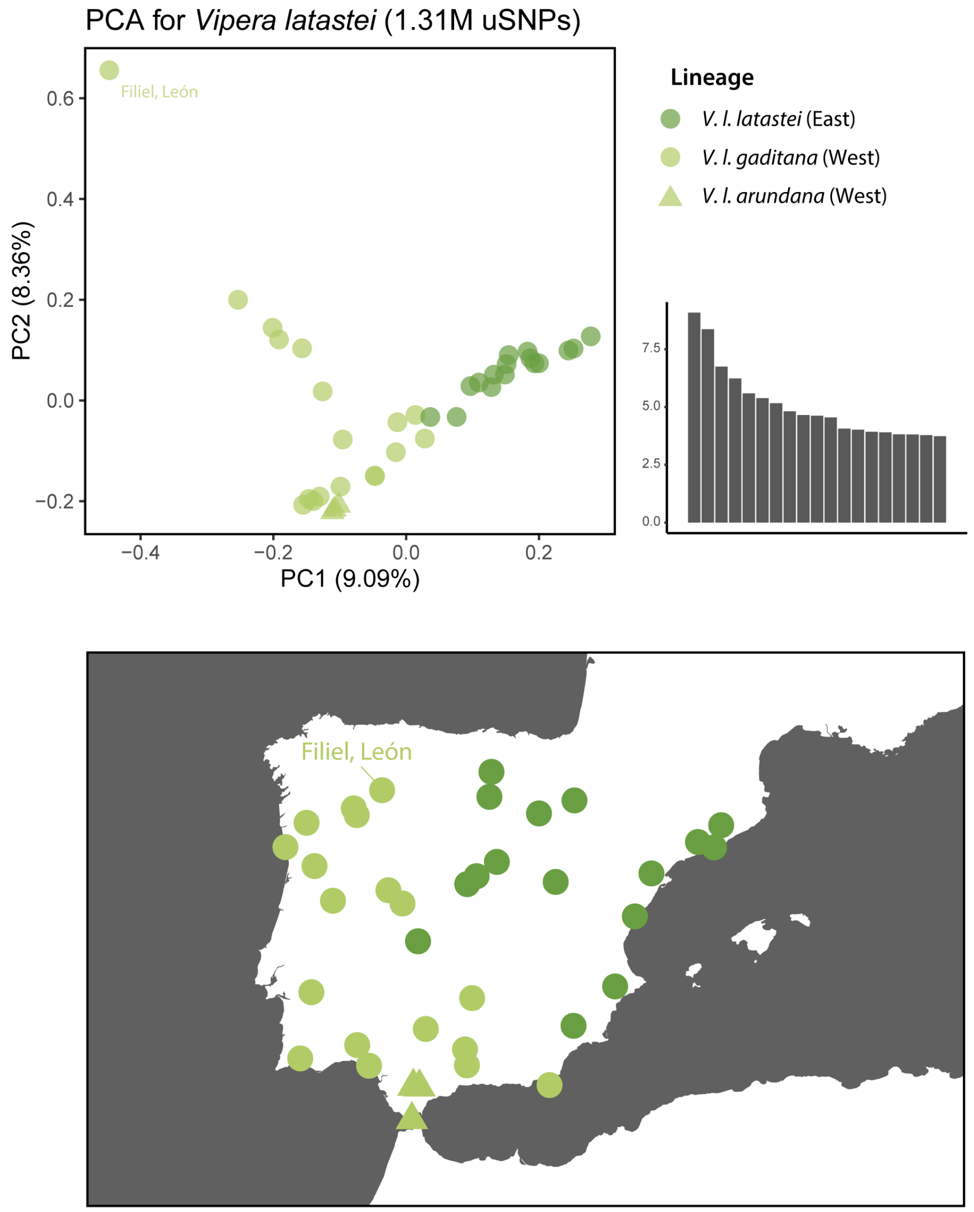


**Fig. S18. Genomic PCA of *Vipera latastei* (top), performed with 1.31M uSNPs and 37 whole genomes and geographic origin of the samples (bottom).** PC1 (9.09% of explained variance) depicts a West-East gradient, whereas PC2 (8.36%) explains North-South variation, including admixture with *V. seoanei* in the northwest, more pronounced in an individual from Filiel, León. The individuals belonging to *V. latastei arundana* (triangles) fall within other Western *V. latastei*, which mostly corresponds to *V. latastei gaditana*. *Vipera latastei* East mostly corresponds to *V. latastei latastei*. The bar plot on the right shows the explained variance by component.

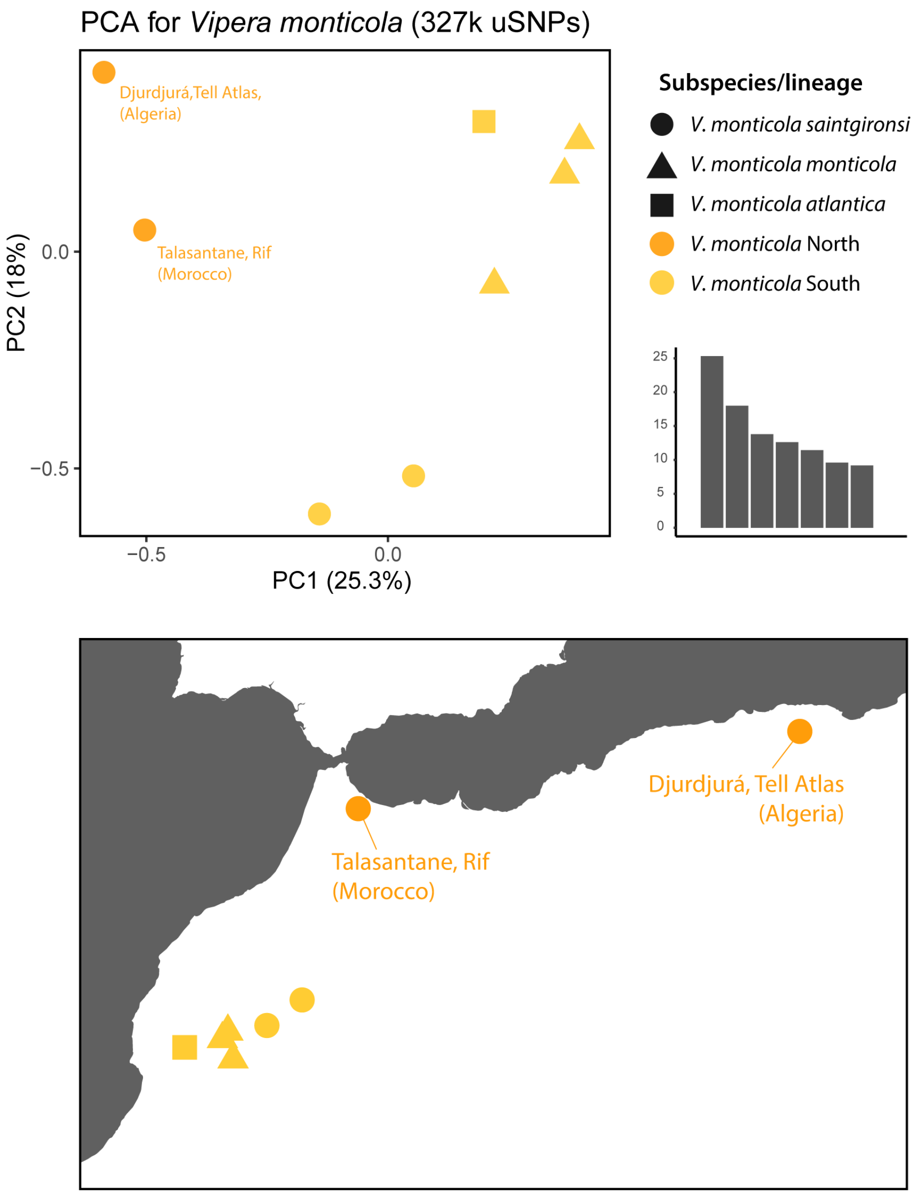


**Fig. S19. Genomic PCA of *Vipera monticola* (top), performed with 327k uSNPs and eight whole genomes and geographic origin of the samples (bottom).** PC1 (25.3% of explained variance) depicts a North-South gradient, from the northern populations attributed to *V. monticola saintgironsi* to the Atlas populations in the south, attributed to three different subspecies. Southern (Atlas) populations form a subtle cluster along PC1 Thus, we differentiate *V. monticola* into North and South (Atlas) lineages instead of using mitochondrial subspecies. PC2 (18%) explores mostly differences from the Western to the Eastern Atlas within the South lineage. The bar plot on the right shows the explained variance by component.

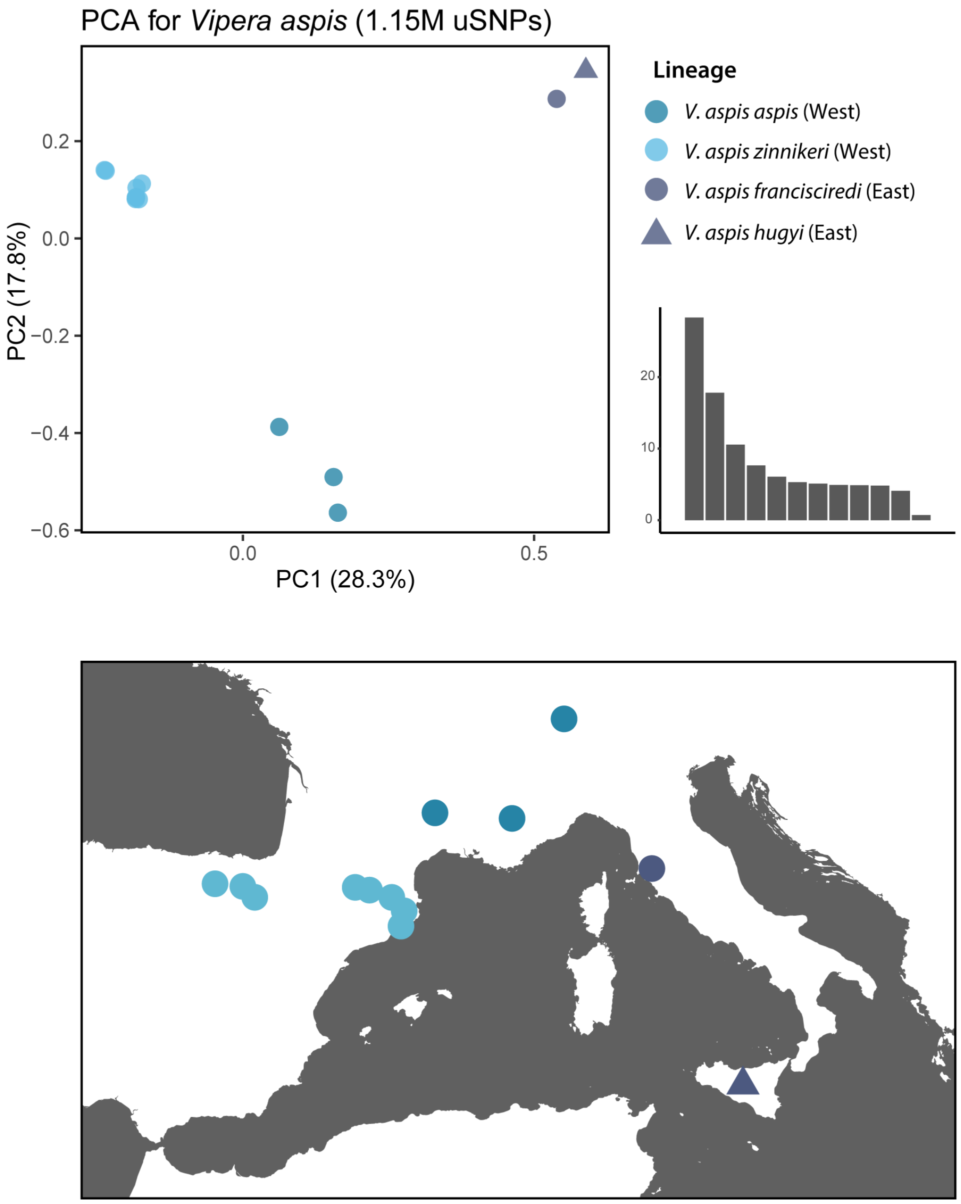


**Fig. S20. Genomic PCA of *Vipera aspis* (top), performed with 1.15M uSNPs and 13 whole genomes and geographic origin of the samples (bottom).** PC1 (28.3% of explained variance) splits *Vipera aspis* into three clusters: *V. a. zinnikeri* and *V. a. aspis*, closer between them, and an Eastern cluster composed by individuals from two subspecies: *V. a. francisciredi* and *V. a. hugyi*. PC2 (17.8%) separates *V. a. aspis* from the rest. The bar plot on the right shows the explained variance by component.

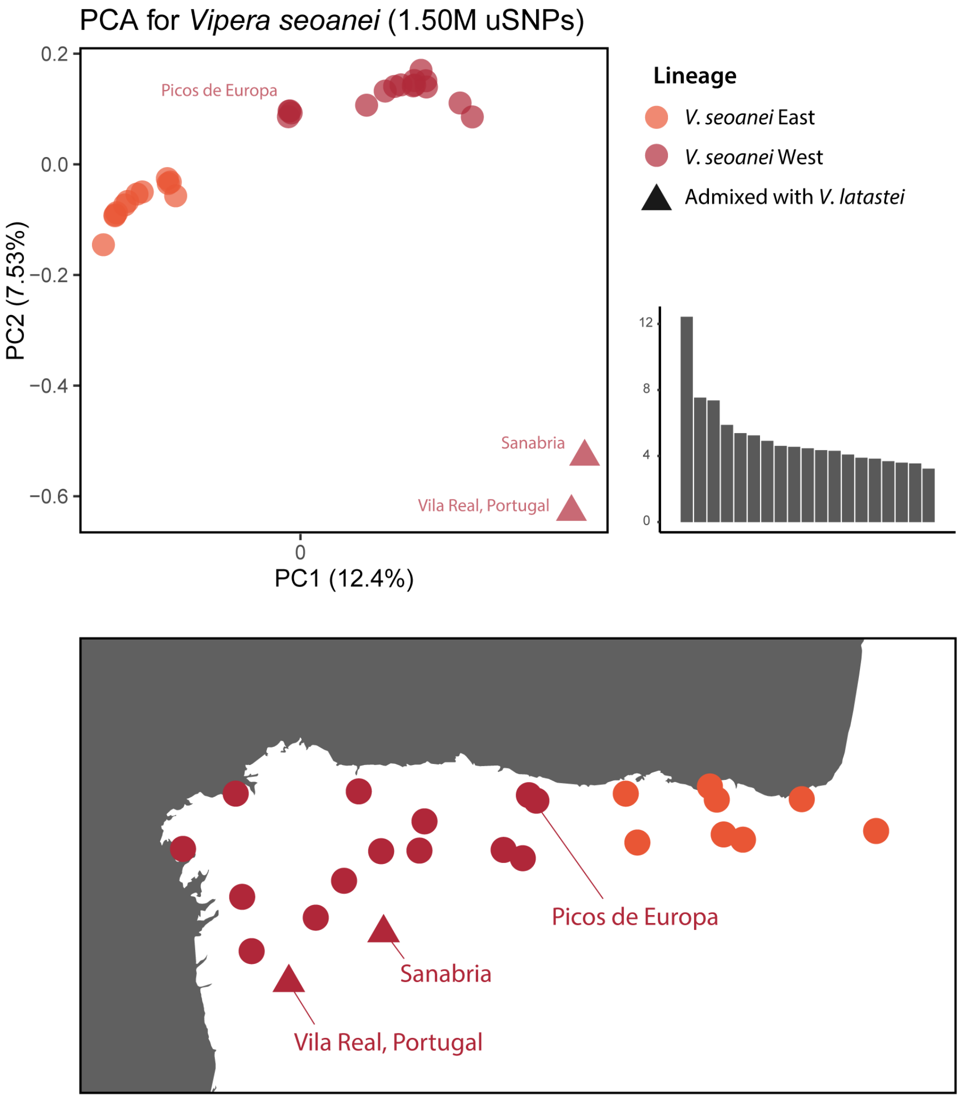


**Fig. S21. Genomic PCA of *Vipera seoanei* (top), performed with 1.50M uSNPs and 30 whole genomes and geographic origin of the samples (bottom).** PC1 (12.4% of explained variance) splits *Vipera seoanei* into Eastern and Western clades, with Picos de Europa populations exhibiting a more intermediate position. PC2 (7.53%) mainly separates two admixed Western individuals with *V. latastei* (triangles). The bar plot on the right shows the explained variance by component.

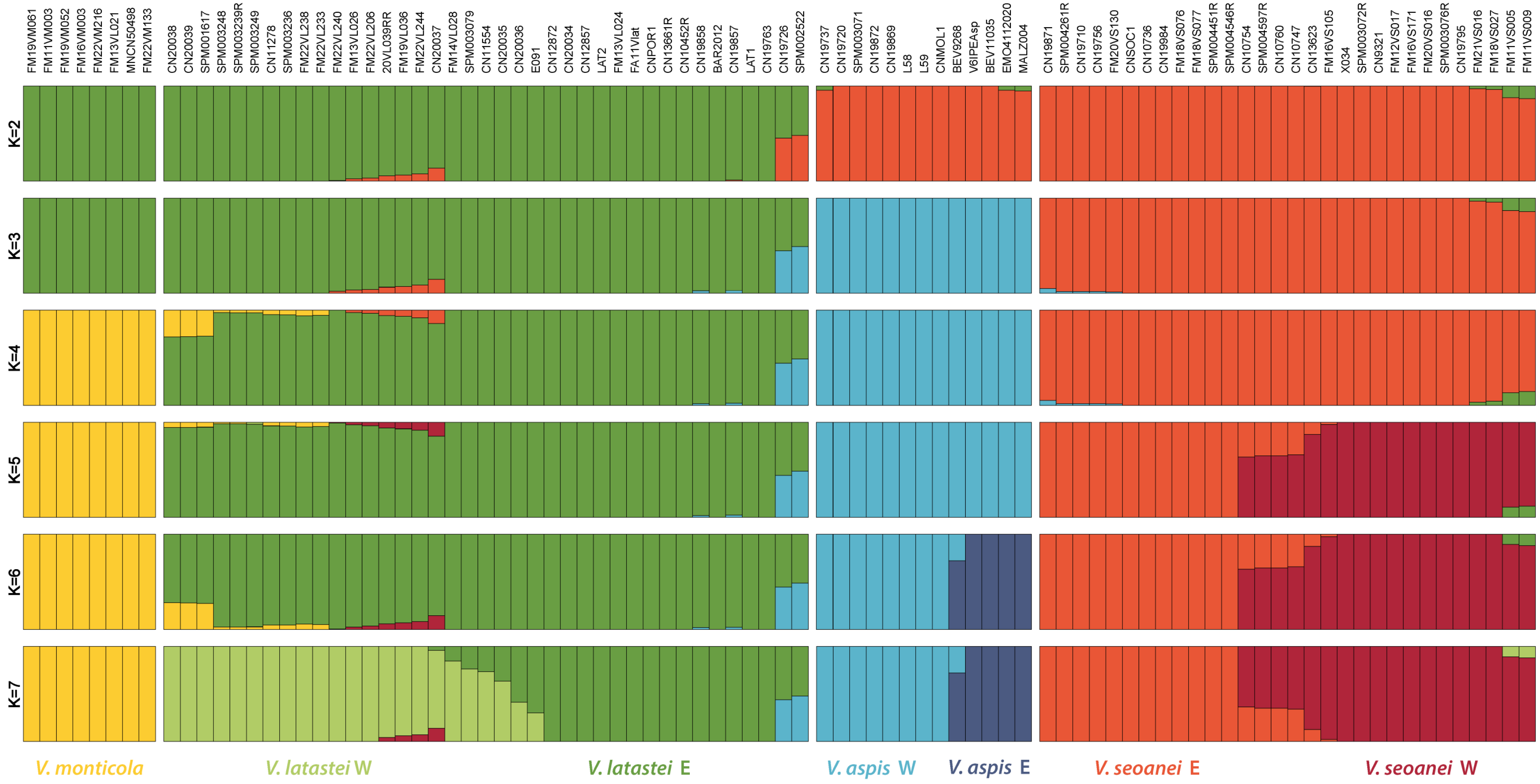


**Fig. S22. Admixture results for Western Mediterranean *Vipera* spp., using 1.59M uSNPs and 90 whole genomes, from K=2 to K=7.** Most likely K is 4, coinciding with the number of species, although K=7, which is depicted as well in Fig. 5A1-2, is more meaningful as it explores intraspecific structure within *V. latastei*, *V. seoanei* and *V. aspis*. Sample localities are shown in Table S1.

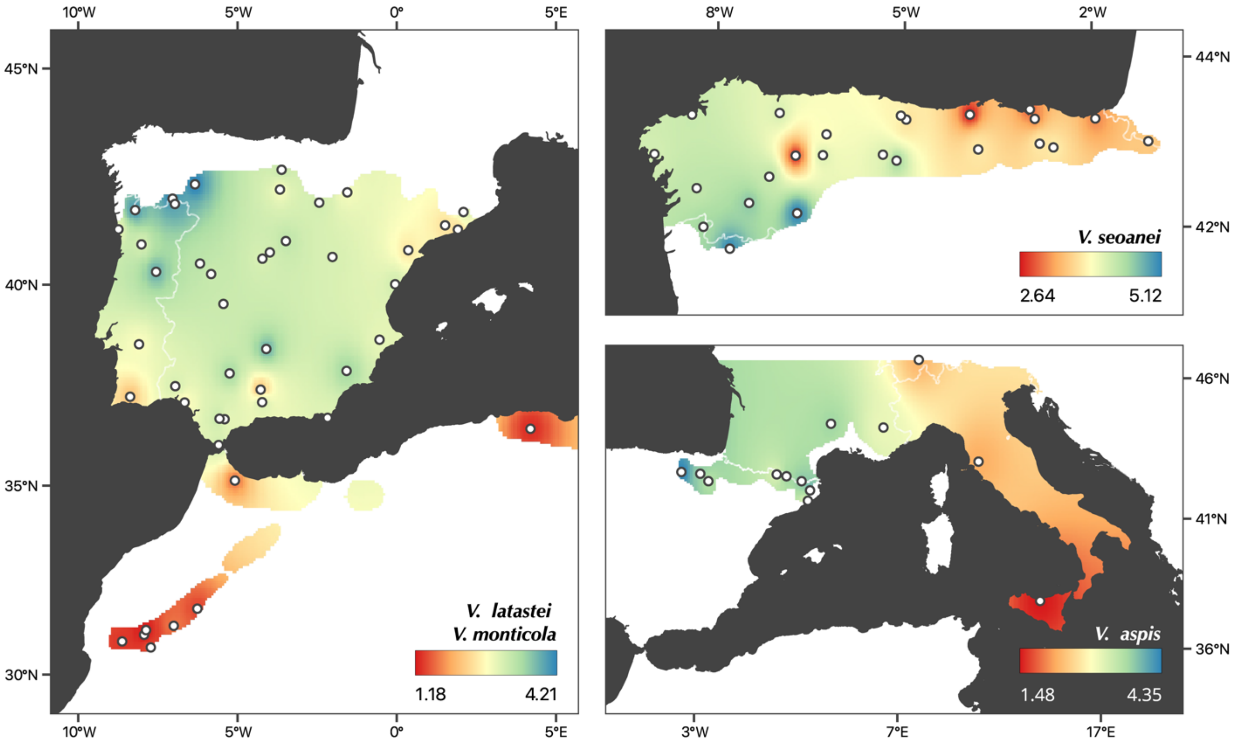


**Fig. S23. Genome-wide heterozygosity interpolations per individual for *V. latastei-monticola* on the left, *V. seoanei* on the upper right corner and *V. aspis* on the bottom right.** Diversity increases from south to north in *V. latastei-monticola*, and from East to West in *V. aspis* and *V. seoanei*. Highest heterozygosity values for the three Iberia species are found in sympatry areas.

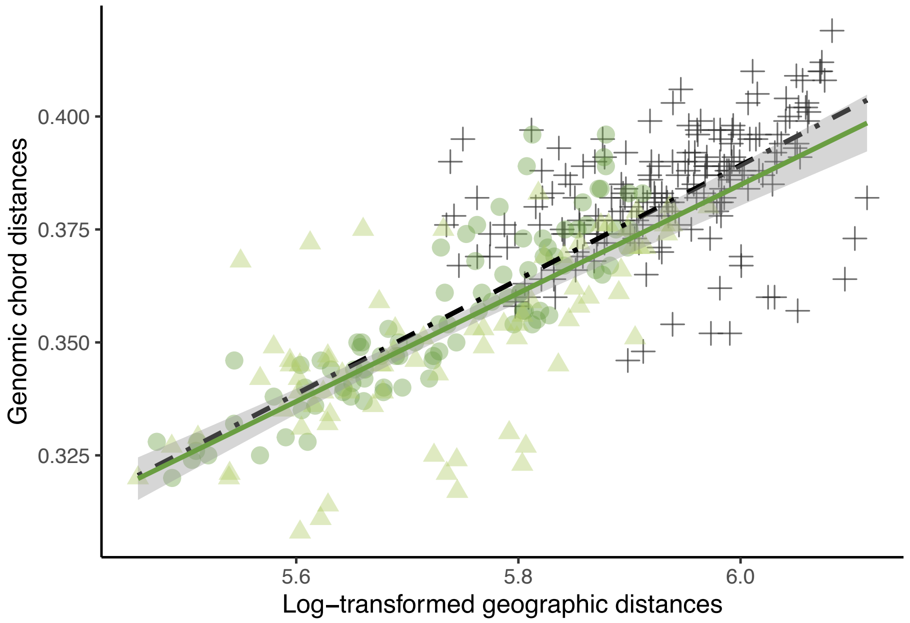


**Fig. S24. Relationship between genomic and log-transformed geographical distances in pairs of individuals of the Western and Eastern lineages of *V. latastei* to explore isolation-by-distance (IBD) patterns.** Green circles and light green triangles indicate distances between individuals belonging to Eastern and Western lineages, respectively. Black crosses indicate distances between individuals belonging to different intraspecific lineages. The solid green line is the regression line fitted only for within-lineage comparisons (i.e., circles and triangles) and its 95%CI in grey. Dashed black line is the regression fitted for all comparisons, either within- or between-lineage (circles, triangles, and crosses), i.e., the IBD pattern needed to explain lineage differences exclusively due to distance. Both regressions do not significantly differ, therefore, genomic differences between *V. latastei* lineages can be explain by IBD.

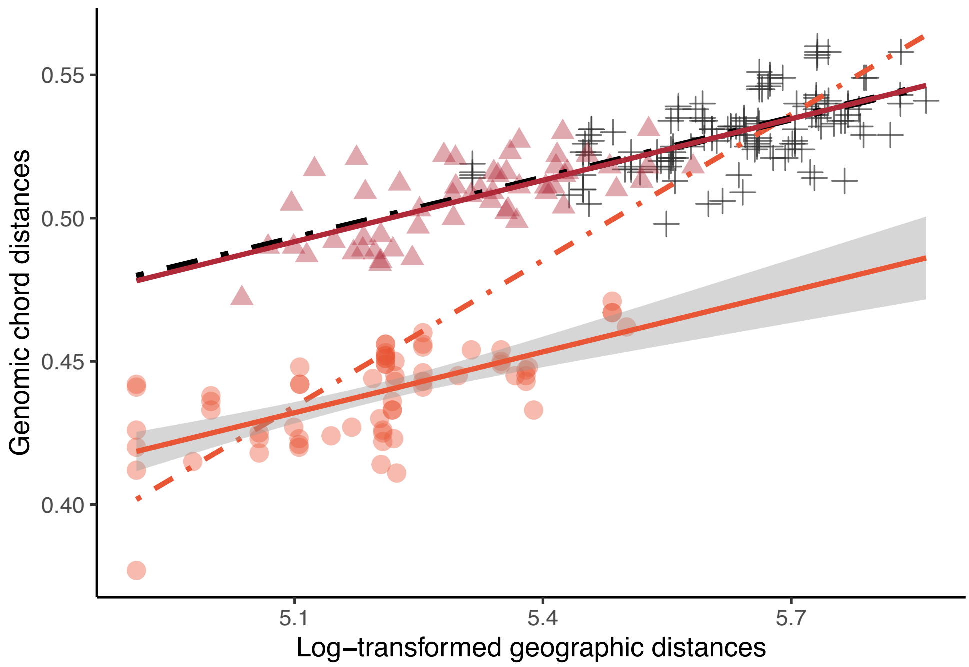


**Fig. S25. Relationship between genomic and log-transformed geographical distances in pairs of individuals of the Western and Eastern lineages of *V. seoanei* to explore isolation-by-distance (IBD) patterns.** Red circles and maroon triangles indicate distances between individuals belonging to Eastern and Western lineages, respectively. Black crosses indicate distances between individuals belonging to different intraspecific lineages. The solid red line is the regression line fitted only for within-East comparisons (i.e., circles) and its 95%CI in grey, whereas the solid maroon line is fitted for within-West comparisons (i.e., triangles). Dashed red and black lines are the regressions fitted for either within-East or within-West and between-lineage comparisons (i.e., either circles or triangles, plus crosses). The intercept of the red and maroon solid regressions is different, despite their similar slope. The slope of the black and red dashed regressions significantly differs. IBD in the Western lineage can explain differentiation of the whole range, but IBD in the Eastern cannot.

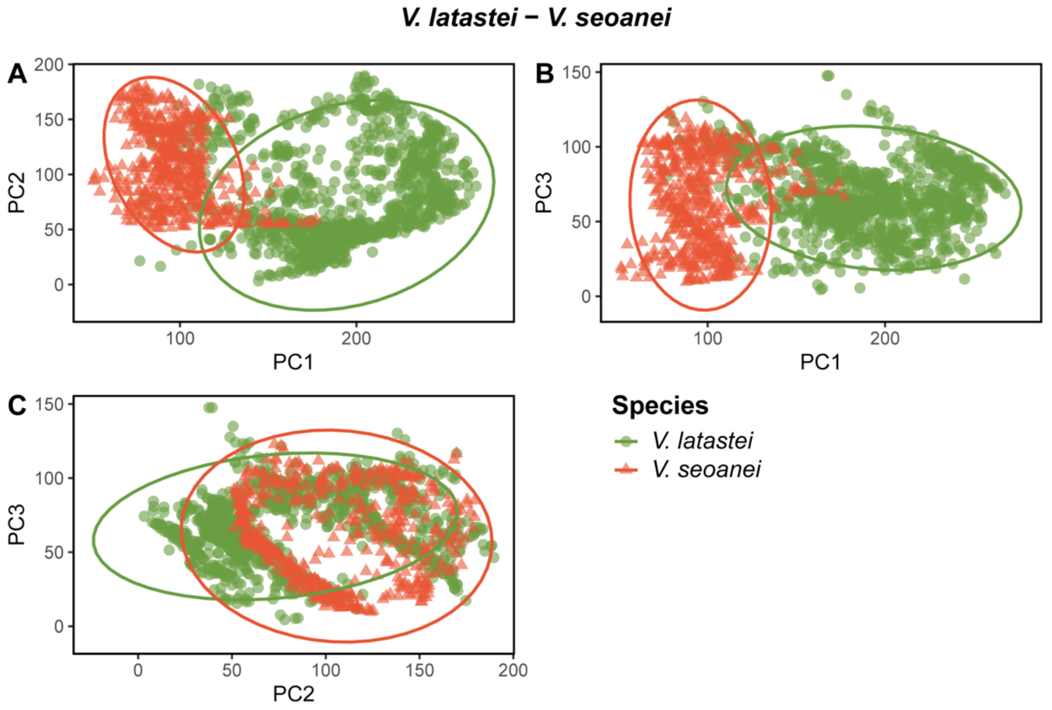


**Fig. S26. Environmental PCA for all *V. latastei* (green circles, n=1,105) and all *V. seoanei* (red triangles, n=508).** Ecological differences between these species are best explained by PC1, i.e., an Eurosiberian *vs.* Mediterranean gradient. Dispersion of the occurrences of each species with 95% CI are depicted.

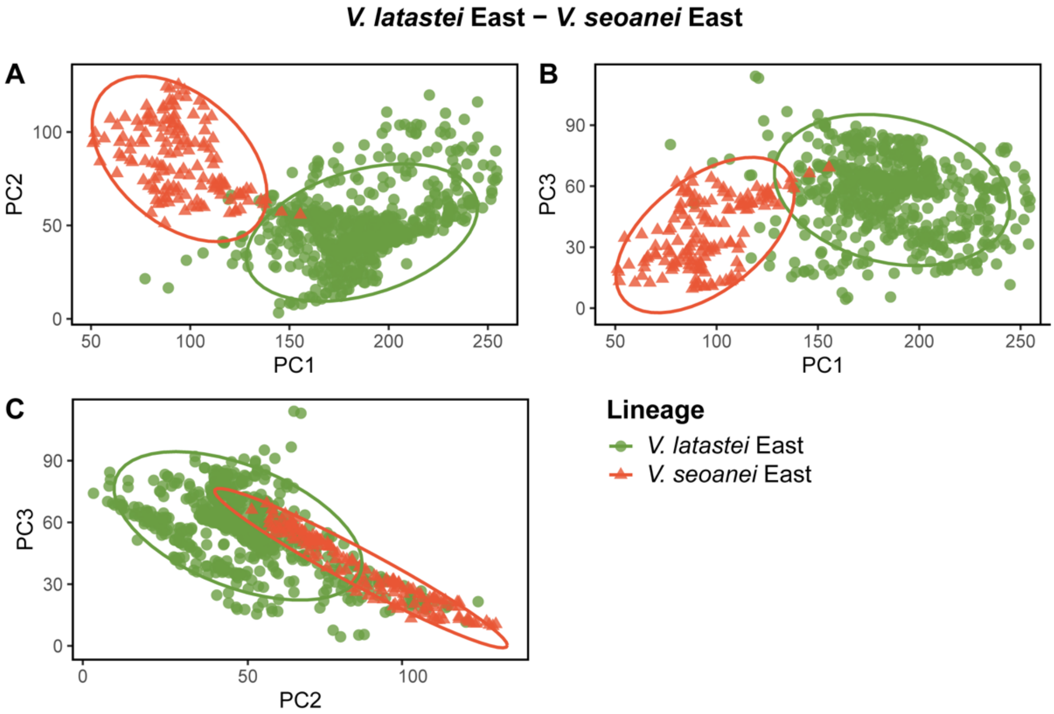


**Fig. S27. Environmental PCA for Eastern *V. latastei* (green circles, n=617) and Eastern *V. seoanei* (red triangles, n=146).** Ecological differences between these lineages are best explained by PC1, i.e., an Eurosiberian *vs.* Mediterranean gradient. Dispersion of the occurrences of each species with 95% CI are depicted.

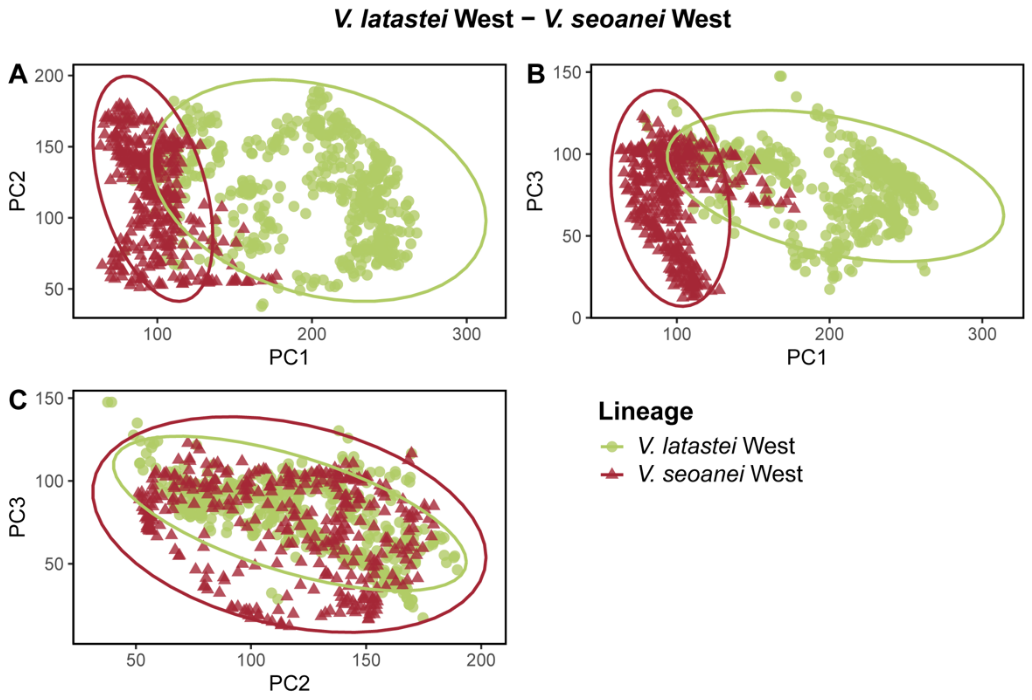


**Fig. S28. Environmental PCA for Western *V. latastei* (light green circles, n=488) and Western *V. seoanei* (maroon triangles, n=362).** Ecological differences between these lineages are best explained by PC1 *vs*. PC2. Dispersion of the occurrences of each species with 95% CI are depicted.

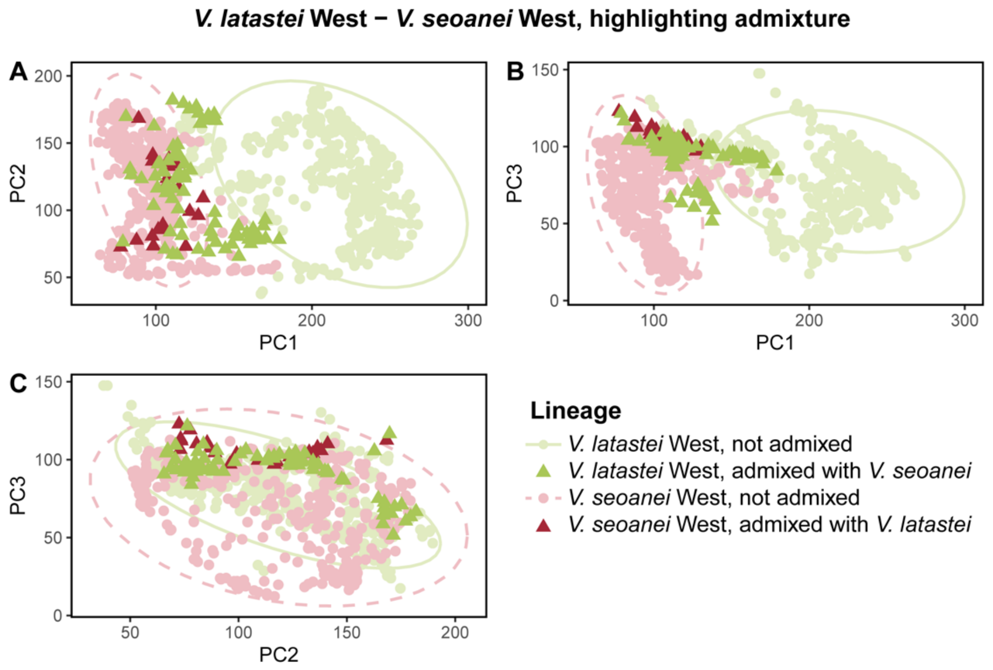


**Fig. S29. Environmental PCA for Western *V. latastei* (light green, n=488) and Western *V. seoanei* (maroon, n=362), highlighting admixed individuals.** Ecological differences between these lineages are best explained by PC1 *vs*. PC2. Dispersion of the occurrences of each species for unadmixed populations with 95% CI are depicted (dashed for Western *V. seoanei*). Circles represent occurrences of putatively unadmixed individuals, whereas triangles indicate admixed individuals. *Vipera latastei* individuals from admixed populations seems to inhabit more Eurosiberian areas than expected for its lineage.

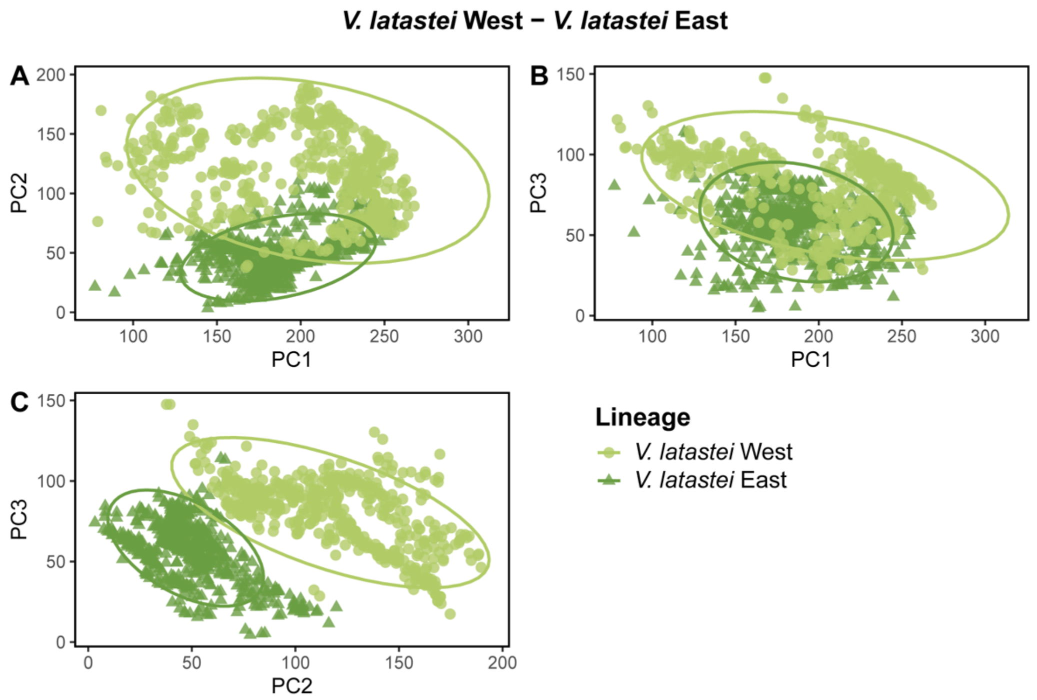


**Fig. S30. Environmental PCA for the Western (light green circles, n=488) and Eastern (green triangles, n=617) lineages of *V. latastei*.** Ecological differences between these lineages are best explained by PC2 vs. PC3. Dispersion of the occurrences of each species for unadmixed populations with 95% CI are depicted.

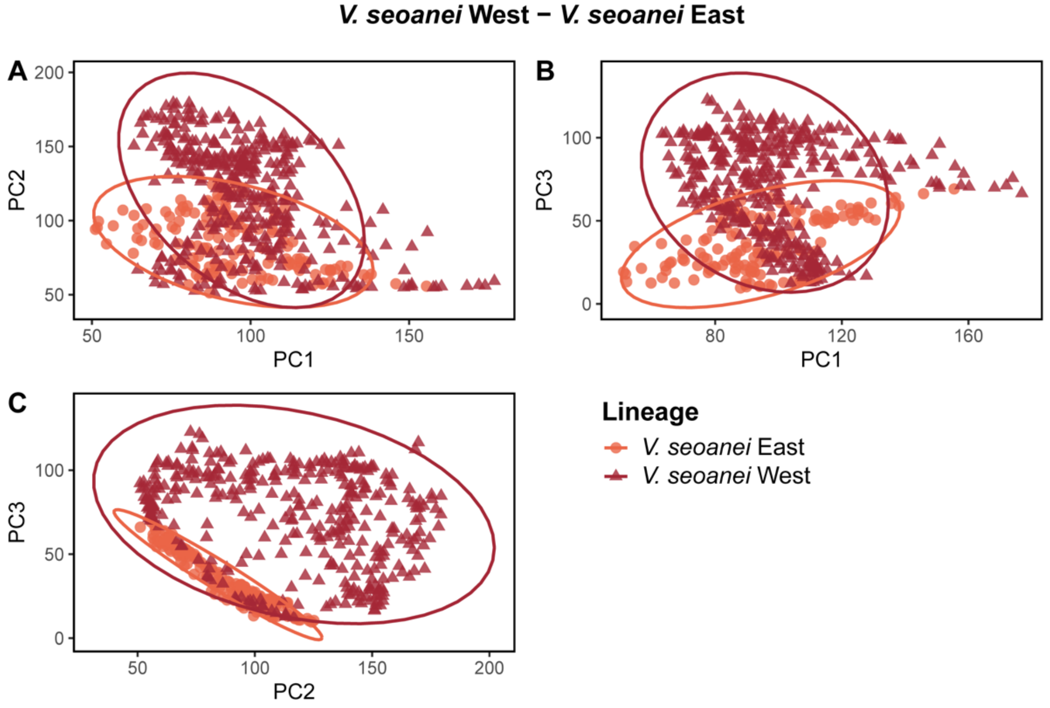


**Fig. S31. Environmental PCA for the Western (maroon triangles, n=362) and Eastern (red circles, n=146) lineages of *V. seoanei*.** *Vipera seoanei* lineages show great overlap in the three PCs. Dispersion of the occurrences of each species with 95% CI are depicted.

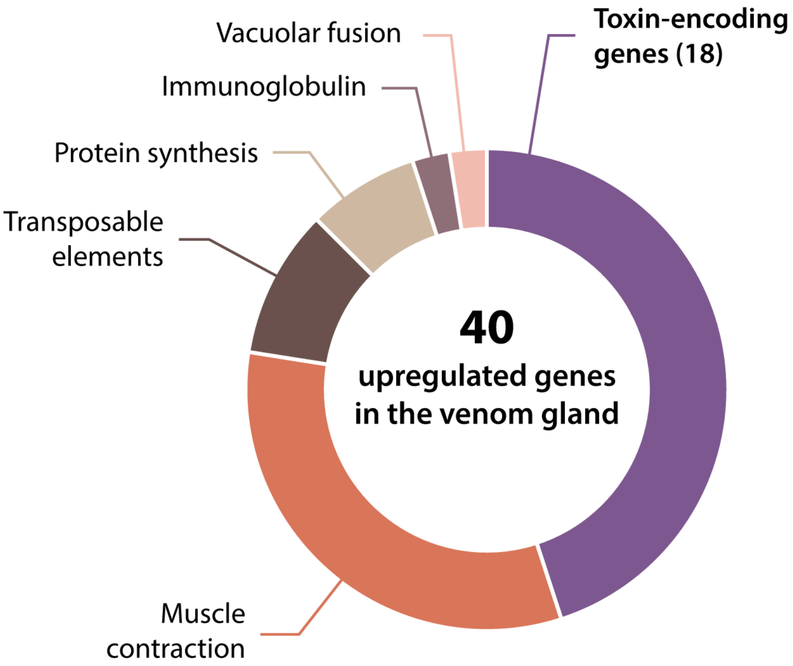


**Fig. S32. Pie chart of the differentially upregulated genes in the venom gland of the Iberian vipers.** From 40 detected genes, 18 are toxin-encoding genes. The rest are related to muscle contraction (13 genes), transposable elements (4). Protein synthesis (3), vacuolar fusion (1) or immunoglobulins (1).

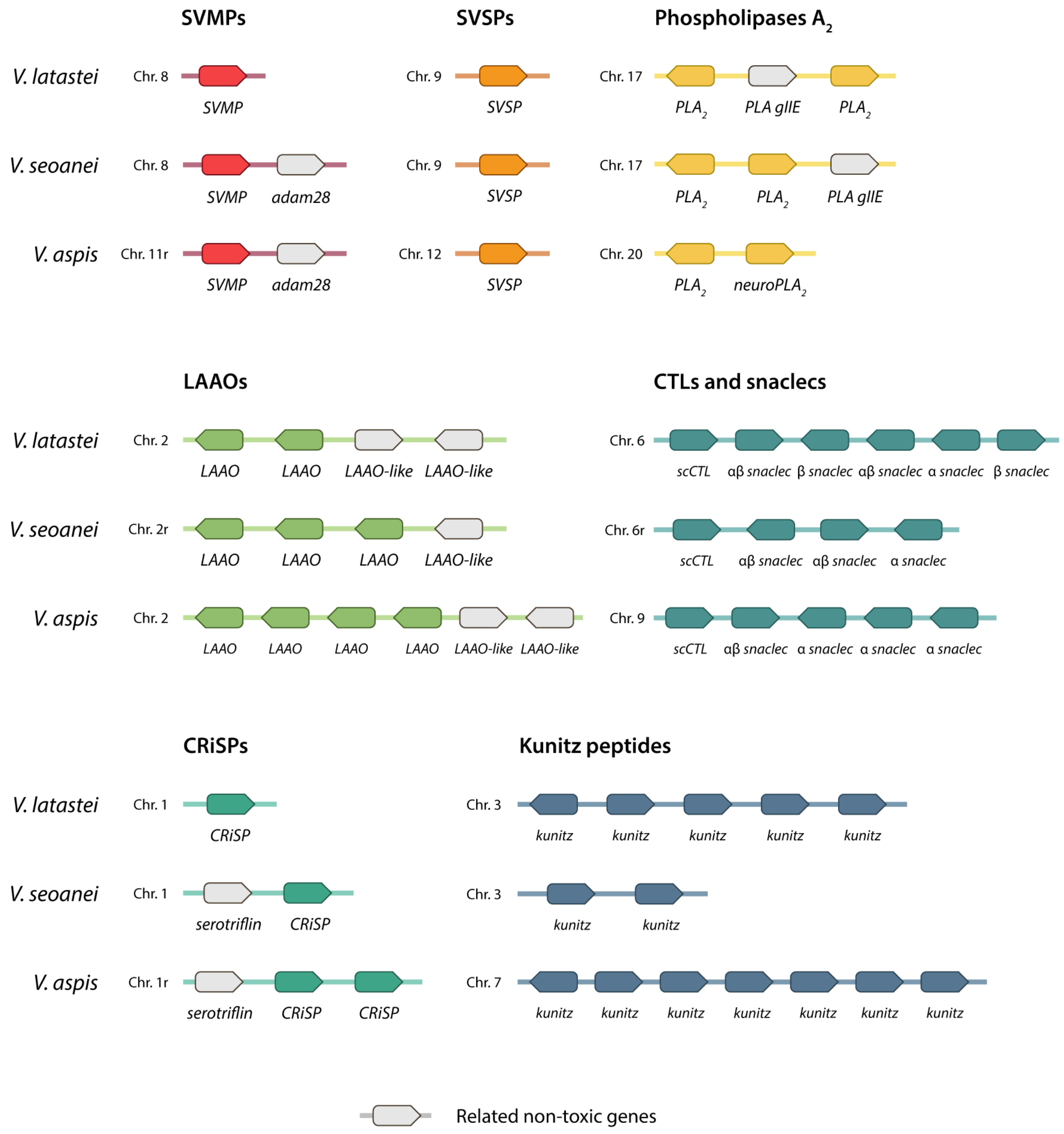


**Fig. S33. Microsynteny of the main toxin-encoding gene families for the three Iberian vipers.** Genes in grey are non-toxic, but potentially related to the toxin copies. Toxin-encoding genes from other non-main families are not represented.

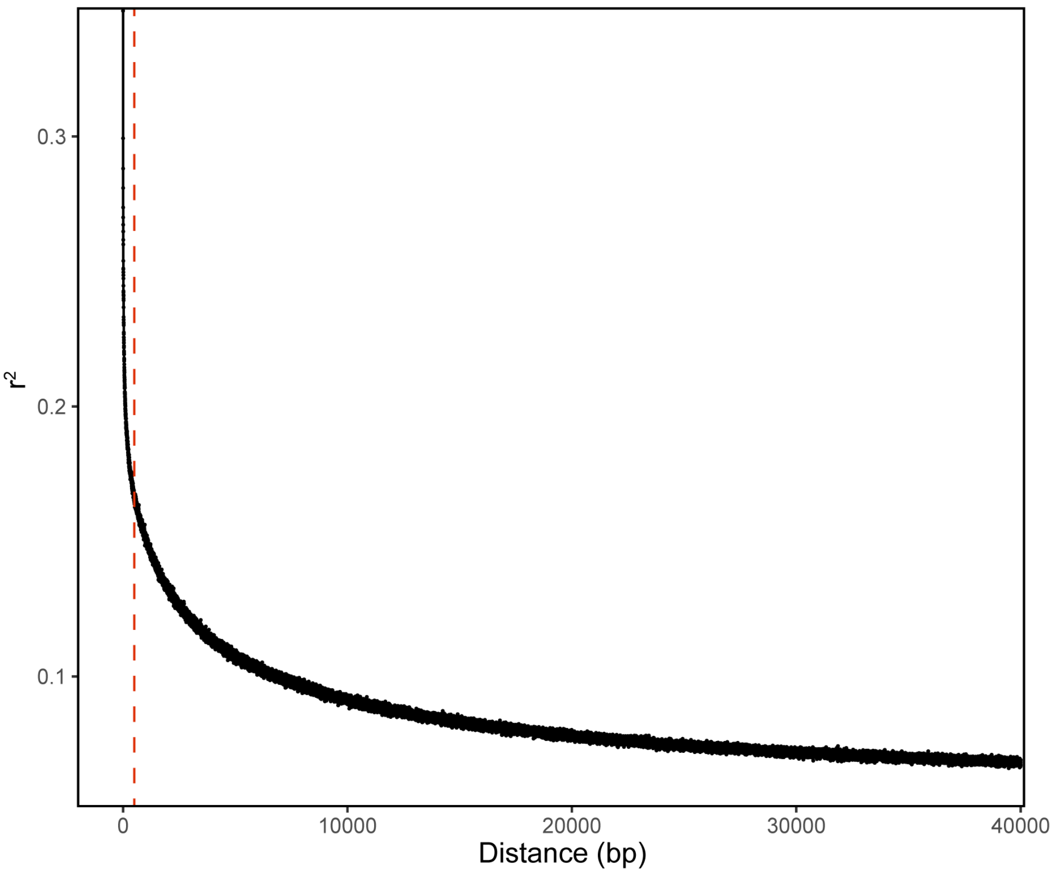


**Fig. S34. Linkage disequilibrium (LD) decay in *Vipera seoanei* across sites.** The vertical line marks LD at sites 0.5 kbp apart. Correlation between sites at this distance is lower than 20%.

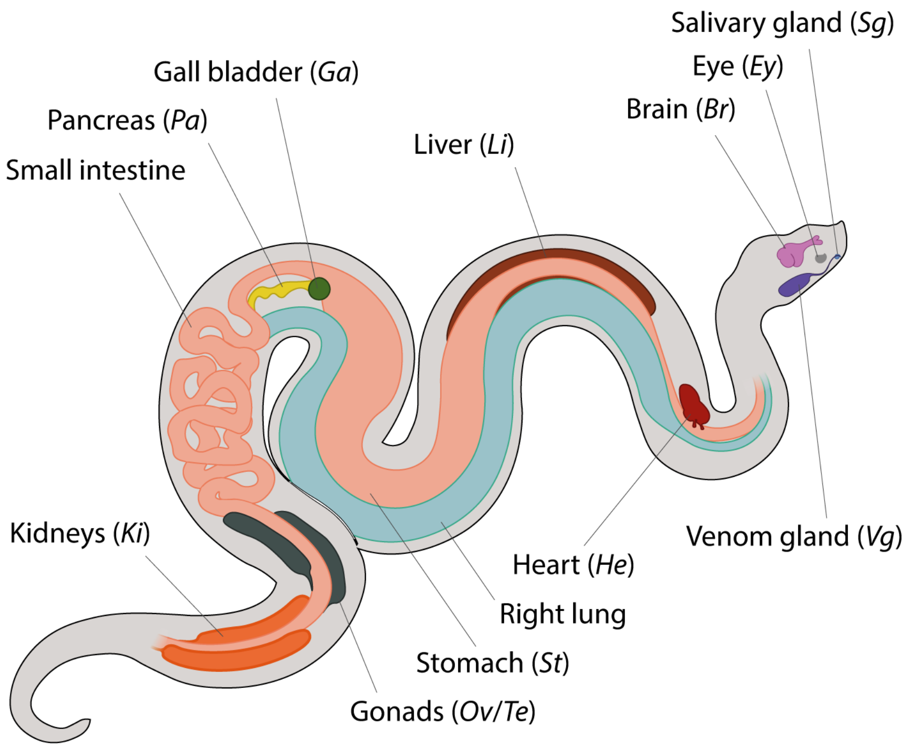


**Fig. S35. Anatomical scheme of a viper.** Tissues and organs with an abbreviation in between bracket have been used for the differential gene expression analysis shown in Fig 6A.

**Table S1. Samples (n=94) used for whole-genome sequencing (WGS).** The table shows sample codes, the lineage to which they belong, their approximate origin and their coverage when mapped to rVipLat1. First eight samples were sequenced at high coverage, and first three were used in genome assembly (CNPOR1, CNSOC1, CNMOL1).

| Sample | Species | Locality | Country | Coverage |
|---|---|---|---|---:|
| **CNPOR1** | *V. latastei* | Els Ports | Spain | 59.08 |
| **CNSOC1** | *V. seoanei* | Socobio | Spain | 50.02 |
| **CNMOL1** | *V. aspis* | La Molina | Spain | 48.96 |
| **MNCN50498** | *V. monticola* | Djurdjurá | Algeria | 48.77 |
| **CN13667** | *V. ursinii* | Kiskunság, Látó-hegy | Hungary | 44.06 |
| **SPM002443** | *V. berus* | Brandon | UK | 39.61 |
| **EMO4112020** | *V. aspis* | Prata | Italy | 39.74 |
| **CN13858** | *V. ammodytes* | Zagreb | Croatia | 41.58 |
| **ATP14** | *V. ammodytes* | Ada Bojana | Montenegro | 7.05 |
| **V6IPEAsp** | *V. aspis* | Grindenwald | Switzerland | 7.65 |
| **BEV11035** | *V. aspis* | Auzet | France | 7.33 |
| **BEV9268** | *V. aspis* | Cévennes | France | 7.00 |
| **CN19720** | *V. aspis* | Girona | Spain | 8.14 |
| **CN19737** | *V. aspis* | N Burgos | Spain | 7.60 |
| **SPM003071** | *V. aspis* | Andorra | Andorra | 6.06 |
| **CN19872** | *V. aspis* | Setcases | Spain | 6.12 |
| **CN19869** | *V. aspis* | Loza | Spain | 6.10 |
| **L58** | *V. aspis* | Alto Aneu | Spain | 5.09 |
| **L59** | *V. aspis* | Montnegre | Spain | 5.62 |
| **MALZ004** | *V. aspis* | Madonie | Italy | 6.49 |
| **E091** | *V. latastei* | Sierra Espuña | Spain | 8.04 |
| **FM14VL028** | *V. latastei* | Ciudad Real | Spain | 8.25 |
| **FM19VL036** | *V. latastei* | O Pereiro | Spain | 7.97 |
| **FM22VL206** | *V. latastei* | Santa Cruz | Portugal | 7.38 |
| **FM22VL233** | *V. latastei* | Évora | Portugal | 7.34 |
| **FM22VL238** | *V. latastei* | Faro | Portugal | 5.04 |
| **FM22VL240** | *V. latastei* | Cocharil | Portugal | 7.63 |
| **FM22VL244** | *V. latastei* | Gerês | Portugal | 9.23 |
| **CN11278** | *V. latastei* | Doñana | Spain | 6.63 |
| **CN20035** | *V. latastei* | Hervás | Spain | 6.04 |
| **CN20036** | *V. latastei* | Navezuelas | Spain | 18.93 |
| **CN20037** | *V. latastei* | Filiel | Spain | 17.86 |
| **CN12872** | *V. latastei* | Alcoi | Spain | 16.97 |
| **CN11554** | *V. latastei* | Cabo de Gata | Spain | 17.53 |
| **CN12857** | *V. latastei* | Navacerrada | Spain | 16.72 |
| **SPM003236** | *V. latastei* | Cobujón | Spain | 6.76 |
| **SPM003248** | *V. latastei* | Sierra de Loja | Spain | 6.66 |
| **SPM003249** | *V. latastei* | Sierra de Hornachuelos | Spain | 7.09 |
| **CN19763** | *V. latastei* | N Burgos | Spain | 6.58 |
| **CN13661R** | *V. latastei* | Alt Penedès | Spain | 5.28 |
| **CN19857** | *V. latastei* | Soria | Spain | 6.39 |
| **FA11Vlat** | *V. latastei* | Castellón | Spain | 7.02 |
| **SPM003079** | *V. latastei* | Peña de Francia | Spain | 6.75 |

| | | | | |
|---|---|---|---|---|
| **SPM003239R** | *V. latastei* | Carcabuey | Spain | 5.05 |
| **SPM001617** | *V. latastei* | Villaluenga del Rosario | Spain | 6.46 |
| **FM13VL026** | *V. latastei* | Montemuro | Portugal | 5.80 |
| **FM13VL024** | *V. latastei* | Peñalén | Spain | 6.14 |
| **BAR2012** | *V. latastei* | Bárdenas Reales | Spain | 6.10 |
| **LAT1** | *V. latastei* | Burgos | Spain | 6.58 |
| **CN10452R** | *V. latastei* | Garraf | Spain | 7.79 |
| **LAT2** | *V. latastei* | La Hiruela | Spain | 7.04 |
| **20VL039RR** | *V. latastei* | Vila do Conde | Portugal | 6.76 |
| **CN19858** | *V. latastei* | Moianés | Spain | 7.16 |
| **CN20038** | *V. latastei* | Tarifa | Spain | 7.01 |
| **CN20039** | *V. latastei* | Hurones | Spain | 6.94 |
| **CN20034** | *V. latastei* | Peguerinos | Spain | 6.27 |
| **CN19726** | *V. latastei x aspis* | N Burgos | Spain | 6.24 |
| **SPM002522** | *V. latastei x aspis* | Boumort | Spain | 6.62 |
| **FM22VM133** | *V. monticola* | Talasantane | Morocco | 10.76 |
| **FM22VM216** | *V. monticola* | Tamda n'Oughmar | Morocco | 9.01 |
| **FM19VM061** | *V. monticola* | Plateau du Tichka | Morocco | 6.06 |
| **FM11VM003** | *V. monticola* | Toubkal | Morocco | 5.42 |
| **FM19VM052** | *V. monticola* | Oukeimedem | Morocco | 6.26 |
| **FM16VM003** | *V. monticola* | Jebel Sirwa | Morocco | 5.80 |
| **FM13VL021** | *V. monticola* | Jebel Azourki | Morocco | 6.50 |
| **CN10736** | *V. seoanei* | Sopelana | Spain | 8.33 |
| **CN10760** | *V. seoanei* | Picos de Europa | Spain | 12.10 |
| **CN13623** | *V. seoanei* | Cistierna | Spain | 8.64 |
| **CN19710** | *V. seoanei* | Gorbea | Spain | 7.21 |
| **CN19756** | *V. seoanei* | Gorbea | Spain | 9.25 |
| **CN19984** | *V. seoanei* | Gorbea | Spain | 8.44 |
| **CN9321** | *V. seoanei* | Salientes | Spain | 7.24 |
| **FM11VS005** | *V. seoanei* | Vila Real | Portugal | 8.10 |
| **FM11VS009** | *V. seoanei* | Sanabria | Spain | 7.09 |
| **FM12VS017** | *V. seoanei* | Alto de la Garganta | Spain | 7.30 |
| **FM16VS105** | *V. seoanei* | Boñar | Spain | 6.98 |
| **FM16VS171** | *V. seoanei* | Cernedo | Spain | 7.14 |
| **FM18VS027** | *V. seoanei* | Gerês | Portugal | 7.91 |
| **FM18VS076** | *V. seoanei* | Hoces del Ebro | Spain | 6.82 |
| **FM18VS077** | *V. seoanei* | Hoces del Ebro | Spain | 6.68 |
| **FM20VS016** | *V. seoanei* | Feáns | Spain | 7.55 |
| **FM20VS130** | *V. seoanei* | Hoces del Ebro | Spain | 9.06 |
| **FM21VS016** | *V. seoanei* | Montederramo | Spain | 7.71 |
| **CN10747** | *V. seoanei* | Picos de Europa | Spain | 6.14 |
| **CN19871** | *V. seoanei* | Irati | Spain | 5.54 |
| **SPM004261R** | *V. seoanei* | Gamboa | Spain | 6.62 |
| **SPM003076R** | *V. seoanei* | Caurel | Spain | 6.33 |
| **CN10754** | *V. seoanei* | Picos de Europa | Spain | 6.75 |
| **CN19795** | *V. seoanei* | Tejedo de Ancares | Spain | 5.15 |
| **SPM004451R** | *V. seoanei* | Artxanda | Spain | 6.15 |
| **SPM004546R** | *V. seoanei* | Donostia | Spain | 5.91 |
| **SPM003072R** | *V. seoanei* | Somiedo | Spain | 6.67 |

| | | | | |
|---|---|---|---|---|
| **SPM004597R** | *V. seoanei* | Covadonga | Spain | 9.86 |
| **X034** | *V. seoanei* | Muros | Spain | 7.42 |

**Table S2. Assembly and annotation quality and completeness parameters of the three genomes assembled in this study (*V. latastei*, *V. seoanei*, and *V. aspis*) and other venomous snake reference genomes.** The genomes here presented stand out for their contig N50. BUSCO stats refer to single-copied complete orthologs of vertebrata odb10 (% out of 3,354 genes) for the assembly or to complete orthologs (single-copied or not) of vertebrata for the annotation, respectively. annot. = annotated; Gb = Gigabases; N. = number; Mb = Megabases.

| | *V. latastei* | *V seoanei* | *V. aspis* | *V. ursinii* | *C. tigris* | *N. naja* |
|---|---|---|---|---|---|---|
| Genome size (Gb) | 1.63 | 1.57 | 1.59 | 1.62 | 1.59 | 1.79 |
| N. of Scaffolds | 56 | 18 | 26 | 384 | 160 | 1,897 |
| Scaffold N50 (Mb) | 222.4 | 236.1 | 113.8 | 212.8 | 207.7 | 223.4 |
| Scaffold L50 | 3 | 3 | 6 | 3 | - | - |
| Contig N50 (Mb) | 44.84 | 75.89 | 35.32 | 2.13 | 2.11 | 0.3 |
| Assembly BUSCO (%) | 97.1 | 96.2 | 96.1 | - | - | - |
| GC% | 40.96 | 40.94 | 40.96 | 40.93 | 39.8 | 40.5 |
| Structurally annot. genes | 59,409 | 61,622 | 57,979 | - | - | - |
| Annotation BUSCO (%) | 98.6 | 97.8 | 97.6 | | | |
| Functionally annot. genes | 26,312 | 25,391 | 26,213 | - | - | - |

**Table S3. Introgression results from ABBA-BABA tests implemented on Dsuite.** Results are obtained from a dataset of 86 samples, comprising all studied *Vipera* lineages and excluding individuals from secondary contacts (see Admixture results), which yielded a total of 12,645,411 SNPs. Only trios showing significant results regarding the robust clustering threshold (*p-value* < 0.001) and Z-score |Z| > 3 are shown. Sensitive and robust clustering *p-values* are calculated based on Kolmogorov-Smirnov tests to correct for homoplasies. amm stands for *V. ammodytes*; ber for *V. berus*; urs for *V. ursinii*; aspasp for *V. aspis aspis*; aspzin for *V. aspis zinnikeri*; asphug for *V. aspis hugyi*; aspfra for *V. aspis francisciredi*; latlat for *V. latastei latastei*; latgad for *V. latastei gaditana*; lataru for *V. latastei arundana*; monN for the Northern cluster of *V. monticola*; monS for the Southern cluster of *V. monticola*; seoE for the Eastern cluster of *V. seoanei* and seoW for the Western cluster of the same species (see Admixture results).

| P1 | P2 | P3 | Dstatistic | Z-score | *p-value* | *f4-ratio* | Sensitive clustering | Robust clustering | BBAA | ABBA | BABA |
|---|---|---|---|---|---|---|---|---|---|---|---|
| amm | aspasp | ber | 0.217 | 26.854 | <0.0001 | 0.1 | <0.0001 | 0.0004 | 219428 | 240074 | 154569 |
| amm | aspasp | latgad | 0.022 | 3.186 | 0.0014 | 0.009 | <0.0001 | 0.0004 | 299566 | 167236 | 159967 |
| amm | aspasp | latlat | 0.043 | 5.248 | <0.0001 | 0.017 | <0.0001 | 0.0001 | 297957 | 172900 | 158719 |
| amm | aspasp | seoE | 0.197 | 19.04 | <0.0001 | 0.08 | <0.0001 | <0.0001 | 259147 | 236950 | 159034 |
| amm | aspasp | seoW | 0.162 | 20.281 | <0.0001 | 0.074 | <0.0001 | <0.0001 | 264977 | 223004 | 160763 |
| amm | aspfra | ber | 0.132 | 15.05 | <0.0001 | 0.054 | <0.0001 | 0.0002 | 219381 | 192091 | 147371 |
| amm | aspfra | seoE | 0.125 | 15.147 | <0.0001 | 0.045 | <0.0001 | <0.0001 | 256252 | 192842 | 149857 |
| amm | aspfra | seoW | 0.101 | 13.694 | <0.0001 | 0.042 | <0.0001 | <0.0001 | 260742 | 184319 | 150377 |
| amm | aspfra | urs | 0.082 | 7.907 | <0.0001 | 0.045 | <0.0001 | <0.0001 | 211949 | 172856 | 146792 |

| | | | | | | | | | | | |
|---|---|---|---|---|---|---|---|---|---|---|---|
| amm | asphug | ber | 0.123 | 13.389 | <0.0001 | 0.05 | <0.0001 | <0.0001 | 217180 | 186356 | 145472 |
| amm | asphug | seoE | 0.118 | 14.361 | <0.0001 | 0.042 | <0.0001 | 0.0005 | 253650 | 187715 | 147975 |
| amm | asphug | seoW | 0.095 | 12.714 | <0.0001 | 0.039 | <0.0001 | 0.0004 | 257921 | 179610 | 148376 |
| amm | asphug | urs | 0.073 | 6.777 | <0.0001 | 0.04 | <0.0001 | <0.0001 | 209998 | 168024 | 145027 |
| amm | aspzin | ber | 0.271 | 23.428 | <0.0001 | 0.136 | <0.0001 | <0.0001 | 203384 | 271894 | 156007 |
| amm | aspzin | lataru | 0.157 | 8.393 | <0.0001 | 0.062 | <0.0001 | <0.0001 | 277905 | 211905 | 154294 |
| amm | aspzin | latgad | 0.17 | 9.009 | <0.0001 | 0.079 | <0.0001 | <0.0001 | 275760 | 217314 | 154219 |
| amm | aspzin | latlat | 0.192 | 9.514 | <0.0001 | 0.086 | <0.0001 | <0.0001 | 273771 | 225299 | 152596 |
| amm | aspzin | monS | 0.142 | 8.014 | <0.0001 | 0.05 | <0.0001 | <0.0001 | 279242 | 206172 | 154847 |
| amm | aspzin | monN | 0.142 | 8.021 | <0.0001 | 0.05 | <0.0001 | <0.0001 | 276853 | 205616 | 154482 |
| amm | aspzin | seoE | 0.344 | 18.115 | <0.0001 | 0.162 | <0.0001 | <0.0001 | 233204 | 309966 | 151168 |
| amm | aspzin | seoW | 0.312 | 18.243 | <0.0001 | 0.166 | <0.0001 | <0.0001 | 239319 | 292256 | 153181 |
| amm | aspzin | urs | 0.121 | 14.413 | <0.0001 | 0.076 | <0.0001 | <0.0001 | 203507 | 207862 | 162889 |
| amm | ber | monS | 0.156 | 23.472 | <0.0001 | 0.057 | <0.0001 | 0.0006 | 226440 | 210182 | 153305 |
| amm | ber | monN | 0.156 | 23.902 | <0.0001 | 0.057 | <0.0001 | 0.0003 | 224317 | 209582 | 153036 |
| amm | seoE | lataru | 0.438 | 73.146 | <0.0001 | 0.247 | <0.0001 | <0.0001 | 187523 | 373556 | 145931 |
| amm | seoW | lataru | 0.467 | 79.751 | <0.0001 | 0.271 | <0.0001 | <0.0001 | 180167 | 392817 | 142677 |
| amm | seoE | latgad | 0.466 | 79.509 | <0.0001 | 0.313 | <0.0001 | <0.0001 | 183034 | 394159 | 143528 |
| amm | seoW | latgad | 0.493 | 86.412 | <0.0001 | 0.341 | <0.0001 | <0.0001 | 175786 | 413332 | 140384 |
| amm | seoE | latlat | 0.447 | 77.089 | <0.0001 | 0.279 | <0.0001 | 0.0002 | 185118 | 381503 | 145978 |
| amm | seoW | latlat | 0.473 | 82.9 | <0.0001 | 0.304 | <0.0001 | <0.0001 | 178057 | 399779 | 143021 |
| amm | seoE | monS | 0.416 | 65.352 | <0.0001 | 0.204 | <0.0001 | <0.0001 | 190560 | 358995 | 148199 |
| amm | seoW | monS | 0.445 | 72.423 | <0.0001 | 0.226 | <0.0001 | <0.0001 | 183221 | 377847 | 144962 |
| amm | seoE | monN | 0.416 | 65.754 | <0.0001 | 0.207 | <0.0001 | <0.0001 | 188345 | 358276 | 147812 |
| amm | seoW | monN | 0.446 | 72.621 | <0.0001 | 0.229 | <0.0001 | <0.0001 | 181033 | 377037 | 144601 |
| seoE | urs | amm | 0.123 | 20.921 | <0.0001 | 0.085 | <0.0001 | <0.0001 | 283681 | 174271 | 136017 |
| seoW | urs | amm | 0.136 | 22.158 | <0.0001 | 0.093 | <0.0001 | <0.0001 | 282854 | 176143 | 133974 |
| asphug | aspfra | aspasp | 0.08 | 20.694 | <0.0001 | 0.035 | <0.0001 | 0.0043 | 302166 | 90602.1 | 77169 |
| aspzin | aspasp | aspfra | 0.222 | 15.876 | <0.0001 | 0.162 | <0.0001 | <0.0001 | 317264 | 194230 | 123727 |
| aspfra | aspasp | ber | 0.149 | 10.812 | <0.0001 | 0.049 | <0.0001 | 0.0006 | 512130 | 147663 | 109444 |
| aspfra | aspasp | seoE | 0.127 | 9.542 | <0.0001 | 0.036 | <0.0001 | 0.0014 | 561441 | 144528 | 111851 |
| aspfra | aspasp | seoW | 0.107 | 8.59 | <0.0001 | 0.034 | <0.0001 | 0.003 | 573695 | 137513 | 111029 |
| aspfra | aspasp | urs | 0.058 | 7.209 | <0.0001 | 0.023 | <0.0001 | 0.0067 | 515976 | 118702 | 105760 |
| aspzin | aspasp | asphug | 0.218 | 15.849 | <0.0001 | 0.142 | <0.0001 | <0.0001 | 324578 | 188829 | 121220 |
| asphug | aspasp | ber | 0.159 | 10.905 | <0.0001 | 0.053 | <0.0001 | 0.0024 | 496737 | 149799 | 108702 |
| asphug | aspasp | seoE | 0.136 | 9.889 | <0.0001 | 0.039 | <0.0001 | 0.0069 | 545030 | 146705 | 111492 |
| asphug | aspasp | seoW | 0.114 | 9.035 | <0.0001 | 0.037 | <0.0001 | 0.0056 | 556777 | 139457 | 110815 |
| aspasp | aspzin | lataru | 0.237 | 13.379 | <0.0001 | 0.058 | <0.0001 | <0.0001 | 760719 | 140788 | 86807.1 |
| aspasp | aspzin | latgad | 0.243 | 13.655 | <0.0001 | 0.07 | <0.0001 | <0.0001 | 756182 | 143706 | 87604.4 |
| aspasp | aspzin | latlat | 0.25 | 13.716 | <0.0001 | 0.07 | <0.0001 | <0.0001 | 749671 | 147201 | 88393 |
| aspasp | aspzin | monS | 0.226 | 13.237 | <0.0001 | 0.049 | <0.0001 | <0.0001 | 764227 | 137123 | 86592.2 |
| aspasp | aspzin | monN | 0.227 | 13.356 | <0.0001 | 0.05 | <0.0001 | <0.0001 | 760875 | 136837 | 86254.4 |
| aspasp | aspzin | seoE | 0.282 | 17.119 | <0.0001 | 0.09 | <0.0001 | <0.0001 | 661396 | 184639 | 103430 |

| | | | | | | | | | | | |
|---|---|---|---|---|---|---|---|---|---|---|---|
| aspasp | aspzin | seoW | 0.277 | 16.863 | <0.0001 | 0.099 | <0.0001 | <0.0001 | 678767 | 178070 | 100915 |
| aspasp | ber | lataru | 0.171 | 19.351 | <0.0001 | 0.072 | <0.0001 | 0.0013 | 312272 | 222328 | 157479 |
| aspasp | ber | latgad | 0.196 | 22.257 | <0.0001 | 0.099 | <0.0001 | 0.0008 | 306233 | 233096 | 156769 |
| aspasp | ber | latlat | 0.171 | 18.85 | <0.0001 | 0.083 | <0.0001 | 0.0013 | 304790 | 229586 | 162444 |
| aspasp | ber | monS | 0.151 | 16.806 | <0.0001 | 0.056 | <0.0001 | 0.0009 | 316639 | 214461 | 158307 |
| aspasp | ber | monN | 0.151 | 17.229 | <0.0001 | 0.057 | <0.0001 | 0.0017 | 314296 | 213871 | 157818 |
| aspasp | seoE | lataru | 0.427 | 58.956 | <0.0001 | 0.243 | <0.0001 | <0.0001 | 266972 | 375728 | 150909 |
| aspasp | seoW | lataru | 0.452 | 67.645 | <0.0001 | 0.268 | <0.0001 | <0.0001 | 245961 | 397186 | 149779 |
| aspasp | urs | lataru | 0.09 | 11.038 | <0.0001 | 0.036 | <0.0001 | <0.0001 | 276827 | 193687 | 161832 |
| aspasp | seoE | latgad | 0.45 | 61.71 | <0.0001 | 0.306 | <0.0001 | <0.0001 | 260056 | 393983 | 149445 |
| aspasp | seoW | latgad | 0.473 | 70.421 | <0.0001 | 0.334 | <0.0001 | <0.0001 | 239187 | 415389 | 148461 |
| aspasp | urs | latgad | 0.094 | 11.615 | <0.0001 | 0.044 | <0.0001 | <0.0001 | 273138 | 197332 | 163359 |
| aspasp | seoE | latlat | 0.414 | 55.069 | <0.0001 | 0.266 | <0.0001 | <0.0001 | 260777 | 379907 | 157464 |
| aspasp | seoW | latlat | 0.437 | 60.683 | <0.0001 | 0.292 | <0.0001 | <0.0001 | 240259 | 400579 | 156833 |
| aspasp | urs | latlat | 0.072 | 8.032 | <0.0001 | 0.033 | <0.0001 | <0.0001 | 271764 | 195229 | 169070 |
| aspasp | seoE | monS | 0.409 | 54.169 | <0.0001 | 0.204 | <0.0001 | <0.0001 | 272056 | 363127 | 152373 |
| aspasp | seoW | monS | 0.435 | 62.634 | <0.0001 | 0.225 | <0.0001 | <0.0001 | 251017 | 384135 | 151219 |
| aspasp | urs | monS | 0.084 | 10.892 | <0.0001 | 0.03 | <0.0001 | <0.0001 | 279544 | 190410 | 160963 |
| aspasp | seoE | monN | 0.41 | 55.366 | <0.0001 | 0.207 | <0.0001 | <0.0001 | 269674 | 362430 | 151771 |
| aspasp | seoW | monN | 0.436 | 64.107 | <0.0001 | 0.228 | <0.0001 | <0.0001 | 248701 | 383349 | 150651 |
| aspasp | urs | monN | 0.084 | 11.159 | <0.0001 | 0.03 | <0.0001 | 0.0002 | 277206 | 189860 | 160391 |
| asphug | aspfra | aspzin | 0.073 | 18.345 | <0.0001 | 0.03 | <0.0001 | 0.0037 | 362513 | 83084.7 | 71735.4 |
| aspfra | aspzin | ber | 0.217 | 13.554 | <0.0001 | 0.086 | <0.0001 | <0.0001 | 456421 | 189871 | 122092 |
| aspfra | aspzin | lataru | 0.208 | 9.267 | <0.0001 | 0.062 | <0.0001 | <0.0001 | 568568 | 162126 | 106224 |
| aspfra | aspzin | latgad | 0.218 | 9.615 | <0.0001 | 0.077 | <0.0001 | <0.0001 | 565194 | 166041 | 106522 |
| aspfra | aspzin | latlat | 0.235 | 9.978 | <0.0001 | 0.08 | <0.0001 | <0.0001 | 560835 | 171382 | 106135 |
| aspfra | aspzin | monS | 0.193 | 9.002 | <0.0001 | 0.051 | <0.0001 | <0.0001 | 570838 | 157427 | 106598 |
| aspfra | aspzin | monN | 0.193 | 9.1 | <0.0001 | 0.051 | <0.0001 | <0.0001 | 567936 | 157096 | 106203 |
| aspfra | aspzin | seoE | 0.327 | 13.553 | <0.0001 | 0.122 | <0.0001 | <0.0001 | 493700 | 225942 | 114494 |
| aspfra | aspzin | seoW | 0.308 | 13.155 | <0.0001 | 0.13 | <0.0001 | <0.0001 | 506119 | 215162 | 113832 |
| aspfra | ber | lataru | 0.178 | 26.057 | <0.0001 | 0.076 | <0.0001 | 0.0079 | 266750 | 217633 | 151755 |
| aspfra | ber | latgad | 0.207 | 30.714 | <0.0001 | 0.104 | <0.0001 | 0.0044 | 261869 | 229096 | 150548 |
| aspfra | ber | latlat | 0.191 | 28.606 | <0.0001 | 0.092 | <0.0001 | 0.0083 | 261255 | 226497 | 153790 |
| aspfra | ber | monS | 0.154 | 21.163 | <0.0001 | 0.057 | <0.0001 | 0.0054 | 269957 | 208946 | 153172 |
| aspfra | ber | monN | 0.154 | 21.805 | <0.0001 | 0.058 | <0.0001 | 0.0092 | 267774 | 208368 | 152707 |
| aspfra | seoE | lataru | 0.43 | 61.276 | <0.0001 | 0.246 | <0.0001 | 0.0011 | 229416 | 367136 | 146206 |
| aspfra | seoW | lataru | 0.457 | 70.105 | <0.0001 | 0.27 | <0.0001 | <0.0001 | 214371 | 387048 | 144240 |
| aspfra | urs | lataru | 0.101 | 18.478 | <0.0001 | 0.039 | <0.0001 | 0.0003 | 252666 | 184948 | 151019 |
| aspfra | seoE | latgad | 0.456 | 65.405 | <0.0001 | 0.311 | <0.0001 | 0.0007 | 223712 | 385898 | 144246 |
| aspfra | seoW | latgad | 0.48 | 74.379 | <0.0001 | 0.339 | <0.0001 | <0.0001 | 208823 | 405779 | 142438 |
| aspfra | urs | latgad | 0.11 | 20.982 | <0.0001 | 0.05 | <0.0001 | 0.0001 | 249561 | 189003 | 151493 |
| aspfra | seoE | latlat | 0.427 | 61.395 | <0.0001 | 0.274 | <0.0001 | 0.0057 | 225074 | 372889 | 149579 |
| aspfra | seoW | latlat | 0.452 | 68.798 | <0.0001 | 0.299 | <0.0001 | <0.0001 | 210466 | 391986 | 148053 |

| | | | | | | | | | | | |
|---|---|---|---|---|---|---|---|---|---|---|---|
| aspfra | urs | latlat | 0.096 | 17.169 | <0.0001 | 0.042 | <0.0001 | 0.001 | 248871 | 187631 | 154673 |
| aspfra | seoE | monS | 0.41 | 56.24 | <0.0001 | 0.205 | <0.0001 | 0.0008 | 233230 | 353759 | 148187 |
| aspfra | seoW | monS | 0.437 | 64.942 | <0.0001 | 0.226 | <0.0001 | <0.0001 | 218217 | 373304 | 146254 |
| aspfra | urs | monS | 0.09 | 16.006 | <0.0001 | 0.031 | <0.0001 | 0.0002 | 254601 | 181034 | 151117 |
| aspfra | seoE | monN | 0.41 | 57.418 | <0.0001 | 0.208 | <0.0001 | 0.0003 | 231007 | 353073 | 147578 |
| aspfra | seoW | monN | 0.438 | 66.161 | <0.0001 | 0.229 | <0.0001 | <0.0001 | 216046 | 372524 | 145679 |
| aspfra | urs | monN | 0.09 | 16.789 | <0.0001 | 0.032 | <0.0001 | 0.0006 | 252403 | 180551 | 150591 |
| asphug | aspzin | ber | 0.226 | 13.879 | <0.0001 | 0.09 | <0.0001 | <0.0001 | 443276 | 191001 | 120694 |
| asphug | aspzin | lataru | 0.206 | 9.392 | <0.0001 | 0.062 | <0.0001 | <0.0001 | 551842 | 161725 | 106560 |
| asphug | aspzin | latgad | 0.216 | 9.713 | <0.0001 | 0.076 | <0.0001 | <0.0001 | 548561 | 165635 | 106856 |
| asphug | aspzin | latlat | 0.232 | 9.993 | <0.0001 | 0.08 | <0.0001 | <0.0001 | 544299 | 171004 | 106581 |
| asphug | aspzin | monS | 0.189 | 9.036 | <0.0001 | 0.05 | <0.0001 | <0.0001 | 553884 | 156895 | 106998 |
| asphug | aspzin | monN | 0.189 | 9.079 | <0.0001 | 0.051 | <0.0001 | <0.0001 | 550999 | 156487 | 106720 |
| asphug | aspzin | seoE | 0.332 | 13.72 | <0.0001 | 0.125 | <0.0001 | <0.0001 | 479814 | 226759 | 113773 |
| asphug | aspzin | seoW | 0.312 | 13.401 | <0.0001 | 0.132 | <0.0001 | <0.0001 | 491744 | 215796 | 113275 |
| asphug | ber | latgad | 0.207 | 31.052 | <0.0001 | 0.104 | <0.0001 | 0.0067 | 255618 | 226631 | 149035 |
| asphug | ber | monS | 0.153 | 21.146 | <0.0001 | 0.057 | <0.0001 | 0.0062 | 263370 | 206495 | 151717 |
| asphug | ber | monN | 0.153 | 21.731 | <0.0001 | 0.057 | <0.0001 | 0.0082 | 261160 | 205858 | 151363 |
| asphug | seoE | lataru | 0.429 | 61.358 | <0.0001 | 0.246 | <0.0001 | 0.0002 | 224174 | 362952 | 144858 |
| asphug | seoW | lataru | 0.456 | 70.749 | <0.0001 | 0.27 | <0.0001 | <0.0001 | 209738 | 382645 | 142850 |
| asphug | urs | lataru | 0.101 | 19.066 | <0.0001 | 0.039 | <0.0001 | 0.0012 | 246698 | 182799 | 149366 |
| asphug | seoE | latgad | 0.455 | 65.746 | <0.0001 | 0.31 | <0.0001 | 0.0001 | 218625 | 381552 | 142961 |
| asphug | seoW | latgad | 0.48 | 75.14 | <0.0001 | 0.338 | <0.0001 | <0.0001 | 204332 | 401223 | 141096 |
| asphug | urs | latgad | 0.11 | 21.744 | <0.0001 | 0.05 | <0.0001 | 0.0009 | 243740 | 186848 | 149864 |
| asphug | seoE | latlat | 0.426 | 61.296 | <0.0001 | 0.273 | <0.0001 | 0.0009 | 219911 | 368659 | 148235 |
| asphug | seoW | latlat | 0.451 | 69.18 | <0.0001 | 0.298 | <0.0001 | <0.0001 | 205884 | 387558 | 146635 |
| asphug | urs | latlat | 0.096 | 17.94 | <0.0001 | 0.042 | <0.0001 | 0.0051 | 243026 | 185481 | 153044 |
| asphug | seoE | monS | 0.408 | 56.031 | <0.0001 | 0.204 | <0.0001 | 0.0001 | 227822 | 349666 | 146967 |
| asphug | seoW | monS | 0.436 | 65.016 | <0.0001 | 0.225 | <0.0001 | <0.0001 | 213404 | 368933 | 144976 |
| asphug | urs | monS | 0.089 | 16.364 | <0.0001 | 0.031 | <0.0001 | 0.0011 | 248527 | 178837 | 149649 |
| asphug | seoE | monN | 0.409 | 57.105 | <0.0001 | 0.207 | <0.0001 | <0.0001 | 225575 | 348892 | 146490 |
| asphug | seoW | monN | 0.436 | 66.213 | <0.0001 | 0.229 | <0.0001 | <0.0001 | 211198 | 368061 | 144517 |
| asphug | urs | monN | 0.089 | 16.835 | <0.0001 | 0.031 | <0.0001 | 0.0016 | 246289 | 178259 | 149225 |
| seoW | ber | aspzin | 0.046 | 4.775 | <0.0001 | 0.025 | <0.0001 | <0.0001 | 420215 | 188751 | 171994 |
| urs | ber | aspzin | 0.179 | 13.989 | <0.0001 | 0.089 | <0.0001 | <0.0001 | 274862 | 203821 | 141893 |
| aspzin | seoE | lataru | 0.341 | 24.868 | <0.0001 | 0.196 | <0.0001 | 0.0002 | 308281 | 335871 | 164929 |
| aspzin | seoW | lataru | 0.371 | 28.897 | <0.0001 | 0.222 | <0.0001 | <0.0001 | 283212 | 357342 | 163805 |
| aspzin | seoE | latgad | 0.365 | 26.781 | <0.0001 | 0.254 | <0.0001 | 0.0002 | 299738 | 352496 | 163949 |
| aspzin | seoW | latgad | 0.393 | 30.733 | <0.0001 | 0.284 | <0.0001 | 0.0002 | 274857 | 373963 | 163020 |
| aspzin | seoE | latlat | 0.32 | 20.642 | <0.0001 | 0.211 | <0.0001 | <0.0001 | 299998 | 337959 | 174222 |
| aspzin | seoW | latlat | 0.348 | 23.301 | <0.0001 | 0.238 | <0.0001 | <0.0001 | 275518 | 358740 | 173694 |
| urs | aspzin | latlat | 0.077 | 3.562 | 0.0004 | 0.039 | <0.0001 | <0.0001 | 262625 | 211774 | 181420 |
| aspzin | seoE | monS | 0.327 | 24.752 | <0.0001 | 0.163 | <0.0001 | <0.0001 | 315462 | 325350 | 165028 |

| | | | | | | | | | | | |
|---|---|---|---|---|---|---|---|---|---|---|---|
| aspzin | seoW | monS | 0.358 | 28.945 | <0.0001 | 0.185 | <0.0001 | <0.0001 | 290397 | 346402 | 163913 |
| aspzin | seoE | monN | 0.327 | 25.059 | <0.0001 | 0.165 | <0.0001 | <0.0001 | 312909 | 324687 | 164512 |
| aspzin | seoW | monN | 0.358 | 29.404 | <0.0001 | 0.188 | <0.0001 | <0.0001 | 287914 | 345641 | 163423 |
| urs | seoE | aspzin | 0.16 | 7.697 | <0.0001 | 0.097 | <0.0001 | <0.0001 | 248267 | 253347 | 183463 |
| urs | seoW | aspzin | 0.11 | 6.254 | <0.0001 | 0.065 | <0.0001 | <0.0001 | 251153 | 235940 | 189065 |
| ber | seoE | lataru | 0.404 | 79.096 | <0.0001 | 0.184 | <0.0001 | 0.0005 | 468788 | 265888 | 112856 |
| ber | seoW | lataru | 0.446 | 87.188 | <0.0001 | 0.21 | <0.0001 | <0.0001 | 460410 | 283522 | 108548 |
| urs | ber | lataru | 0.112 | 28.187 | <0.0001 | 0.038 | <0.0001 | 0.0081 | 342919 | 156203 | 124748 |
| monS | latgad | ber | 0.198 | 41.672 | <0.0001 | 0.033 | <0.0001 | <0.0001 | 755316 | 87851.4 | 58792.9 |
| monN | latgad | ber | 0.191 | 39.968 | <0.0001 | 0.032 | <0.0001 | 0.0009 | 754204 | 85822 | 58273.4 |
| ber | seoE | latgad | 0.411 | 85.718 | <0.0001 | 0.23 | <0.0001 | <0.0001 | 454526 | 276102 | 115361 |
| ber | seoW | latgad | 0.451 | 93.985 | <0.0001 | 0.261 | <0.0001 | <0.0001 | 446236 | 293634 | 111143 |
| ber | seoE | latlat | 0.387 | 78.584 | <0.0001 | 0.2 | <0.0001 | 0.004 | 458924 | 266179 | 117697 |
| ber | seoW | latlat | 0.427 | 84.381 | <0.0001 | 0.228 | <0.0001 | <0.0001 | 450893 | 282908 | 113737 |
| ber | seoE | monS | 0.4 | 76.057 | <0.0001 | 0.156 | <0.0001 | 0.0012 | 479224 | 259054 | 111009 |
| ber | seoW | monS | 0.443 | 85.564 | <0.0001 | 0.179 | <0.0001 | <0.0001 | 470845 | 276288 | 106700 |
| urs | ber | monS | 0.093 | 22.362 | <0.0001 | 0.027 | <0.0001 | 0.0044 | 348502 | 149941 | 124505 |
| ber | seoE | monN | 0.401 | 77.246 | <0.0001 | 0.159 | <0.0001 | 0.0011 | 476431 | 258584 | 110509 |
| ber | seoW | monN | 0.444 | 86.228 | <0.0001 | 0.181 | <0.0001 | <0.0001 | 468094 | 275741 | 106245 |
| urs | ber | monN | 0.093 | 22.399 | <0.0001 | 0.027 | <0.0001 | 0.0028 | 346143 | 149462 | 124134 |
| seoW | ber | urs | 0.23 | 47.053 | <0.0001 | 0.152 | <0.0001 | 0.0009 | 345398 | 193518 | 121182 |
| seoE | seoW | lataru | 0.08 | 17.316 | <0.0001 | 0.032 | <0.0001 | <0.0001 | 751170 | 151809 | 129214 |
| urs | seoE | lataru | 0.42 | 74.83 | <0.0001 | 0.215 | <0.0001 | <0.0001 | 279812 | 311067 | 127195 |
| urs | seoW | lataru | 0.455 | 83.03 | <0.0001 | 0.24 | <0.0001 | <0.0001 | 273059 | 328718 | 123155 |
| latlat | latgad | seoW | 0.076 | 17.983 | <0.0001 | 0.026 | <0.0001 | 0.0011 | 618020 | 111763 | 95878.6 |
| monS | latgad | urs | 0.109 | 38.052 | <0.0001 | 0.021 | <0.0001 | 0.0014 | 769813 | 67797.3 | 54459.8 |
| seoE | seoW | latgad | 0.077 | 17.721 | <0.0001 | 0.04 | <0.0001 | <0.0001 | 730254 | 155731 | 133335 |
| urs | seoE | latgad | 0.442 | 82.269 | <0.0001 | 0.274 | <0.0001 | <0.0001 | 271949 | 327377 | 126664 |
| urs | seoW | latgad | 0.475 | 90.582 | <0.0001 | 0.303 | <0.0001 | <0.0001 | 265292 | 344937 | 122723 |
| monS | latlat | seoE | 0.145 | 25.832 | <0.0001 | 0.034 | <0.0001 | 0.0075 | 628460 | 111099 | 83029.5 |
| monN | latlat | seoE | 0.136 | 24.965 | <0.0001 | 0.032 | <0.0001 | 0.003 | 627109 | 108800 | 82673.3 |
| seoE | seoW | latlat | 0.075 | 16.509 | <0.0001 | 0.035 | <0.0001 | <0.0001 | 742208 | 151801 | 130491 |
| urs | seoE | latlat | 0.421 | 78.826 | <0.0001 | 0.242 | <0.0001 | 0.0006 | 274792 | 316048 | 128783 |
| urs | seoW | latlat | 0.454 | 86.331 | <0.0001 | 0.268 | <0.0001 | <0.0001 | 268349 | 332769 | 125055 |
| monS | monN | seoW | 0.028 | 7.311 | <0.0001 | 0.003 | <0.0001 | 0.0062 | 769088 | 32012.8 | 30245.8 |
| seoE | seoW | monS | 0.081 | 17.23 | <0.0001 | 0.027 | <0.0001 | <0.0001 | 766143 | 148539 | 126372 |
| urs | seoE | monS | 0.404 | 66.375 | <0.0001 | 0.179 | <0.0001 | 0.0001 | 286092 | 300293 | 127474 |
| urs | seoW | monS | 0.44 | 77.228 | <0.0001 | 0.201 | <0.0001 | <0.0001 | 279361 | 317539 | 123460 |
| seoE | seoW | monN | 0.08 | 17.264 | <0.0001 | 0.027 | <0.0001 | <0.0001 | 762522 | 148121 | 126077 |
| urs | seoE | monN | 0.405 | 67.442 | <0.0001 | 0.182 | <0.0001 | <0.0001 | 283696 | 299641 | 126964 |
| urs | seoW | monN | 0.441 | 77.969 | <0.0001 | 0.204 | <0.0001 | <0.0001 | 276990 | 316808 | 122972 |

**Table S4. *f-branch* statistics for the introgression analysis with 86 *Vipera* samples and sensitive clustering threshold.** *f-branch* values are computed only with trios whose Dstatistic *p-value* < 0.001, sensitive clustering threshold *p-value* < 0.001 and Z-score |Z| > 3. Lineage codes are the same as in Table S3. Dashes indicate comparisons that cannot be made. Fig. 4A shows the placement of the branches (bX) in the species tree.

| branch | branch_descendants | Outgroup | latlat | latgad | lataru | monN | monS | amm | aspzin | aspasp | aspfra | asphug | urs | berus | seoW | seoE |
|---|---|---|---|---|---|---|---|---|---|---|---|---|---|---|---|---|
| b2 | amm, asp, urs, berus, seo | - | - | - | - | - | - | - | - | - | - | - | - | - | - | - |
| b3 | mon, lat | - | - | - | - | - | - | - | - | - | - | - | - | - | - | - |
| b4 | amm, asp | - | 0 | 0 | 0 | 0 | 0 | - | - | - | - | - | - | - | - | - |
| b5 | urs, berus, seo | - | 0.042 | 0.05 | 0.039 | 0.031 | 0.03 | - | - | - | - | - | - | - | - | - |
| b6 | lat | - | - | - | - | - | - | 0 | 0.008 | 0.003 | 0.001 | 0.001 | 0.008 | 0.013 | 0.027 | 0.021 |
| b7 | mon | - | - | - | - | - | - | 0 | 0 | 0 | 0 | 0 | 0 | 0 | 0 | 0 |
| b8 | amm | - | 0 | 0 | 0 | 0 | 0 | - | - | - | - | - | 0 | 0 | 0 | 0 |
| b9 | asp | - | 0.007 | 0 | 0 | 0 | 0 | - | - | - | - | - | 0.04 | 0.05 | 0.039 | 0.042 |
| b10 | urs | - | 0 | 0 | 0 | 0 | 0 | 0.085 | 0 | 0 | 0.028 | 0.026 | - | - | - | - |
| b11 | berus, seo | - | 0.052 | 0.057 | 0.038 | 0.027 | 0.027 | 0 | 0.065 | 0 | 0 | 0 | - | - | - | - |
| b12 | latgad, lataru | - | - | - | - | 0.075 | 0.07 | 0 | 0 | 0 | 0 | 0 | 0 | 0 | 0 | 0 |
| b13 | latlat | - | - | - | - | 0 | 0 | 0.002 | 0.017 | 0.011 | 0.007 | 0.006 | 0.005 | 0.009 | 0.007 | 0.006 |
| b14 | monS | - | 0 | 0 | 0 | - | - | 0 | 0 | 0 | 0 | 0 | 0 | 0 | 0 | 0 |
| b15 | monN | - | 0.02 | 0.073 | 0.023 | - | - | 0.003 | 0.002 | 0 | 0 | 0.002 | 0 | 0 | 0.003 | 0.002 |
| b16 | aspasp, aspzin | - | 0.01 | 0 | 0 | 0 | 0 | 0 | - | - | - | - | 0.026 | 0.051 | 0.035 | 0.037 |
| b17 | aspfra, asphug | - | 0 | 0 | 0 | 0 | 0 | 0.049 | - | - | - | - | 0 | 0 | 0 | 0 |
| b18 | seo | - | 0.2 | 0.23 | 0.184 | 0.159 | 0.156 | 0 | 0 | 0 | 0 | 0 | 0 | - | - | - |
| b19 | berus | - | 0 | 0 | 0 | 0 | 0 | 0.075 | 0.012 | 0.063 | 0.051 | 0.047 | 0.15 | - | - | - |
| b20 | latgad | - | 0 | - | - | 0 | 0 | 0 | 0 | 0 | 0.004 | 0.004 | 0.012 | 0.02 | 0.04 | 0 |
| b21 | lataru | - | 0.059 | - | - | 0.148 | 0.138 | 0 | 0 | 0 | 0 | 0 | 0 | 0 | 0 | 0 |
| b22 | aspasp | - | 0 | 0 | 0 | 0 | 0 | 0.047 | - | - | 0.162 | 0.142 | 0 | 0 | 0 | 0 |
| b23 | aspzin | - | 0.07 | 0.07 | 0.058 | 0.05 | 0.049 | 0 | - | - | 0 | 0 | 0.008 | 0.04 | 0.099 | 0.09 |
| b24 | aspfra | - | 0 | 0 | 0 | 0 | 0 | 0 | 0.03 | 0.035 | - | - | 0.005 | 0.005 | 0.003 | 0.003 |
| b25 | asphug | - | 0 | 0 | 0 | 0 | 0 | 0 | 0 | 0 | - | - | 0 | 0 | 0 | 0 |
| b26 | seoW | - | 0.035 | 0.04 | 0.032 | 0.027 | 0.027 | 0 | 0 | 0 | 0 | 0 | 0 | 0 | - | - |
| b27 | seoE | - | 0 | 0 | 0 | 0 | 0 | 0.009 | 0.034 | 0.023 | 0.016 | 0.015 | 0.006 | 0.008 | - | - |

**Table S5. *f-branch* statistics for the introgression analysis with 86 *Vipera* samples and robust clustering threshold.** *f-branch* values are computed only with trios whose Dstatistic *p-value* < 0.001, robust clustering threshold *p-value* < 0.001 and Z-score |Z| > 3. Lineage codes are the same as in Table S3. Dashes indicate comparisons that cannot be made. Fig. 4A shows the placement of the branches (bX) in the species tree.

| branch | branch_descendants | Outgroup | latlat | latgad | lataru | monN | monS | amm | aspzin | aspasp | aspfra | asphug | urs | berus | seoW | seoE |
|---|---|---|---|---|---|---|---|---|---|---|---|---|---|---|---|---|
| b2 | amm, asp, urs, berus, seo | - | - | - | - | - | - | - | - | - | - | - | - | - | - | - |
| b3 | mon, lat | - | - | - | - | - | - | - | - | - | - | - | - | - | - | - |
| b4 | amm, asp | - | 0 | 0 | 0 | 0 | 0 | - | - | - | - | - | - | - | - | - |
| b5 | urs, berus, seo | - | 0 | 0.044 | 0 | 0.03 | 0.03 | - | - | - | - | - | - | - | - | - |
| b6 | lat | - | - | - | - | - | - | 0 | 0 | 0 | 0 | 0 | 0 | 0 | 0 | 0 |
| b7 | mon | - | - | - | - | - | - | 0 | 0 | 0 | 0 | 0 | 0 | 0 | 0 | 0 |
| b8 | amm | - | 0 | 0 | 0 | 0 | 0 | - | - | - | - | - | 0 | 0 | 0 | 0 |
| b9 | asp | - | 0 | 0 | 0 | 0 | 0 | - | - | - | - | - | 0 | 0.05 | 0.039 | 0.042 |
| b10 | urs | - | 0 | 0 | 0 | 0 | 0 | 0.085 | 0 | 0 | 0 | 0 | - | - | - | - |
| b11 | berus, seo | - | 0 | 0 | 0.038 | 0.027 | 0.027 | 0 | 0.065 | 0 | 0 | 0 | - | - | - | - |
| b12 | latgad, lataru | - | - | - | - | 0 | 0 | 0 | 0 | 0 | 0 | 0 | 0 | 0 | 0 | 0 |
| b13 | latlat | - | - | - | - | 0 | 0 | 0 | 0 | 0 | 0 | 0 | 0 | 0 | 0 | 0 |
| b14 | monS | - | 0 | 0 | 0 | - | - | 0 | 0 | 0 | 0 | 0 | 0 | 0 | 0 | 0 |
| b15 | monN | - | 0 | 0 | 0 | - | - | 0 | 0 | 0 | 0 | 0 | 0 | 0 | 0.003 | 0 |
| b16 | aspasp, aspzin | - | 0 | 0 | 0 | 0 | 0 | 0 | - | - | - | - | 0 | 0.051 | 0.035 | 0.037 |
| b17 | aspfra, asphug | - | 0 | 0 | 0 | 0 | 0 | 0 | - | - | - | - | 0 | 0 | 0 | 0 |
| b18 | seo | - | 0.2 | 0.23 | 0.184 | 0.159 | 0.156 | 0 | 0 | 0 | 0 | 0 | 0 | - | - | - |
| b19 | berus | - | 0 | 0 | 0 | 0 | 0 | 0 | 0.012 | 0 | 0 | 0 | 0.076 | - | - | - |
| b20 | latgad | - | 0 | - | - | 0 | 0 | 0 | 0 | 0 | 0 | 0 | 0 | 0 | 0 | 0 |
| b21 | lataru | - | 0 | - | - | 0 | 0 | 0 | 0 | 0 | 0 | 0 | 0 | 0 | 0 | 0 |
| b22 | aspasp | - | 0 | 0 | 0 | 0 | 0 | 0 | - | - | 0.162 | 0.142 | 0 | 0 | 0 | 0 |
| b23 | aspzin | - | 0.07 | 0.07 | 0.058 | 0.05 | 0.049 | 0 | - | - | 0 | 0 | 0 | 0 | 0.099 | 0.09 |
| b24 | aspfra | - | 0 | 0 | 0 | 0 | 0 | 0 | 0.03 | 0.035 | - | - | 0 | 0 | 0 | 0 |
| b25 | asphug | - | 0 | 0 | 0 | 0 | 0 | 0 | 0 | 0 | - | - | 0 | 0 | 0 | 0 |
| b26 | seoW | - | 0.035 | 0.04 | 0.032 | 0.027 | 0.027 | 0 | 0 | 0 | 0 | 0 | 0 | 0 | - | - |
| b27 | seoE | - | 0 | 0 | 0 | 0 | 0 | 0 | 0 | 0 | 0 | 0 | 0 | 0 | - | - |

**Table S6. Niche overlap analyses between *V. latastei* and *V. seoanei*, including their Western and Eastern genomic lineages.** Niche overlap is measured by the Sørenson index (K) and the Overlap Index (OI). As well, the niche hypervolume of each species' niche of the comparison is shown, the intersection and union among them, and the unique components of each compared species (Uniq. spX relative to spY). Vlat stands for *V. latastei*, Vseo for *V. seoanei*, W or E for Western or Eastern lineage of both species. Some comparisons (pure) excluded admixed populations from other species.

| sp1 | sp2 | Vol. sp1 | Vol. sp2 | Intersection | Union | Uniq. sp1 rel. to sp2 | Uniq. sp2 rel. to sp1 | K | *p* | $K_{max}$ | OI | *p* |
|---|---|---|---|---|---|---|---|---|---|---|---|---|
| **Vlat (all)** | Vseo (all) | 2435322.1 | 1336650.6 | 410952.5 | 3361020.2 | 2024369.5 | 925698.1 | 0.215 | 0.01 | 0.696 | 0.309 | 0.01 |
| **Vlat (pure)** | Vseo (pure) | 2080461.9 | 1267075.6 | 235355.1 | 3112182.4 | 1845106.8 | 1031720.5 | 0.146 | 0.01 | 0.747 | 0.196 | 0.01 |
| **VlatE** | VlatW | 656106.52 | 1868500.26 | 78470.62 | 2446136.16 | 577635.89 | 1790029.64 | 0.062 | 0.01 | 0.521 | 0.119 | 0.01 |
| **VseoW** | VseoE | 1225437 | 322170.2 | 204707.1 | 1342900 | 1020729.9 | 117463.1 | 0.255 | 0.01 | 0.413 | 0.617 | 0.01 |
| **VlatE** | VseoE | 657782.28 | 321384.46 | 34745.53 | 944421.21 | 623036.76 | 286638.93 | 0.072 | 0.01 | 0.661 | 0.109 | 0.01 |
| **VlatW (all)** | VseoW (all) | 1893085.7 | 1233109.9 | 388438.7 | 2737757 | 1504647 | 844671.2 | 0.252 | 0.01 | 0.791 | 0.318 | 0.01 |
| **VlatW (pure)** | Vseo W (pure) | 1468990.1 | 1178598.9 | 107860.7 | 2539728.3 | 1361129.4 | 1070738.2 | 0.083 | 0.01 | 0.89 | 0.093 | 0.01 |

**Table S7. Factor loadings of ecological PCA with 19 WorldClim v2.1 (Fick & R.J. Hijmans, 2017) variables for the three main Principal Components (88.04% of explained variance).** Main factor loadings in bold.

| Variable | Meaning | PC1 | PC2 | PC3 |
|---|---|---|---|---|
| **bio 1** | Annual Mean Temperature | 0.697 | 0.595 | -0.305 |
| bio 2 | Mean Diurnal Range | 0.723 | -0.443 | 0.264 |
| bio 3 | Isothermality (BIO2/BIO7) | 0.171 | 0.524 | -0.255 |
| bio 4 | Temperature Seasonality | 0.535 | **-0.730** | 0.355 |
| bio 5 | Max Temperature of Warmest Month | **0.946** | -0.034 | 0.094 |
| bio 6 | Min Temperature of Coldest Month | 0.243 | **0.872** | -0.394 |
| bio 7 | Temperature Annual Range (BIO5-BIO6) | 0.650 | -0.630 | 0.352 |
| bio 8 | Mean Temperature of Wettest Quarter | 0.442 | 0.068 | **-0.736** |
| bio 9 | Mean Temperature of Driest Quarter | 0.536 | 0.434 | 0.207 |
| bio 10 | Mean Temperature of Warmest Quarter | 0.901 | 0.242 | -0.126 |
| bio 11 | Mean Temperature of Coldest Quarter | 0.438 | 0.791 | -0.392 |
| bio 12 | Annual Precipitation | -0.853 | 0.417 | 0.247 |
| bio 13 | Precipitation of Wettest Month | -0.681 | 0.627 | 0.296 |
| bio 14 | Precipitation of Driest Month | -0.896 | -0.325 | -0.147 |
| bio 15 | Precipitation Seasonality | 0.462 | **0.787** | 0.303 |
| bio 16 | Precipitation of Wettest Quarter | -0.683 | 0.627 | 0.317 |
| bio 17 | Precipitation of Driest Quarter | **-0.924** | -0.258 | -0.108 |
| bio 18 | Precipitation of Warmest Quarter | **-0.919** | -0.235 | -0.163 |
| bio 19 | Precipitation of Coldest Quarter | -0.632 | 0.659 | 0.376 |

**Table S8. Toxin-encoding genes identified in *V. latastei* genome rVipLat1.** Genes are ordered by chromosome and sorted out in toxin families. The transcript name is the specific isoform whose CDS has been used in selection analyses. Peptide hits are the number of unique peptide sequences from the *V. latastei* proteome which have been found in that isoform. Upreg. Column shows whether that gene has been found to be significantly upregulated in the venom gland. Microsynteny of the primary and secondary toxin families is displayed in Fig. S33. Three non-venomous genes related to toxin-encoding counterparts are shown at the bottom.

| Family | Gene | Transcript | Peptide hits | Upreg. | Chrom | Strand | Description |
|---|---|---|---|---|---|---|---|
| **CRiSP** | g_96919 | long_reads1.PB.4209.307 | 10 | YES | 1 | + | Cysteine Rich Venom Protein-1 |
| **5'nucleotidase** | g_97788 | long_reads1.PB.5295.83 | 1 | | 1 | + | 5'-nucleotidase |
| **PDE** | g_98211 | long_reads1.PB.5717.6 | 3 | | 1 | + | Venom phosphodiesterase |
| **LAAO** | g_48392 | long_reads1.PB.13771.87 | 20 | | 2 | - | L-amino acid oxidase LAAO-II |
| **LAAO** | g_48393 | anno4.cro_tig_rna-XM_039361015.1_R1 | 3 | | 2 | - | L-amino acid oxidase |
| **Kunitz** | g_162 | long_reads1.PB.14656.7 | 0 | YES | 3 | - | Serine protease inhibitor 3 |
| **Kunitz** | g_5167 | anno4.cro_tig_rna-XM_039342386.1_R0 | 1 | | 3 | + | Kunitz peptide |
| **Kunitz** | g_14506 | anno4.pan_gut_rna-XM_034405137.1_R7 | 7 | | 3 | + | Kunitz-type serine protease inhibitor |
| **Kunitz** | g_14507 | anno4.pan_gut_rna-XM_034405137.1_R5 | 6 | | 3 | + | Kunitz-type serine protease inhibitor |
| **Kunitz** | g_14509 | anno4.pan_gut_rna-XM_034405137.1_R4 | 7 | YES | 3 | + | Kunitz-type serine protease inhibitor |
| **vNGF** | g_1242 | long_reads1.PB.15420.184 | 1 | YES | 3 | + | Venom nerve growth factor |
| **vNGF** | g_14670 | anno4.cro_tig_rna-XM_039356926.1_R1 | 1 | | 3 | + | Venom nerve growth factor |
| **svMPi** | g_82078 | long_reads1.PB.21558.84 | 1 | YES | 5 | - | MPi-3 |
| **svMPi** | g_82081 | long_reads1.PB.21557.998 | 1 | YES | 5 | + | endogenous tripeptide metalloproteinase inhibitor precursor |
| **CTL** | g_71781 | long_reads1.PB.23578.6 | 0 | YES | 6 | + | Ancestral single-chain lectin toxin |

| | | | | | | | |
|---|---|---|---|---|---|---|---|
| **CTL** | g_71812 | long_reads1.PB.23585.6 | 5 | YES | 6 | + | Snaclec subunit A and B. Transcript example of subunit B: long_reads1.PB.23585.231 |
| **CTL** | g_77666 | long_reads1.PB.23587.74 | 1 | YES | 6 | - | Snaclec subunit B |
| **CTL** | g_71813 | long_reads1.PB.23587.53 | 6 | YES | 6 | - | Snaclec subunit A and B. Transcript example of subunit B: long_reads1.PB.23587.253 |
| **CTL** | g_71814 | long_reads1.PB.23587.423 | 3 | YES | 6 | - | Snaclec subunit A |
| **CTL** | g_71817 | long_reads1.PB.23585.597 | 0 | YES | 6 | + | Snaclec subunit B |
| **PLB** | g_72774 | long_reads1.PB.24340.6 | 1 | | 6 | + | Phospholipase B |
| **svMP** | g_134876 | long_reads1.PB.27234.89 | 13 | YES | 8 | + | Metalloproteinase MPIII |
| **svSP** | g_130459 | long_reads1.PB.28174.636 | 11 | YES | 9 | - | Serine protease |
| **svVEGF** | g_131055 | anno1.g48961.t1 | 7 | YES | 9 | + | vammin-1' |
| **XaaPro** | g_38002 | long_reads1.PB.30060.88 | 6 | | 12 | + | xaa-Pro aminopeptidase 2 |
| **CYS** | g_79330 | anno4.pan_gut_rna-XM_034427211.1_R7 | 1 | | 16 | + | Cystatin 1 |
| **PLA** | g_133624 | long_reads1.PB.32683.3 | 13 | YES | 17 | - | Ammodytin I2(A) |
| **PLA** | g_133627 | long_reads1.PB.32686.117 | 9 | YES | 17 | + | Ammodytin I1(D) |
| **LAAO-like** | g_48408 | anno4.cro_tig_rna-XM_039361019.1_R0 | | | 2 | - | L-amino-acid oxidase-like. Non venomous |
| **LAAO-like** | g_50929 | anno2.g17375.t1 | | | 2 | + | L-amino acid oxidase-like. Non venomous |
| **PLA_gIIE** | g_134145 | anno4.pan_gut_rna-XM_034424295.1_R9 | | | 17 | + | Ancestral non-venomous $PLA_2$. Non venomous |

**Table S9. Toxin-encoding genes identified in *V. aspis* genome rVipAsp1.** Genes are ordered by chromosome and sorted out in toxin families. The transcript name is the specific isoform whose CDS has been used in selection analyses. Peptide hits are the number of unique peptide sequences from the *V. aspis* proteome which have been found in that isoform. Microsynteny of the primary and secondary toxin families is displayed in Fig. S33. Four non-venomous genes related to toxin-encoding counterparts are shown at the bottom.

| Family | Gene | Transcript | Peptide hits | Chrom. | Strand | Description |
|---|---|---|---|---|---|---|
| **CRiSP** | g_33918 | long_reads1.PB.695.69 | 12 | 1 | - | Cysteine-rich venom protein |
| **CRiSP** | g_28182 | long_reads1.PB.695.4 | 14 | 1 | - | Cysteine-rich venom protein |
| **5’nucleotidase** | g_28618 | long_reads1.PB.1209.71 | 22 | 1 | + | 5'-nucleotidase |
| **5’nucleotidase** | g_28619 | long_reads1.PB.1209.244 | 14 | 1 | + | 5'-nucleotidase |
| **PDE** | g_29051 | long_reads1.PB.1629.1 | 15 | 1 | + | Phosphodiesterase |
| **vQC** | g_29579 | long_reads1.PB.2193.13 | 11 | 1 | - | Glutaminyl-peptide cyclotransferase |
| **svVEGF** | g_29661 | long_reads1.PB.2291.57 | 4 | 1 | + | Vascular endothelial growth factor A |
| **LAAO** | g_6353 | long_reads1.PB.7791.79 | 33 | 2 | - | L-amino acid oxidase |
| **LAAO** | g_3579 | anno4.cro_tig_rna-XM_039361015.1_R2 | 2 | 2 | - | L-amino acid oxidase |
| **LAAO** | g_6356 | anno4.cro_tig_rna-XM_039361013.1_R4 | 3 | 2 | - | L-amino acid oxidase |
| **LAAO** | g_3580 | anno4.cro_tig_rna-XM_039361019.1_R3 | 5 | 2 | - | L-amino acid oxidase |
| **PPi** | g_13950 | long_reads1.PB.15243.2 | 1 | 5 | - | Peptidyl-prolyl cis-trans isomerase C |
| **Kunitz** | g_70866 | anno4.cro_tig_rna-XM_039342380.1_R0 | 0 | 7 | - | Serine protease inhibitor 3 |
| **Kunitz** | g_74066 | anno4.cro_tig_rna-XM_039342386.1_R1 | 0 | 7 | + | Kunitz/BPTI-like toxin |
| **Kunitz** | g_74067 | anno4.pan_gut_rna-XM_034405137.1_R4 | 4 | 7 | + | Kunitz-type serine protease inhibitor |
| **Kunitz** | g_74068 | anno4.pan_gut_rna-XM_034405137.1_R5 | 5 | 7 | + | Kunitz-type serine protease inhibitor |

| | | | | | | |
|---|---|---|---|---|---|---|
| **Kunitz** | g_74069 | anno4.pan_gut_rna-XM_034405137.1_R6 | 5 | 7 | + | Kunitz-type serine protease inhibitor |
| **Kunitz** | g_74071 | long_reads1.PB.18914.159 | 3 | 7 | + | Kunitz-type serine protease inhibitor |
| **Kunitz** | g_74072 | anno4.pan_gut_rna-XM_034405137.1_R7 | 1 | 7 | + | Kunitz-type serine protease inhibitor |
| **vNGF** | g_71982 | long_reads1.PB.19627.61 | 13 | 7 | + | Venom nerve growth factor |
| **RLAP** | g_72151 | long_reads1.PB.19854.45 | 4 | 7 | + | Renin-like aspartic protease |
| **Prosaposin** | g_56972 | long_reads1.PB.21396.97 | 4 | 8 | - | Prosaposin isoform X2 |
| **svMPi** | g_58598 | long_reads1.PB.22989.178 | 22 | 8 | - | Endogenous tripeptide metalloproteinase inhibitor precursor |
| **svMPi** | g_59515 | long_reads1.PB.22987.159 | 27 | 8 | + | Endogenous tripeptide metalloproteinase inhibitor precursor |
| **CTL** | g_78881 | long_reads1.PB.23130.9 | 7 | 9 | + | Ancestral single-chain lectin toxin |
| **CTL** | g_78909 | long_reads1.PB.23137.139 | 22 | 9 | + | Snaclec subunit A and B. transcript example of subunit B: long_reads1.PB.23137.121 |
| **CTL** | g_78912 | long_reads1.PB.23138.95 | 17 | 9 | - | Snaclec subunit A |
| **CTL** | g_78913 | long_reads1.PB.23138.61 | 2 | 9 | - | Snaclec subunit A |
| **CTL** | g_78914 | long_reads1.PB.23138.490 | 6 | 9 | - | Snaclec subunit A |
| **PLB** | g_79927 | long_reads1.PB.23926.4 | 9 | 9 | + | Phospholipase B |
| **hyaluronidase** | g_80652 | long_reads1.PB.24587.28 | 3 | 9 | + | Hyaluronidase |
| **aminopeptidase** | g_40343 | long_reads1.PB.25597.11 | 18 | 10 | + | Glutamyl aminopeptidase |
| **svMP** | g_9891 | long_reads1.PB.27853.3060 | 41 | 11 | - | Metalloproteinase of class P-III |
| **CP** | g_67109 | long_reads1.PB.28470.8 | 2 | 12 | + | Cysteine proteinase 2-like |
| **svSP** | g_67321 | long_reads1.PB.28631.537 | 21 | 12 | - | Serine proteinase SP-8 |
| **svVEGF** | g_67922 | long_reads1.PB.28961.109 | 15 | 12 | + | vammin-1' |
| **DPP** | g_84414 | long_reads1.PB.29453.2 | 3 | 13 | - | dipeptidase 2-like |
| **XaaPro** | g_75857 | long_reads1.PB.29901.62 | 26 | 14 | - | xaa-Pro aminopeptidase 2 |
| **PPi** | g_69564 | long_reads1.PB.30554.65 | 3 | 15 | + | Peptidyl-prolyl cis-trans isomerase B |

| | | | | | | |
|---|---|---|---|---|---|---|
| **PLA** | g_11106 | long_reads1.PB.32751.122 | 28 | 20 | - | ammodytin I1 |
| **neuroPLA** | g_11113 | anno4.cro_tig_rna-XM_039367476.1_R2 | 4 | 20 | + | Vaspin basic subunit variant. Transcript example of acidic subunit: anno4.cro_tig_rna-XM_039367475.1_R0 |
| **CRISP-like** | g_33919 | anno4.cro_tig_rna-XM_039329609.1_R0 | 0 | 1 | - | serotriflin. Non venomous. |
| **LAAO-like** | g_6374 | anno4.naj_naj_rna-gnl_WGS:SOZL_mRNA16818_R5 | 0 | 2 | - | L-amino acid oxidase-like. Non venomous. |
| **LAAO-like** | g_5192 | anno2.g34869.t1 | 0 | 2 | + | L-amino acid oxidase-like. Non venomous. |
| **svMP-ancestral** | g_9895 | long_reads1.PB.27853.3260 | 1 | 11 | - | ADAM28. Non venomous. |

**Table S10. Toxin-encoding genes identified in *V. seoanei* genome rVipSeo1.** Genes are ordered by chromosome and sorted out in toxin families. The transcript name is the specific isoform whose CDS has been used in selection analyses. Peptide hits are the number of unique peptide sequences from the *V. seoanei* proteome which have been found in that isoform. Microsynteny of the primary and secondary toxin families is displayed in Fig. S33. Four non-venomous genes related to toxin-encoding counterparts are shown at the bottom.

| Family | Gene | Transcript | Peptide hits | Chrom. | Strand | Description |
|---|---|---|---|---|---|---|
| **CRiSP** | g_41326 | long_reads1.PB.4352.28 | 17 | 1 | + | Cysteine-rich venom protein |
| **LAAO** | g_877 | long_reads1.PB.8947.374 | 30 | 2 | + | L-amino acid oxidase LAAO-II |
| **LAAO** | g_875 | anno1.g991.t1 | 4 | 2 | + | L-amino acid oxidase |
| **LAAO** | g_7610 | anno4.cro_tig_rna-XM_039361015.1_R1 | 6 | 2 | + | L-amino acid oxidase |
| **Kunitz** | g_95072 | anno4.pan_gut_rna-XM_034424188.1_R7 | 1 | 3 | + | Kunitz-type serine protease inhibitor |
| **Kunitz** | g_114630 | long_reads1.PB.14544.37 | 3 | 3 | + | Kunitz-type serine protease inhibitor |

| | | | | | | |
|---|---|---|---|---|---|---|
| **vNGF** | g_96318 | long_reads1.PB.15297.63 | 8 | 3 | + | Venom nerve growth factor 1 |
| **CTL** | g_136710 | long_reads1.PB.26260.83 | 3 | 6 | + | Snaclec subunit A |
| **CTL** | g_136709 | long_reads1.PB.26261.190 | 13 | 6 | - | Snaclec subunit A and B. Transcript example of subunit B: long_reads1.PB.26261.539 |
| **CTL** | g_136713 | long_reads1.PB.26260.403 | 9 | 6 | + | Snaclec subunit A and B. Transcript example of subunit B: long_reads1.PB.26260.348 |
| **CTL** | g_136734 | long_reads1.PB.26269.19 | 0 | 6 | - | Ancestral single-chain lectin toxin |
| **svMP** | g_143356 | long_reads1.PB.28146.617 | 22 | 8 | + | Zinc Metalloproteinase of class P-III |
| **svSP** | g_26734 | long_reads1.PB.29112.1266 | 23 | 9 | - | Snake Venom Serine Protease |
| **svVEGF** | g_27370 | long_reads1.PB.29435.74 | 24 | 9 | + | Vammin-1 |
| **PLA** | g_117442 | long_reads1.PB.33347.218 | 24 | 17 | - | Ammodytin I1 |
| **PLA** | g_117449 | long_reads1.PB.33345.60 | 21 | 17 | + | Ammodytin I2 |
| **CRiSP-like** | g_66533 | anno4.cro_tig_rna-XM_039329609.1_R0 | 0 | 1 | + | Serotriflin. Non venomous. |
| **LAAO-like** | g_10173 | anno4.naj_naj_rna-gnl_WGS:SOZL_mRNA16818_R4 | 0 | 2 | + | L-amino-acid oxidase-like. Non venomous. |
| **svMP-ancestral** | g_145534 | anno4.cro_tig_rna-XM_039354198.1_R0 | 1 | 8 | + | ADAM28. Non venomous. |
| **PLA_gIIE** | g_117452 | long_reads1.PB.33346.10 | 0 | 17 | + | Phospholipase A2-like group IIE. Non venomous. |

**Table S11. Selection tests on toxin-encoding genes per family.** $d_N/d_S$ tests were performed with BUSTED and FEL implemented on Datamonkey. BUSTED looks for codons with episodic diversifying selection (significant values in bold), whereas FEL tests for codons with either pervasive diversifying or purifying selection, with a fixed *p-value* in 0.1. Only toxin families with at least 3 copies in at least 2 species (spp.) were tested.

| | | | | BUSTED | | FEL | | |
|---|---|---|---|---|---|---|---|---|
| **Family** | **spp.** | **copies** | **codons** | ***p-value*** | **codons** | **p-*value*** | **codons diversifying sel.** | **codons purifying sel.** |
| SVMP | 3 | 3 | 614 | 0.018 | 4 | 0.1 | 0 | 27 |
| $PLA_2$ | 3 | 6 | 143 | <0.001 | 20 | 0.1 | 3 | 6 |
| snaclec | 3 | 12 | 162 | <0.001 | 19 | 0.1 | 15 | 19 |
| CRISP | 3 | 4 | 260 | <0.001 | 51 | 0.1 | 1 | 3 |
| LAAO | 3 | 9 | 549 | <0.001 | 56 | 0.1 | 8 | 33 |
| SVVEGF | 3 | 3 | 131 | 0.5 | 0 | 0.1 | 0 | 2 |
| SVSP | 3 | 3 | 259 | 0.0042 | 10 | 0.1 | 0 | 5 |
| SVNGF | 3 | 4 | 260 | 0.38 | 3 | 0.1 | 0 | 1 |
| Kunitz | 3 | 12 | 87 | <0.001 | 11 | 0.1 | 2 | 8 |
| SVMPi | 2 | 4 | 253 | <0.001 | 7 | 0.1 | 0 | 8 |
| 5'-nucleotidase | 2 | 3 | 592 | 0.5 | 0 | 0.1 | 0 | 17 |

**Table S12. Whole-genome datasets used in this study.** The table shows whether the dataset have high- or low-coverage, the number of individuals (n), species (spp.), SNPs (if not all sites) and the analyses that were performed with them.

| Coverage | Dataset Name | n | spp. | SNPs | Analyses | Comments |
|---|---|---|---|---|---|---|
| **High** | *high_coverage* | 8 | 7 | all sites | PSMC, ROHs | |
| **High** | *high_coverage+out* | 8 | 8 | all sites | Twisst | With *Crotalus* as outgroup |
| **Low** | *all_sites* | 95 | 7 | all sites | Heterozygosity | |
| **Low** | *westmed* | 90 | 4 | 1,588,578 | PCA, Admixture | 0.5-kbp thinning |
| **Low** | *latastei-monticola* | 45 | 2 | 1,308,394 | PCA | 0.5-kbp thinning, no hybrids |
| **Low** | *latastei* | 37 | 1 | 1,309,324 | PCA, IBD | 0.5-kbp thinning, no hybrids |
| **Low** | *monticola* | 8 | 1 | 326,645 | PCA | 0.5-kbp thinning |
| **Low** | *aspis* | 13 | 1 | 1,152,108 | PCA, IBD | 0.5-kbp thinning, no hybrids |
| **Low** | *seoanei* | 30 | 1 | 1,504,794 | PCA | 0.5-kbp thinning |
| **Low** | *linked_SNPs* | 86 | 8 | 12,645,411 | Dsuite | No LD-pruned, no hybrids, no intraspecific admixture, with *Crotalus* as outgroup |
| **Low** | *snapp_species* | 8 | 8 | 1,225,190 | SNAPP | 0.5-kbp thinning, with *Crotalus* as outgroup, 0% missingness |
| **Low** | *snapp_subspecies* | 15 | 7 | 126,083 | SNAPP | 10-kbp thinning, 0% missingness |
| **Low** | *treemix_subspecies* | 15 | 7 | 1,095,590 | TreeMix | 0.5-kbp thinning, 0% missingness |

**Table S13. Toxin families in *Vipera*, with the number of gene copies found for each one, the relative expression (*sensu* Giribaldi et al., 2020), and the normalized ubiquity of diversifying and purifying selection, corrected by number of copies and length of the alignment.** These values have been used in linear regression shown in Fig. 6D.

| Family | Copies | Expression (%) | Diversification | Purification |
|---|---|---|---|---|
| SVMP | 3 | 28.31 | 0 | 1 |
| PLA | 6 | 15.45 | 0.45 | 0.48 |
| snaclec | 12 | 12.28 | 1 | 0.67 |
| CRISP | 4 | 0.79 | 0.12 | 0.20 |
| LAAO | 9 | 1.07 | 0.21 | 0.46 |
| SVVEGF | 3 | 3.19 | 0 | 0.35 |
| SVSP | 3 | 13.31 | 0 | 0.44 |
| SVNGF | 4 | 1.89 | 0 | 0.07 |
| Kunitz | 12 | 9.87 | 0.25 | 0.52 |
| SVMPi | 4 | 13.2 | 0 | 0.54 |
| 5’-nucleotidase | 3 | 0.25 | 0 | 0.65 |

**Table S14. Samples used for phylogenetic inference and the lineage to which they belong.** All these samples were used to build the *snapp_subspecies* and *treemix_subspecies* datasets, as well as for extracting their mitogenomes to infer the mitochondrial phylogeny, but only the first seven samples were included in *snapp_species* dataset (see Table S11).

| Sample | Species | Subespecies/lineage | *snapp_species* |
|---|---|---|---|
| **ATP14** | *V. ammodytes* | Northwestern | Y |
| **CNMOL1** | *V. aspis* | *zinnikeri* | Y |
| **SPM002443** | *V. berus* | *berus* | Y |
| **CNPOR1** | *V. latastei* | *latastei* | Y |
| **FM19VM052** | *V. monticola* | *monticola* | Y |
| **CNSOC1** | *V. seoanei* | East | Y |
| **CN13667** | *V. ursinii* | *rakosiensis* | Y |
| **MALZ004** | *V. aspis* | *hugyi* | N |
| **FM22VL233** | *V. latastei* | *gaditana* | N |
| **EMO4112020** | *V. aspis* | *francisciredi* | N |
| **BEV11035** | *V. aspis* | *aspis* | N |
| **CN20038** | *V. latastei* | *arundana* | N |
| **MNCN50498** | *V. monticola* | *saintgironsi* | N |
| **FM19VM061** | *V. monticola* | *atlantica* | N |
| **SPM003076R** | *V. seoanei* | West | N |

**Table S15. Partitions for the mitochondrial protein-coding gene alignment of 10,695 bp arranged by PartitionFinder.** This partition scheme and its respective nucleotide alignment were used to infer the mitochondrial phylogeny of the studied *Vipera* spp., depicted in Figs. 2, S7.

| Partition | Best Model | Sites | Genes and positions |
|---|---|---|---|
| 1 | TVM+I | 676 | ATP6_1st, ND4L_1st, ND4_1, ND3_1st |
| 2 | GTR+I+G | 1617 | ND2_2nd, ND4L_2nd, ND4_2nd, ND5_2nd ND3_2nd, ATP6_2nd |
| 3 | GTR+I+G | 465 | ATP6_3rd, ND4_3rd |
| 4 | TRN+I | 110 | ATP8_1st, ATP8_2nd |
| 5 | TIM+I+G | 286 | ATP8_3rd, COX2_3rd |
| 6 | TRNEF+G | 534 | COX1_1st |
| 7 | K81UF+I | 534 | COX1_2nd |
| 8 | GTR+I+G | 909 | ND3_3rd, COX1_3rd, COX3_3rd |
| 9 | TRN+I | 231 | COX2_1st |
| 10 | TIM+I | 552 | COX2_2nd, ND1_2nd |
| 11 | GTR+I+G | 954 | COX3_1st, CTYB_1st, ND1_1st |
| 12 | TIM+I | 633 | CTYB_2nd, COX3_2nd |
| 13 | TIM+I+G | 969 | CTYB_3rd, ND5_3rd |
| 14 | TIM+I+G | 762 | ND1_3rd, ND4L_3rd, ND2_3rd |
| 15 | GTR+G | 941 | ND2_1st, ND5_1st |
| 16 | TRN+I+G | 174 | ND6_1st |
| 17 | HKY+G | 174 | ND6_2nd |
| 18 | TIM+G | 174 | ND6_3rd |

**Table S16. Cross Validation (CV) average and standard deviation (SD) values for Admixture analysis.** A likelihood plateau is reached at K=3, although there are four species in the dataset. The highest likelihood is reached at K=5, although K=7 has the highest biological significance.

| K | CV Average | CV SD |
|---|---|---|
| 1 | 0.4371 | 0.0002 |
| 2 | 0.3231 | 0.0211 |
| 3 | 0.2484 | 0.0002 |
| 4 | 0.2453 | 0.0063 |
| 5 | 0.2439 | 0.0011 |
| 6 | 0.2463 | 0.0019 |
| 7 | 0.2599 | 0.0092 |
| 8 | 0.2623 | 0.0047 |
| 9 | 0.2709 | 0.0061 |
| 10 | 0.2813 | 0.0048 |
| 11 | 0.2948 | 0.0109 |
| 12 | 0.3038 | 0.0072 |
